# Genome-wide computational prediction of miRNAs encoded by influenza A virus (H3N2) predicts target genes involved in pulmonary and antiviral innate immunity

**DOI:** 10.64898/2026.05.18.725090

**Authors:** Maria Ajmal Siddiqi, Harsh Kumar, Mohit Mazumder

## Abstract

Influenza A virus (IAV) causes significant morbidity and mortality worldwide. Understanding how viral RNAs may regulate host genes through microRNA-like mechanisms can clarify pathogenesis and reveal therapeutic targets. In this study, we screened all eight IAV H3N2 RNA segments (PB2, PB1, PA, HA, NP, NA, M, and NS) using an ab initio computational pipeline; five segments (PB2, PB1, PA, HA, and M) met the VMir scoring threshold for further analysis, while NP, NA, and NS were excluded due to low pre-miRNA scores. Mature miRNAs were identified using MatureBayes, and target genes in the human genome were predicted with the miRDB server. From these targets, we selected two genes per qualifying segment (10 genes total) based on their functional relevance to influenza infection and supporting literature; all selected genes are unique to their respective segment. We identified 10 segment-specific target genes (IFNL1, DDX60, SAMHD1, MAVS, IRF4, BIRC2, AGO1, MAP3K1, NOD1, and TNFAIP1) and one common target across all five analyzed segments (CADM2). Gene Ontology and pathway analyses showed enrichment in interferon signaling, RIG-I-like receptor pathways, antiviral restriction, RNA interference, and inflammatory responses. Literature supports roles for these genes in pulmonary and antiviral innate immunity. Our findings provide a basis for experimental validation and may help the research community better understand influenza virus pathogenesis and identify novel therapeutic candidates.

**GRAPHICAL ABSTRACT:** 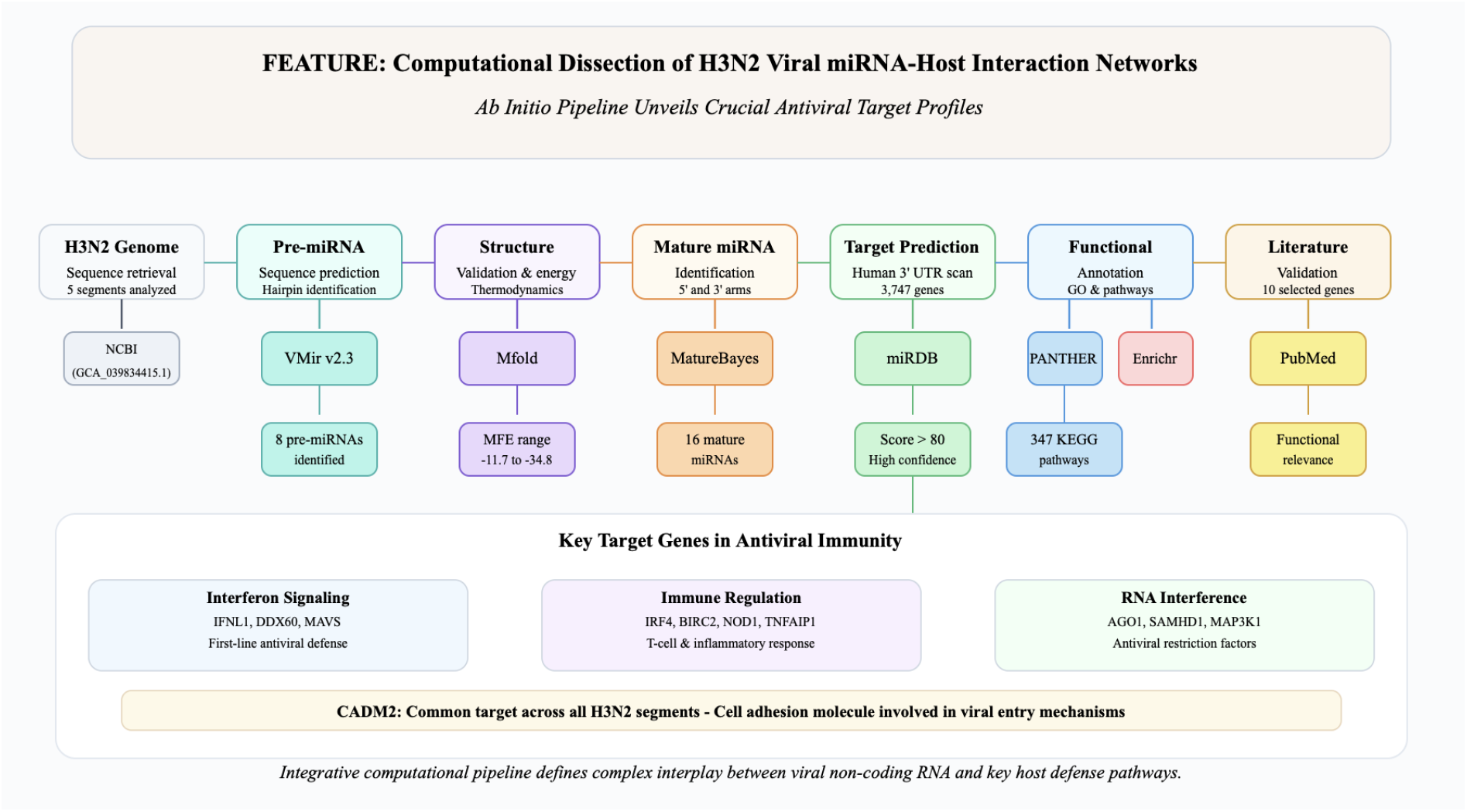

## INTRODUCTION

Influenza is an acute respiratory infection caused by influenza viruses and remains a major global public health concern. It is responsible for an estimated 3-5 million cases of severe illness and around 290,000 to 650,000 respiratory deaths worldwide each year. Among the four influenza virus subtypes, IAV subtype H3N2 is one of the main circulating seasonal strains and often dominates influenza outbreaks. This subtype significantly contributes to the global disease burden and is frequently linked to increased morbidity, especially among older adults [1,2]. The genome of IAV consists of eight segments of single-stranded, negative-sense RNA that encode the polymerase subunits (PB2, PB1, PA), the hemagglutinin subtype H3 (HA), the neuraminidase subtype N2 (NA), the nucleoprotein (NP), the matrix (M), and the non-structural (NS) proteins. Of these, segments HA and NA encode surface glycoproteins responsible for viral entry and release, while the other segments encode internal proteins essential for viral replication, assembly, and interaction with the host [2,3]. Clinical manifestations of H3N2 infection typically present as acute respiratory distress characterized by pyrexia, cough, pharyngitis, rhinorrhea, myalgia, and significant fatigue; advanced pathogenesis may progress to severe dyspnea [1]. The definitive diagnosis of the H3N2 subtype is established using laboratory methods such as real-time reverse transcription-polymerase chain reaction (RT-PCR), rapid antigen assays, or gold-standard viral culture [2]. Elucidating the regulatory influence of H3N2 on the host transcriptome is essential for deciphering viral pathogenesis and identifying novel therapeutic interventions.

MicroRNAs (miRNAs) are small (∼22 nt) non-coding RNAs that regulate gene expression after transcription by binding to complementary sites on mRNA. This binding can either inhibit translation or cleave mRNA, depending on the level of complementarity [4,5,6]. MiRNAs have been identified in plants, animals, and fungi [7,8]. Additionally, viral-encoded miRNAs have been reported in several virus families and can influence host defense, cell differentiation, apoptosis, and cell proliferation. The involvement of small viral RNAs in respiratory infections and immune regulation has been demonstrated [5,9]. However, identifying miRNAs experimentally often requires expression analysis in specific cell types and involves time-consuming cloning techniques. To understand disease causes more quickly, computational prediction of miRNAs from genome analysis can provide early insights, helping to track their role in disease development during outbreaks [7,8].

Computational prediction of microRNAs generally employs two main strategies: homology-based and ab initio approaches. Homology-based methods rely on evolutionary conservation and are therefore limited in their ability to identify new microRNAs. Conversely, ab initio approaches analyze genomic sequences for characteristic hairpin secondary structures, enabling the discovery of previously uncharacterized pre-miRNAs and offering broader discovery potential [7,10]. To date, RNA virus-encoded miRNAs have been predicted in several viruses [11]. Therefore, in this study, the genome of the influenza A virus (IAV) subtype H3N2, comprising eight segments (PB2, PB1, PA, HA, NP, NA, M, and NS), was analyzed to predict mature viral miRNAs. The predicted miRNAs were then screened for potential target genes in the human genome, followed by gene ontology and pathway enrichment analyses to assess their functional significance.

## MATERIALS AND METHODS

### Data Retrieval

H3N2 segment sequences (PB2, PB1, PA, HA, NP, NA, M, and NS) were obtained from the NCBI genome database (https://www.ncbi.nlm.nih.gov/genome/) using the accession number GCA_039834415.1. The segments consist of single-stranded, negative-sense RNA with a linear structure.

### Potential pre-miRNAs & mature miRNAs prediction

An ab initio-based pre-miRNA prediction software package, Vmir (v2.3), was used to identify IAV H3N2 pre-miRNAs. The VMir package includes two modules: the VMir analyzer, which predicts pre-miRNAs, and the VMir viewer, which displays them [12]. The analysis was performed using default parameters (window count: 500, conformation: linear, orientation: both) in the VMir analyzer. Additionally, filtering parameters (min. hairpin size: 70, min. score: ≥150, and min. window count: ≥35) were applied in the VMir viewer to select top-scoring pre-miRNAs, as described previously in the literature [13,14]. Secondary structures (Supplementary Figure S1) and minimum free energy (MFE) of pre-miRNAs were obtained using the web server Mfold (http://unafold.rna.albany.edu/?q=mfold) [15]. Mature miRNAs were predicted from pre-miRNAs using the Mature Bayes web server (http://mirna.imbb.forth.gr/MatureBayes.html), which employs a Naïve Bayes classifier and sequence/structure information to predict mature miRNA positions on the 5′ and 3′ arms [16].

### Target gene prediction and functional enrichment analysis

Predicted mature miRNA sequences served as input for the miRDB custom prediction module (http://mirdb.org/) to identify human target genes by analyzing the 3′ UTRs (untranslated regions) of the human genome for potential hybridization [17]. High-scoring targets (e.g., score >80) were kept. Gene Ontology (GO) terms for biological processes, molecular functions, cellular components, and pathway enrichment were retrieved using PANTHER (http://www.pantherdb.org) and Enrichr (https://amp.pharm.mssm.edu/Enrichr/) with NCBI Gene IDs [18,19]. A systematic literature review was performed to confirm the involvement of selected target genes in influenza pathogenesis. From the complete list of predicted targets for each segment, two genes per segment were selected based on (i) their functional relevance to influenza infection or antiviral immunity, (ii) supporting literature, and (iii) participation in host-virus pathways. To ensure segment-specific interpretation, we confirmed that each selected gene appeared only in its target segment’s list. Additionally, a common target was identified across all segments.

### Statistical Analysis and Validation

Gene Ontology (GO) enrichment analysis was conducted using PANTHER with Fisher’s exact test. Pathway enrichment analysis was performed with Enrichr, applying adjusted p-values calculated through the Benjamini-Hochberg false discovery rate (FDR) correction. Pathways with adjusted p-values less than 0.001 were regarded as significantly enriched. Target gene predictions with miRDB scores of 80 or higher were kept, based on previous validation studies showing more than 95% reliability at this threshold (Wong & Wang, 2015).

## RESULTS

VMir analysis was conducted on H3N2 segment sequences. Segments 5 (NP), 6 (NA), and 8 (NS) produced low VMir scores (130.7, 143.5, and 124, respectively) and were excluded; the analysis proceeded with the remaining five segments (PB2, PB1, PA, HA, M). After applying filtering parameters in the VMir viewer, eight high-scoring pre-miRNAs from these segments were retained. Two pre-miRNAs were located on the direct strand, while six were on the reverse strand. Additionally, all eight pre-miRNAs ranged from 73 to 122 nt in length, with VMir scores of 158.3-202.8. The sequence, rank, score, length, and orientation are listed in Table 1.

**Table 1:**
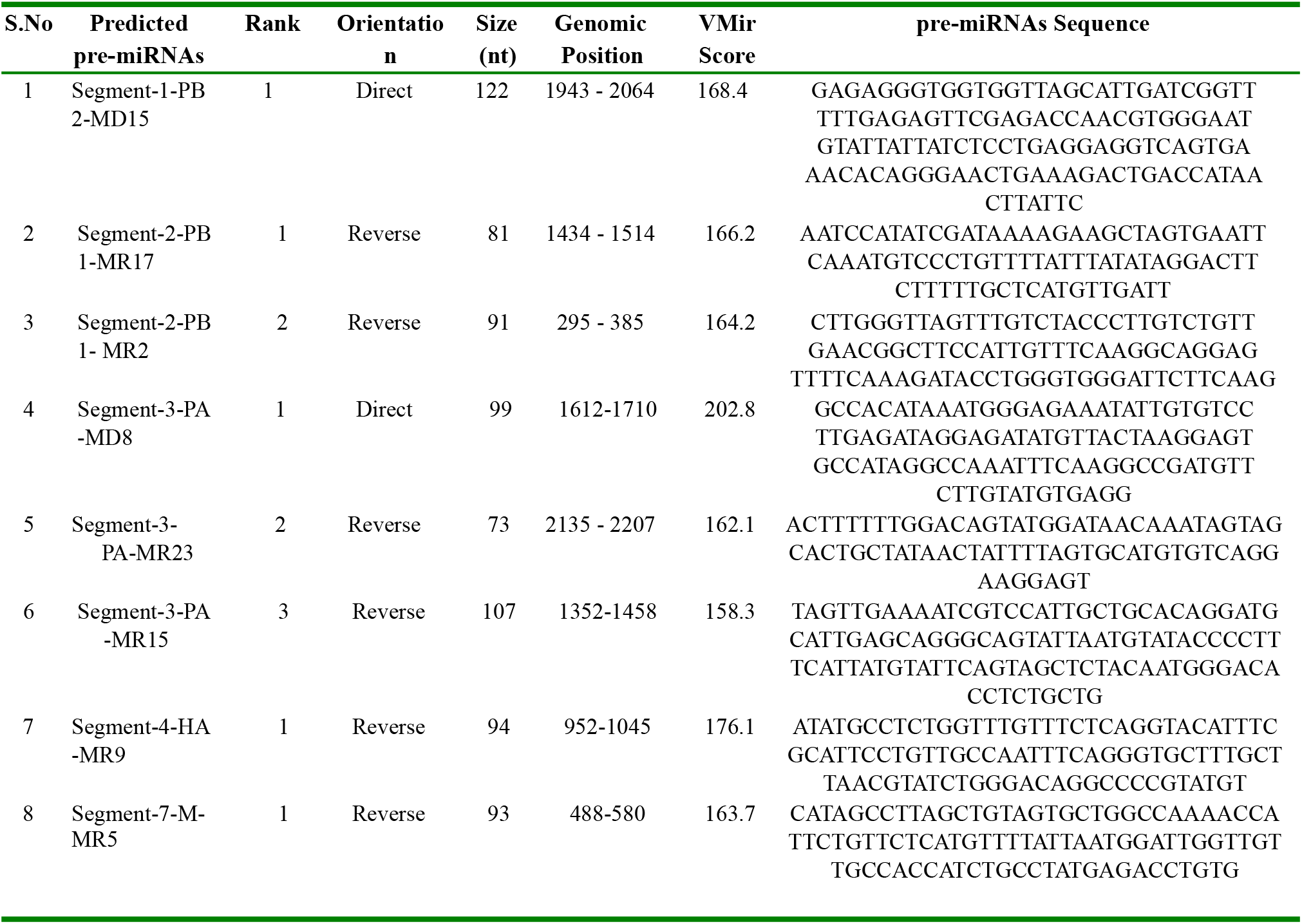
pre-miRNAs rank, orientation, size, genomic position, score, and sequence as predicted by VMir.

Ab initio-based prediction methods often produce false-positive pre-miRNA candidates because pseudo-hairpin loop structures are frequently detected during screening. [20,21]. To improve reliability, the eight predicted pre-miRNAs were further analyzed for their minimum free energy (MFE) using the Mfold algorithm (Table 2). The thermodynamic stability of the secondary structure is a key feature of genuine pre-miRNAs; therefore, estimating the MFE serves as an additional validation step to evaluate the structural feasibility and potential biological significance of the predicted candidates [22].

**Table 2:** Minimum Free Energy (MFE) of potential pre-miRNAs calculated by Mfold.

| Predicted pre-miRNA | Potential | Minimum Free Energy (MFE) ($-\Delta G$.kcal/mol) |
| --- | --- | --- |
| Segment-1-PB2- MD15 | Potential | -34.80 |
| Segment-2-PB1- MR17 | Potential | -11.70 |
| Segment-2-PB1- MR2 | Potential | -24.30 |
| Segment-3-PA- MD8 | Potential | -30.80 |
| Segment-3-PA- MR23 | Potential | -18.10 |
| Segment-3-PA- MR15 | Potential | -29.80 |
| Segment-4-HA- MR9 | Potential | -28.50 |
| Segment-7-M- MR5 | Potential | -26.40 |

After calculating the MFE of pre-miRNAs, the Mature Bayes server was used to retrieve mature miRNA sequences. A total of 16 mature miRNAs were obtained from 8 precursors at both 5’ and 3’ stem locations, as shown in Table 3. Both strands were retained for target prediction, as mentioned in the literature; either can be incorporated into the RNA-Induced Silencing Complex (RISC) [23].

**Table 3:**
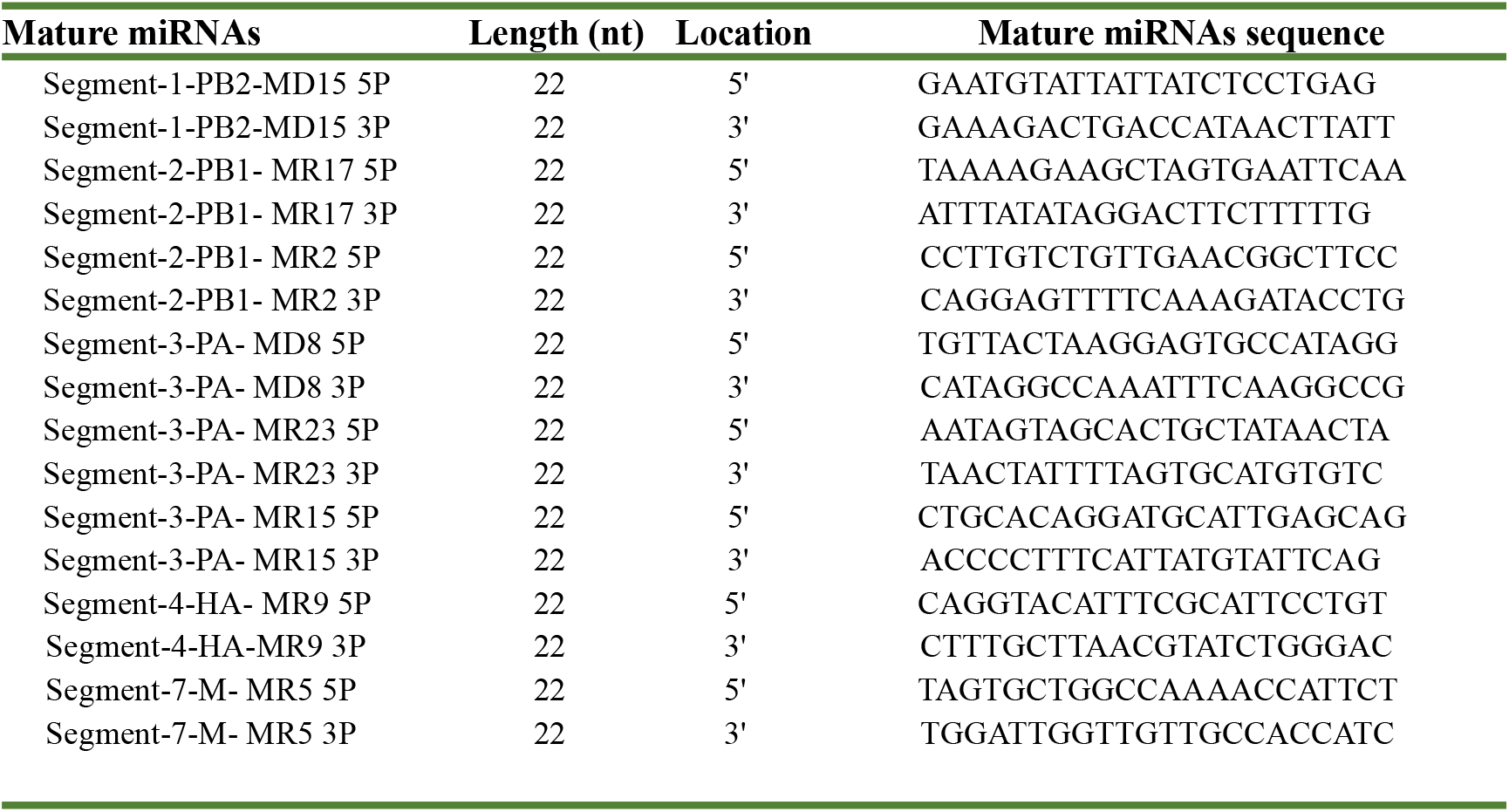
Mature miRNAs length, location, and sequence as predicted by MatureBayes.

The mature miRNA sequences were used in the miRDB custom prediction module to identify targets in human 3′ UTRs. Target genes with prediction scores above 80 were retained for further analysis, as scores above this threshold are generally considered highly reliable and typically do not require additional validation [24]. We predicted 414 genes in Segment 1 (PB 2), 2712 genes in Segment 2 (PB 1), 674 genes in Segment 3 (PA), 527 genes in Segment 4 (HA), and 166 genes in Segment 7 (M) (Supplementary Table S1).

Target genes identified across all segments were combined, resulting in a total of 3747 genes. These genes were then analyzed using the PANTHER database for Gene Ontology (GO) classification to determine their distribution across the three main functional domains. In the Biological Process category, most genes were assigned to cellular process (GO: 0009987), biological regulation (GO: 0065007), and metabolic process (GO: 0008152). Genes were also categorized under response to stimulus (GO: 0050896) and immune system process (GO: 0002376), suggesting potential roles in host immune responses during influenza infection.

Molecular Function analysis showed that binding (GO: 0005488) and catalytic activity (GO: 0003824) were the most common functional classes, followed by transcription regulator activity (GO: 0140110), transporter activity (GO: 0005215), and molecular function regulator activity (GO: 0098772). Translation regulator activity (GO: 0045182) was also observed. In the Cellular Component category, the target genes were mainly associated with protein-containing complex (GO:0032991) and cellular anatomical entity (GO:0110165). The target genes were further analyzed for pathway enrichment using Enrichr, which identified 347 KEGG pathways. Pathways were ranked by adjusted p-values, with those having adjusted p-values < 0.001 considered significantly enriched. The top enriched pathways included MicroRNAs in cancer (hsa 05206), MAPK signaling pathway (hsa 04010), ERBB signaling pathway (hsa 04012), and cGMP-PKG signaling pathway (hsa 04022) (Figure 1).

**Figure 1:**
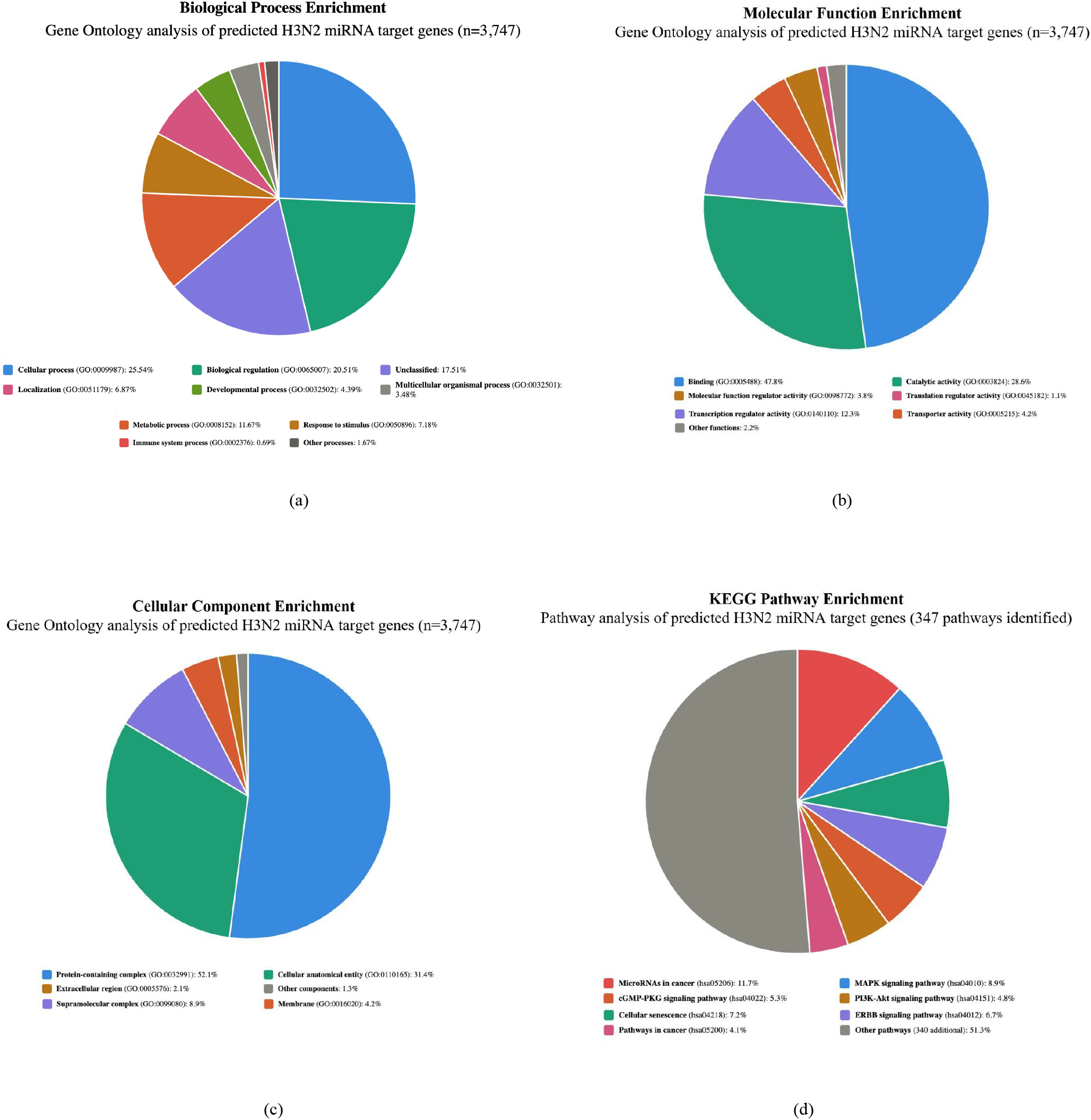
Gene Ontology (GO) and KEGG pathway enrichment analysis of predicted H3N2 miRNA target genes. Functional annotation of 3,747 target genes predicted by miRDB (score >80) across five H3N2 genome segments (PB2, PB1, PA, HA, M). **(a) Biological Process enrichment** shows predominant representation in cellular processes (25.54%, GO:0009987), biological regulation (20.51%, GO:0065007), and metabolic processes (11.67%, GO:0008152). Notably, 7.18% of genes are involved in response to stimulus (GO:0050896) and 0.69% in immune system processes (GO:0002376), indicating potential roles in host-virus interactions. **(b) Molecular Function analysis** reveals binding activity as the most abundant function (47.8%, GO:0005488), followed by catalytic activity (28.6%, GO:0003824) and transcription regulator activity (12.3%, GO:0140110). Translation regulator activity (1.1%, GO:0045182) represents a smaller but functionally significant category for viral regulation. **(c) Cellular Component distribution** shows enrichment in protein-containing complexes (52.1%, GO:0032991) and cellular anatomical entities (31.4%, GO:0110165), consistent with targeting of essential cellular machinery. **(d) KEGG pathway enrichment** (347 total pathways identified) highlights microRNAs in cancer (11.7%, hsa05206) as the top-enriched pathway, followed by MAPK signaling (8.9%, hsa04010), cellular senescence (7.2%, hsa04218), and ERBB signaling (6.7%, hsa04012). Enrichment analysis performed using PANTHER (Fisher’s exact test) and Enrichr with Benjamini-Hochberg FDR correction. Pathways with adjusted p-values <0.001 were considered significantly enriched. The predominance of cancer-related and cell-signaling pathways suggests that viral miRNAs may modulate host cell survival and proliferation mechanisms during infection.

From the predicted target genes for each H3N2 segment, two genes per segment were systematically selected based on their involvement in influenza-related pathways and supporting literature. The chosen genes and their documented roles in viral pathogenesis are summarized in Table 4.

**Table 4:**
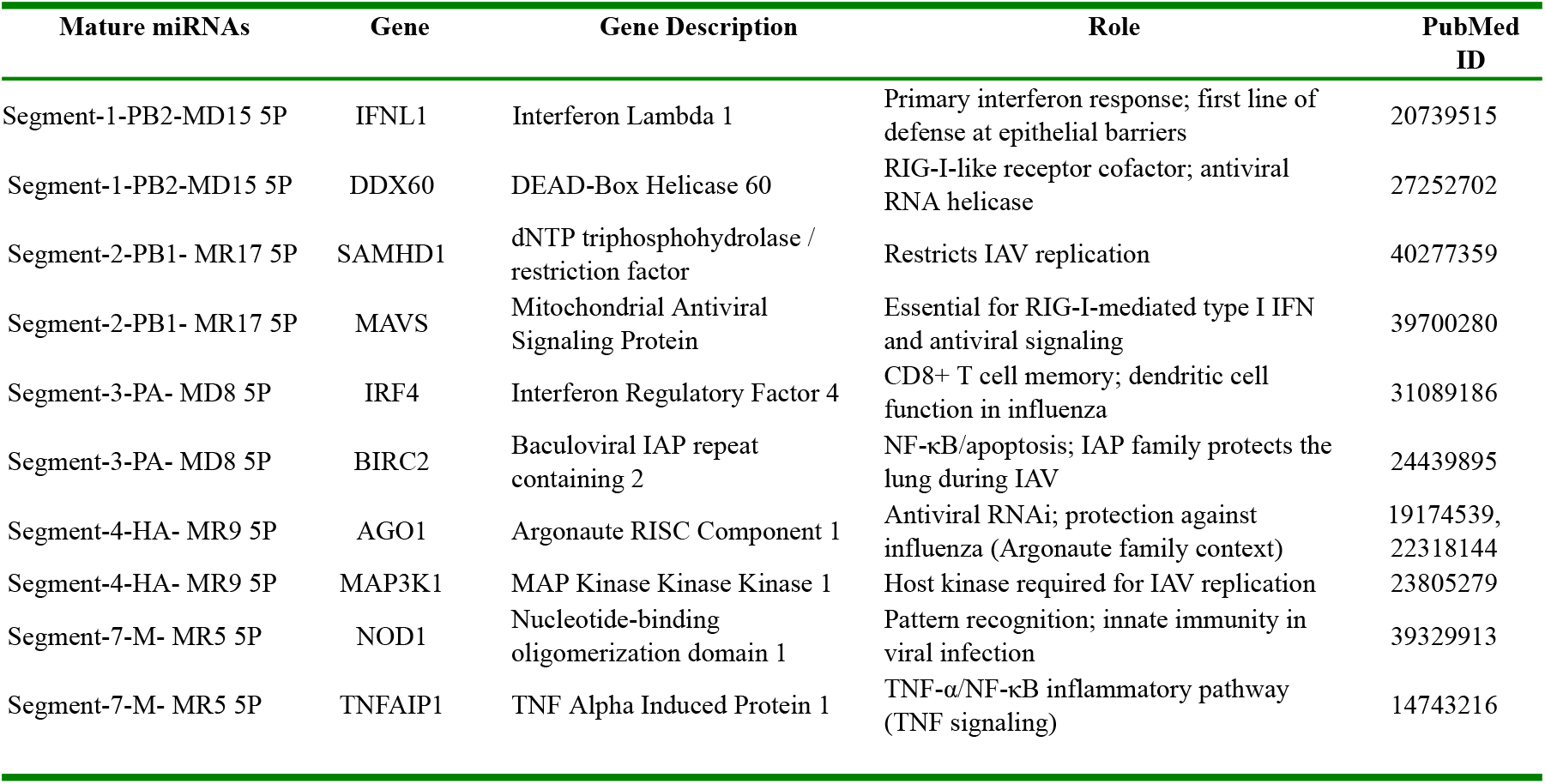
Screened target genes from H3N2 segments and their role with literature evidence.

Notably, CADM2 (Cell Adhesion Molecule 2) was identified as a common target across all five segments (Figure 2, Table 5).

**Figure 2:**
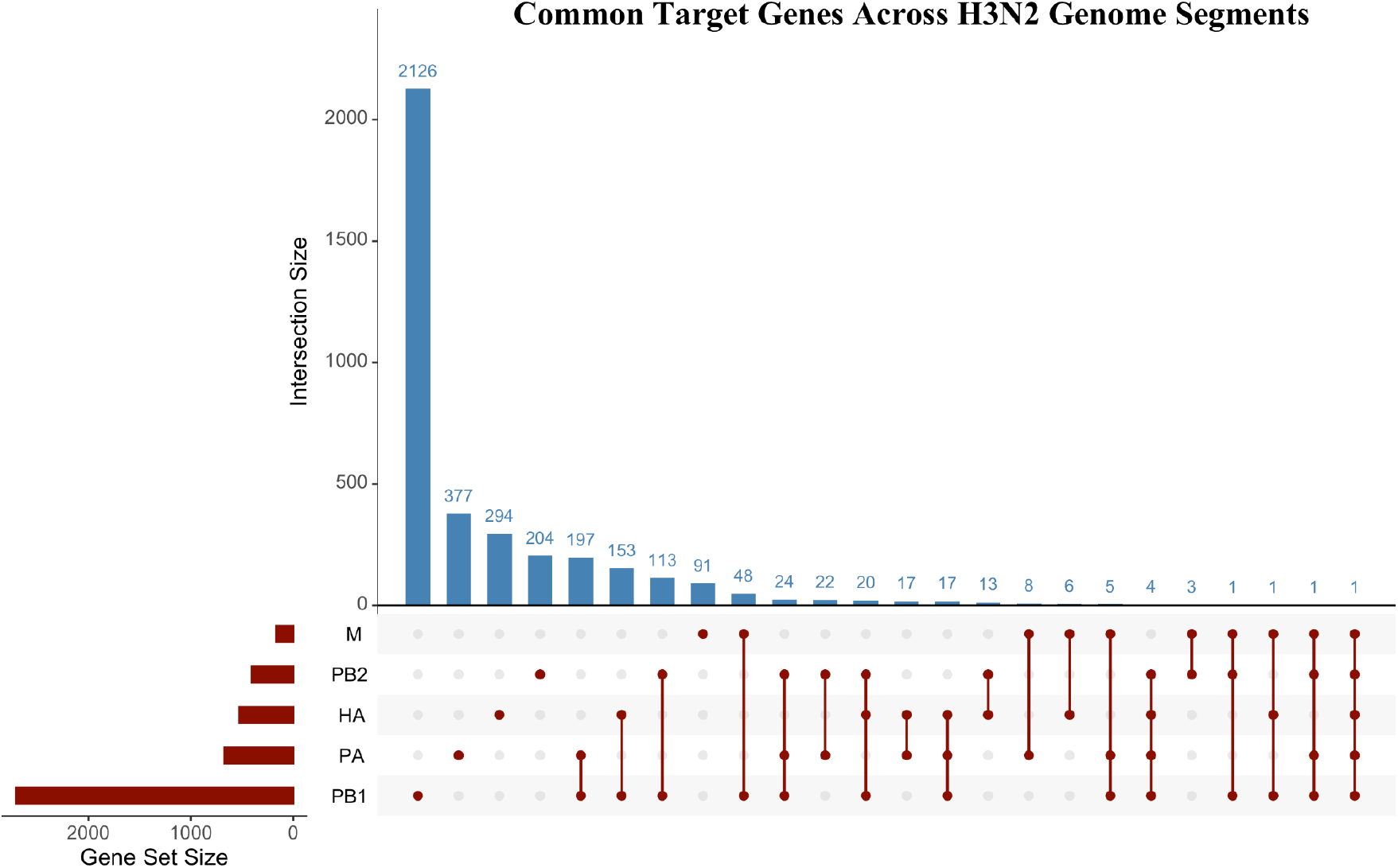
Identification of common target genes across H3N2 genome segments. UpSet plot visualization displaying the intersection patterns of predicted target genes across five analyzed H3N2 segments (PB2: 414 genes, PB1: 2,712 genes, PA: 674 genes, HA: 527 genes, M: 166 genes). (Top bar chart) shows intersection sizes, with the largest group (2,126 genes) representing segment-specific targets found in only one segment. (Bottom matrix) indicates which segments contribute to each intersection, with filled circles showing segment participation. The analysis reveals predominantly segment-specific targeting (>95% of predictions), with only small overlaps between segments. Notably, CADM2 (Cell Adhesion Molecule 2) emerges as the sole gene predicted to be targeted across all five segments (intersection size = 1), suggesting a conserved viral strategy to regulate this cell adhesion molecule, which is involved in viral entry mechanisms. The sparse intersection pattern indicates that viral miRNAs from different H3N2 segments largely target distinct host pathways, reflecting specialized regulatory functions of segment-specific viral miRNAs in host-virus interactions.

**Table 5:** Common target genes across all H3N2 segments and their role with literature evidence.

| Common Gene | Gene Description | PubmedID | Role | miRDB Target scores |
| --- | --- | --- | --- | --- |
| CADM2 | Cell Adhesion Molecule 2 | PMC10231178 | Cell adhesion; hemagglutinin (HA) proteins for cell entry | 88 in Segment-1-PB2-MD15 5P<br>100 in Segment-2-PB1- MR17 5P<br>91 in Segment-3-PA- MR23 3P<br>89 in Segment-4-HA-MR9 3P<br>81 in Segment-7-M- MR5 5P |

The UpSet analysis revealed minimal overlap between segments, with 95.2% of predicted targets being segment-specific, emphasizing distinct regulatory roles.

## DISCUSSION

In the genomic age, computational prediction has become a key method for exploring miRNA biology, complementing traditional cloning and expression-based techniques. Our ab initio pipeline (VMir, MatureBayes, miRDB) enabled rapid, genome-wide screening of H3N2 segments and identified a specific set of host genes that may plausibly be regulated by viral miRNAs.

We found that several predicted miRNAs from the polymerase segments (PB2, PB1) target genes involved in interferon and RIG-I-like receptor signaling, such as IFNL1, DDX60, SAMHD1, and MAVS [25]. IFNL1 encodes type III interferon, which provides frontline antiviral protection at epithelial barriers [26, 27], while DDX60 and MAVS are key mediators of RLR-dependent interferon production [28,29,30]. SAMHD1 has been reported to limit influenza replication [31, 32]. These findings suggest that H3N2-encoded miRNAs may weaken early innate sensing and interferon responses, thereby promoting efficient viral replication in the respiratory epithelium.

We also identified genes linking viral miRNAs to RNA interference, T-cell memory, and lung pathology. AGO1 is a key component of the RNAi machinery that can both restrict viruses and be exploited by them [33,34]. IRF4 is essential for optimal CD8^+^ T-cell memory responses during influenza infection [35]. BIRC2 (a member of the IAP family) and MAP3K1 participate in the MAPK and NF-κB signaling pathways, which regulate cell death and the production of inflammatory cytokines [36,37,38]. Targeting these molecules could help H3N2 balance viral replication with host survival by modulating apoptosis, inflammation, and adaptive immune responses. Finally, NOD1, TNFAIP1, and CADM2 reveal additional layers of regulation involving pattern recognition, TNF/NF-κB signaling, and cell adhesion. NOD1 helps detect intracellular danger signals [39], while TNFAIP1 is involved in TNF-dependent signaling pathways [40]. CADM2, a cell adhesion molecule reported to interact with the hemagglutinin of measles virus, is predicted to be a common target across all analyzed H3N2 segments [41]. The coordinated regulation of these genes by viral miRNAs could potentially influence inflammatory responses and facilitate viral spread within the lung.

This work identifies IFNL1, DDX60, SAMHD1, MAVS, IRF4, BIRC2, AGO1, MAP3K1, NOD1, TNFAIP1, and CADM2 as key host nodes that H3N2-derived miRNAs may target to influence innate and adaptive immunity and contribute to lung pathology. Although these predictions need experimental validation, they provide specific, testable hypotheses about how H3N2-derived miRNAs could play a role in acute respiratory disease and recurrent seasonal infections.

Limitations of this study include the in silico approach to miRNA prediction, the lack of expression data for the predicted viral miRNAs, and the use of broad target prediction thresholds that may yield false positives. Future research should aim to detect these miRNAs during infection, verify direct targeting of the highlighted genes, and assess their functional effects in relevant respiratory cell types and in vivo models.

## CONCLUSION

This study used a genome-wide computational pipeline (VMir, MatureBayes, miRDB) to analyze five H3N2 segments and identified 8 pre-miRNAs, 16 mature miRNAs, and target gene sets ranging from 166 (Segment 7) to 2712 (Segment 2) genes per segment. One gene (CADM2) was common across all segments. From these targets, we selected 10 segment-specific genes involved in interferon and RLR signaling, antiviral restriction, RNA interference, immune cell regulation, and inflammatory responses, with recent literature supporting their roles in antiviral innate and pulmonary immunity. These predictions provide a basis for experimental validation and could aid in understanding IAV H3N2 pathogenesis and developing antiviral strategies. The computational framework described here can also be applied to other influenza A virus subtypes (e.g., H1N1) and strains.

This study has several important limitations that should be considered when interpreting the results. First, the purely computational nature of this analysis means that the predicted miRNAs have not been experimentally validated. Ab initio miRNA prediction methods, while powerful, are known to produce false positives due to the prevalence of pseudo-hairpin structures in genomic sequences. The MFE calculations provide additional structural validation, but experimental confirmation through RNA-seq or Northern blot analysis of infected cells would be essential to validate these predictions. Second, our analysis excluded three viral segments (NP, NA, NS) due to low VMir scores, potentially missing biologically relevant miRNAs that may not conform to typical miRNA structural features. Third, the target prediction relies solely on miRDB scores >80, which, while highly reliable, may miss functionally important targets with lower computational scores but biological relevance. The exclusive focus on 3’UTR targeting also overlooks potential coding sequence or 5’UTR targets.

If experimentally validated, the identified viral miRNAs and their host targets could inform therapeutic strategies. The targeting of interferon signaling components (IFNL1, DDX60, MAVS) suggests the potential to enhance antiviral responses by inhibiting miRNAs. Antisense oligonucleotides targeting viral miRNAs could theoretically restore host antiviral gene expression. Additionally, the common targeting of CADM2 across all segments suggests a conserved viral strategy that could be exploited for broad-spectrum antiviral development.

## AUTHOR CONTRIBUTIONS

**MAS**: Conceptualization, methodology design, data curation, computational analysis, formal analysis, validation, visualization, writing-original draft preparation, writing-review and editing. **HK**: Computational analysis, data processing, validation, writing, review, and editing. **MM**: Conceptualization, methodology design, supervision, project administration, funding acquisition, resources, writing-review and editing, corresponding author responsibilities. All authors have read and agreed to the published version of the manuscript.

## FUNDING

This research was conducted as part of OmicsLogic’s internal research and development program. No external funding was received for this study.

## CONFLICTS OF INTEREST

MM is the founder and CEO of OmicsLogic Bioinformatics & Data Science India Private Limited and OmicsLogic Inc. HK is employed by OmicsLogic. MAS declares no competing interests. The authors declare that the research was conducted in the absence of any commercial or financial relationships that could be construed as a potential conflict of interest.

## DATA AVAILABILITY STATEMENT

The H3N2 genome sequences used in this study are publicly available from the NCBI genome database (accession number: GCA_039834415.1). The computational pipeline and predicted miRNA sequences are available upon reasonable request from the corresponding author. All software tools used are publicly available: VMir (v2.3), MatureBayes, miRDB, PANTHER, and Enrichr.

## ETHICS STATEMENT

This study used only publicly available genomic sequences and computational analyses. No human subjects, animal experiments, or clinical data were involved. Therefore, ethics approval was not required for this research.

## ACKNOWLEDGMENTS

The authors thank the developers of VMir, MatureBayes, miRDB, PANTHER, and Enrichr for making their tools publicly available. We acknowledge the NCBI for providing access to the H3N2 genomic sequences. We also thank the broader bioinformatics community for their continued efforts in developing computational tools for viral genomics research.

## Notes

https://www.ncbi.nlm.nih.gov/genome/

### Competing Interest Statement

The authors have declared no competing interest.

