## Supplemental Data for "Genome-wide computational prediction of miRNAs encoded by influenza A virus (H3N2) predicts target genes involved in pulmonary and antiviral innate immunity"

### Supplementary Data

#### Supplementary Table S1. List of target gene predicted by miRDB.

| Mature miRNA | Gene Symbol | Gene Description |
|---|---|---|
| Segment-1-PB2-MD15 5P | INO80D | INO80 complex subunit D |
| | RGS4 | regulator of G protein signaling 4 |
| | AUTS2 | AUTS2, activator of transcription and developmental regulator |
| | LHFPL6 | LHFPL tetraspan subfamily member 6 |
| | JMJD1C | jumonji domain containing 1C |
| | EIF5B | eukaryotic translation initiation factor 5B |
| | MBNL3 | muscleblind like splicing regulator 3 |
| | HELQ | helicase, POLQ like |
| | ZFP82 | ZFP82 zinc finger protein |
| | UBR3 | ubiquitin protein ligase E3 component n-recognin 3 |
| | MBLAC2 | metallo-beta-lactamase domain containing 2 |
| | ANKRD17 | ankyrin repeat domain 17 |
| | MYO1B | myosin IB |
| | HAPLN1 | hyaluronan and proteoglycan link protein 1 |
| | ADCY7 | adenylate cyclase 7 |
| | CDK14 | cyclin dependent kinase 14 |
| | RFX5 | regulatory factor X5 |
| | LYPLAL1 | lysophospholipase like 1 |
| | ARPP21 | cAMP regulated phosphoprotein 21 |
| | FAM160B1 | family with sequence similarity 160 member B1 |
| | SH2D1B | SH2 domain containing 1B |
| | ZFHX4 | zinc finger homeobox 4 |
| | YIPF5 | Yip1 domain family member 5 |
| | ELAVL2 | ELAV like RNA binding protein 2 |
| | FGB | fibrinogen beta chain |
| | TPST1 | tyrosylprotein sulfotransferase 1 |
| | TENT4B | terminal nucleotidyltransferase 4B |
| | SCAMP1 | secretory carrier membrane protein 1 |
| | ZMYM2 | zinc finger MYM-type containing 2 |
| | RIC1 | RIC1 homolog, RAB6A GEF complex partner 1 |
| | WDR26 | WD repeat domain 26 |
| | IFNL1 | interferon lambda 1 |
| | ERBIN | erbb2 interacting protein |
| | MEOX2 | mesenchyme homeobox 2 |
| | IRF2BP2 | interferon regulatory factor 2 binding protein 2 |
| | FAM126B | family with sequence similarity 126 member B |
| | ADAMTS1 | ADAM metallopeptidase with thrombospondin type 1 motif 1 |
| | IL18R1 | interleukin 18 receptor 1 |

| | | |
|---|---|---|
| | ETV1 | ETS variant 1 |
| | SENP6 | SUMO specific peptidase 6 |
| | MANEA | mannosidase endo-alpha |
| | IL15RA | interleukin 15 receptor subunit alpha |
| | GNRHR | gonadotropin releasing hormone receptor |
| | RRAS2 | RAS related 2 |
| | SGCZ | sarcoglycan zeta |
| | ADAR | adenosine deaminase, RNA specific |
| | MRPL22 | mitochondrial ribosomal protein L22 |
| | N4BP2 | NEDD4 binding protein 2 |
| | SPDL1 | spindle apparatus coiled-coil protein 1 |
| | NAPG | NSF attachment protein gamma |
| | PDP1 | pyruvate dehyrogenase phosphatase catalytic subunit 1 |
| | HBS1L | HBS1 like translational GTPase |
| | SLC7A11 | solute carrier family 7 member 11 |
| | STX2 | syntaxin 2 |
| | SLC8A3 | solute carrier family 8 member A3 |
| | STRIP1 | striatin interacting protein 1 |
| | NDRG4 | NDRG family member 4 |
| | DHX16 | DEAH-box helicase 16 |
| | MIGA1 | mitoguardin 1 |
| | ARFGEF2 | ADP ribosylation factor guanine nucleotide exchange factor 2 |
| | ZBTB20 | zinc finger and BTB domain containing 20 |
| | NRK | Nik related kinase |
| | UNK | unk zinc finger |
| | TTPA | alpha tocopherol transfer protein |
| | DUSP19 | dual specificity phosphatase 19 |
| | ZNF404 | zinc finger protein 404 |
| | ZNF616 | zinc finger protein 616 |
| | FSD1L | fibronectin type III and SPRY domain containing 1 like |
| | EMC7 | ER membrane protein complex subunit 7 |
| | CBX7 | chromobox 7 |
| | CCR3 | C-C motif chemokine receptor 3 |
| | HMGXB4 | HMG-box containing 4 |
| | P2RY12 | purinergic receptor P2Y12 |
| | KIAA0355 | KIAA0355 |
| | TMED5 | transmembrane p24 trafficking protein 5 |
| | FRMD4A | FERM domain containing 4A |
| | FNDC3B | fibronectin type III domain containing 3B |
| | CHIC2 | cysteine rich hydrophobic domain 2 |
| | TSHZ1 | teashirt zinc finger homeobox 1 |
| | CTTNBP2NL | CTTNBP2 N-terminal like |

| | | |
|---|---|---|
| | HDAC9 | histone deacetylase 9 |
| | DISC1 | DISC1 scaffold protein |
| | FNIP1 | folliculin interacting protein 1 |
| | BCL2 | BCL2, apoptosis regulator |
| | PTPRF | protein tyrosine phosphatase, receptor type F |
| | ZMYND11 | zinc finger MYND-type containing 11 |
| | MYO6 | myosin VI |
| | SLC39A8 | solute carrier family 39 member 8 |
| | INSM1 | INSM transcriptional repressor 1 |
| | SOX7 | SRY-box 7 |
| | GPR180 | G protein-coupled receptor 180 |
| | VTA1 | vesicle trafficking 1 |
| | SLC28A1 | solute carrier family 28 member 1 |
| | DMRT3 | doublesex and mab-3 related transcription factor 3 |
| | ARHGEF16 | Rho guanine nucleotide exchange factor 16 |
| | PDZD2 | PDZ domain containing 2 |
| | HDGFL3 | HDGF like 3 |
| | SPARC | secreted protein acidic and cysteine rich |
| | SEPHS1 | selenophosphate synthetase 1 |
| | STAMBPL1 | STAM binding protein like 1 |
| | GABRB3 | gamma-aminobutyric acid type A receptor beta3 subunit |
| | POLR2M | RNA polymerase II subunit M |
| | GCOM1 | GRINL1A complex locus 1 |
| | CUL3 | cullin 3 |
| | AAK1 | AP2 associated kinase 1 |
| | FAM216B | family with sequence similarity 216 member B |
| | HIPK1 | homeodomain interacting protein kinase 1 |
| | DDX60 | DExD/H-box helicase 60 |
| | CADM2 | cell adhesion molecule 2 |
| | SEC24D | SEC24 homolog D, COPII coat complex component |
| | SEH1L | SEH1 like nucleoporin |
| | STOX1 | storkhead box 1 |
| | PHC3 | polyhomeotic homolog 3 |
| | RAB11FIP2 | RAB11 family interacting protein 2 |
| | EHD3 | EH domain containing 3 |
| | FUT9 | fucosyltransferase 9 |
| | YTHDC1 | YTH domain containing 1 |
| | PDE6D | phosphodiesterase 6D |
| | PPP1R12A | protein phosphatase 1 regulatory subunit 12A |
| | HMGCLL1 | 3-hydroxymethyl-3-methylglutaryl-CoA lyase like 1 |
| | KRAS | KRAS proto-oncogene, GTPase |
| | SGK3 | serum/glucocorticoid regulated kinase family member 3 |

| | | |
|---|---|---|
| | CD44 | CD44 molecule (Indian blood group) |
| | NCKAP1 | NCK associated protein 1 |
| | CDC37L1 | cell division cycle 37 like 1 |
| | CLU | clusterin |
| | ZSWIM6 | zinc finger SWIM-type containing 6 |
| | COBLL1 | cordon-bleu WH2 repeat protein like 1 |
| | TMEM132B | transmembrane protein 132B |
| | ARID4A | AT-rich interaction domain 4A |
| | PRKAR1A | protein kinase cAMP-dependent type I regulatory subunit alpha |
| | DLC1 | DLC1 Rho GTPase activating protein |
| | PRTG | protogenin |
| | HYDIN | HYDIN, axonemal central pair apparatus protein |
| | DERL2 | derlin 2 |
| | ARMCX4 | armadillo repeat containing X-linked 4 |
| | CBLL1 | Cbl proto-oncogene like 1 |
| | SLC18A1 | solute carrier family 18 member A1 |
| | HERC4 | HECT and RLD domain containing E3 ubiquitin protein ligase 4 |
| | NXT2 | nuclear transport factor 2 like export factor 2 |
| | SLC30A7 | solute carrier family 30 member 7 |
| | PCDH11Y | protocadherin 11 Y-linked |
| | PCDH11X | protocadherin 11 X-linked |
| | XPO7 | exportin 7 |
| | GABRA4 | gamma-aminobutyric acid type A receptor alpha4 subunit |
| | ESF1 | ESF1 nucleolar pre-rRNA processing protein homolog |
| | SLC25A24 | solute carrier family 25 member 24 |
| | TCFL5 | transcription factor like 5 |
| | STXBP4 | syntaxin binding protein 4 |
| | ATP2B1 | ATPase plasma membrane Ca2+ transporting 1 |
| | C21orf91 | chromosome 21 open reading frame 91 |
| | GABRG1 | gamma-aminobutyric acid type A receptor gamma1 subunit |
| | CLVS2 | clavesin 2 |
| | APPL1 | adaptor protein, phosphotyrosine interacting with PH domain and leucine zipper 1 |
| | TRIM33 | tripartite motif containing 33 |
| | SETD9 | SET domain containing 9 |
| | FREM2 | FRAS1 related extracellular matrix protein 2 |
| | MFSD14B | major facilitator superfamily domain containing 14B |
| | TENT2 | terminal nucleotidyltransferase 2 |
| | RNASEL | ribonuclease L |
| | ZNF454 | zinc finger protein 454 |
| | MGAT4A | alpha-1,3-mannosyl-glycoprotein 4-beta-N-acetylglucosaminyltransferase A |

| | | |
|---|---|---|
| | RRM2B | ribonucleotide reductase regulatory TP53 inducible subunit M2B |
| | UBN2 | ubinuclein 2 |
| | RNF219 | ring finger protein 219 |
| | GLI2 | GLI family zinc finger 2 |
| | ZBTB34 | zinc finger and BTB domain containing 34 |
| | GLI3 | GLI family zinc finger 3 |
| | H3F3A | H3 histone family member 3A |
| | FAM161A | FAM161A, centrosomal protein |
| | OPRM1 | opioid receptor mu 1 |
| | BIN2 | bridging integrator 2 |
| | BRWD3 | bromodomain and WD repeat domain containing 3 |
| | MTCP1 | mature T cell proliferation 1 |
| | ZER1 | zyg-11 related cell cycle regulator |
| | ZNF714 | zinc finger protein 714 |
| | EBPL | EBP like |
| | CDK12 | cyclin dependent kinase 12 |
| | REST | RE1 silencing transcription factor |
| | MCTP1 | multiple C2 and transmembrane domain containing 1 |
| | AKAP5 | A-kinase anchoring protein 5 |
| | NAA20 | N(alpha)-acetyltransferase 20, NatB catalytic subunit |
| | SPATA9 | spermatogenesis associated 9 |
| | ZBTB11 | zinc finger and BTB domain containing 11 |
| | PROX1 | prospero homeobox 1 |
| | DNAJB11 | DnaJ heat shock protein family (Hsp40) member B11 |
| | FHIT | fragile histidine triad |
| | STAM | signal transducing adaptor molecule |
| | HAND1 | heart and neural crest derivatives expressed 1 |
| | DPYD | dihydropyrimidine dehydrogenase |
| | DNAJB6 | DnaJ heat shock protein family (Hsp40) member B6 |
| | PATJ | PATJ, crumbs cell polarity complex component |
| | WNK3 | WNK lysine deficient protein kinase 3 |
| | DBF4 | DBF4 zinc finger |
| | NSMAF | neutral sphingomyelinase activation associated factor |
| | MON2 | MON2 homolog, regulator of endosome-to-Golgi trafficking |
| | CD24 | CD24 molecule |
| | MYB | MYB proto-oncogene, transcription factor |
| | YTHDF3 | YTH N6-methyladenosine RNA binding protein 3 |
| | DGKH | diacylglycerol kinase eta |
| | CD2AP | CD2 associated protein |
| | C2orf49 | chromosome 2 open reading frame 49 |
| | VDAC2 | voltage dependent anion channel 2 |

| | | |
|---|---|---|
| | MCPH1 | microcephalin 1 |
| | GRIP1 | glutamate receptor interacting protein 1 |
| | FGF12 | fibroblast growth factor 12 |
| | ZNF12 | zinc finger protein 12 |
| | SNX2 | sorting nexin 2 |
| | PCDH7 | protocadherin 7 |
| | DPH6 | diphthamine biosynthesis 6 |
| | CDK2AP1 | cyclin dependent kinase 2 associated protein 1 |
| | ARMH4 | armadillo-like helical domain containing 4 |
| | TFAP2B | transcription factor AP-2 beta |
| | ACYP2 | acylphosphatase 2 |
| | KLHL15 | kelch like family member 15 |
| | RANBP3L | RAN binding protein 3 like |
| | CEP97 | centrosomal protein 97 |
| | VPS29 | VPS29, retromer complex component |
| | ATP23 | ATP23 metallopeptidase and ATP synthase assembly factor homolog |
| | SHOX2 | short stature homeobox 2 |
| | PCF11 | PCF11, cleavage and polyadenylation factor subunit |
| | ADGRF5 | adhesion G protein-coupled receptor F5 |
| | ZNF843 | zinc finger protein 843 |
| | INTS13 | integrator complex subunit 13 |
| | KCNG3 | potassium voltage-gated channel modifier subfamily G member 3 |
| | LMX1A | LIM homeobox transcription factor 1 alpha |
| | GZF1 | GDNF inducible zinc finger protein 1 |
| | CENPE | centromere protein E |
| | COPS8 | COP9 signalosome subunit 8 |
| | EEF1AKMT2 | EEF1A lysine methyltransferase 2 |
| | CHD6 | chromodomain helicase DNA binding protein 6 |
| | ZNF621 | zinc finger protein 621 |
| | PPM1B | protein phosphatase, Mg2+/Mn2+ dependent 1B |
| | C5orf63 | chromosome 5 open reading frame 63 |
| | PHEX | phosphate regulating endopeptidase homolog X-linked |
| | MGAT4B | alpha-1,3-mannosyl-glycoprotein 4-beta-N-acetylglucosaminyltransferase B |
| | RUNDC3B | RUN domain containing 3B |
| | FST | follistatin |
| | MTX2 | metaxin 2 |
| | CDH2 | cadherin 2 |
| | DST | dystonin |
| | CSMD3 | CUB and Sushi multiple domains 3 |
| | TMEM128 | transmembrane protein 128 |

| | | |
|---|---|---|
| | CCNT2 | cyclin T2 |
| | SLF2 | SMC5-SMC6 complex localization factor 2 |
| | NPNT | nephronectin |
| | SARAF | store-operated calcium entry associated regulatory factor |
| | QKI | QKI, KH domain containing RNA binding |
| | EIF1AD | eukaryotic translation initiation factor 1A domain containing |
| | GRIA3 | glutamate ionotropic receptor AMPA type subunit 3 |
| | RICTOR | RPTOR independent companion of MTOR complex 2 |
| | CAV2 | caveolin 2 |
| | RAB21 | RAB21, member RAS oncogene family |
| Segment-1-PB2-MD15 3P | CSTF2T | cleavage stimulation factor subunit 2 tau variant |
| | FAM214A | family with sequence similarity 214 member A |
| | TRIM5 | tripartite motif containing 5 |
| | HLF | HLF, PAR bZIP transcription factor |
| | EAF1 | ELL associated factor 1 |
| | ABLIM1 | actin binding LIM protein 1 |
| | SEC14L1 | SEC14 like lipid binding 1 |
| | CNOT6 | CCR4-NOT transcription complex subunit 6 |
| | IPMK | inositol polyphosphate multikinase |
| | SINHCAF | SIN3-HDAC complex associated factor |
| | ANKRD27 | ankyrin repeat domain 27 |
| | USP33 | ubiquitin specific peptidase 33 |
| | KIAA1549L | KIAA1549 like |
| | PAN3 | poly(A) specific ribonuclease subunit PAN3 |
| | DCAF10 | DDB1 and CUL4 associated factor 10 |
| | GRK5 | G protein-coupled receptor kinase 5 |
| | EPN2 | epsin 2 |
| | PI4K2B | phosphatidylinositol 4-kinase type 2 beta |
| | HOOK3 | hook microtubule tethering protein 3 |
| | PHTF2 | putative homeodomain transcription factor 2 |
| | ELAVL2 | ELAV like RNA binding protein 2 |
| | OSBPL8 | oxysterol binding protein like 8 |
| | RSRP1 | arginine and serine rich protein 1 |
| | DIAPH3 | diaphanous related formin 3 |
| | MITF | melanocyte inducing transcription factor |
| | CARD8 | caspase recruitment domain family member 8 |
| | ZNF594 | zinc finger protein 594 |
| | KIF5B | kinesin family member 5B |
| | CCPG1 | cell cycle progression 1 |
| | COG6 | component of oligomeric golgi complex 6 |
| | GNAL | G protein subunit alpha L |
| | ZNF624 | zinc finger protein 624 |

| | | |
|---|---|---|
| | TREM1 | triggering receptor expressed on myeloid cells 1 |
| | RC3H1 | ring finger and CCCH-type domains 1 |
| | FAM91A1 | family with sequence similarity 91 member A1 |
| | YWHAQ | tyrosine 3-monooxygenase/tryptophan 5-monooxygenase activation protein theta |
| | CTSC | cathepsin C |
| | DPY19L4 | dpy-19 like 4 |
| | AAK1 | AP2 associated kinase 1 |
| | ATP1B1 | ATPase Na+/K+ transporting subunit beta 1 |
| | ESRP2 | epithelial splicing regulatory protein 2 |
| | ERO1A | endoplasmic reticulum oxidoreductase 1 alpha |
| | AK4 | adenylate kinase 4 |
| | GCH1 | GTP cyclohydrolase 1 |
| | GRM5 | glutamate metabotropic receptor 5 |
| | BBX | BBX, HMG-box containing |
| | SH3RF3 | SH3 domain containing ring finger 3 |
| | STRN3 | striatin 3 |
| | GHITM | growth hormone inducible transmembrane protein |
| | DACH1 | dachshund family transcription factor 1 |
| | LRRC58 | leucine rich repeat containing 58 |
| | PADI1 | peptidyl arginine deiminase 1 |
| | ONECUT2 | one cut homeobox 2 |
| | WDCP | WD repeat and coiled coil containing |
| | EPB41L4B | erythrocyte membrane protein band 4.1 like 4B |
| | C7orf31 | chromosome 7 open reading frame 31 |
| | PCDH9 | protocadherin 9 |
| | IMPAD1 | inositol monophosphatase domain containing 1 |
| | SLC4A4 | solute carrier family 4 member 4 |
| | IMPG2 | interphotoreceptor matrix proteoglycan 2 |
| | LYSMD2 | LysM domain containing 2 |
| | SOX2 | SRY-box 2 |
| | NR4A3 | nuclear receptor subfamily 4 group A member 3 |
| | HDAC9 | histone deacetylase 9 |
| | XRCC2 | X-ray repair cross complementing 2 |
| | PRUNE2 | prune homolog 2 with BCH domain |
| | SUCO | SUN domain containing ossification factor |
| | PSMA5 | proteasome subunit alpha 5 |
| | VDAC1 | voltage dependent anion channel 1 |
| | ZBTB41 | zinc finger and BTB domain containing 41 |
| | PDE6D | phosphodiesterase 6D |
| | TENT4B | terminal nucleotidyltransferase 4B |
| | SLC24A2 | solute carrier family 24 member 2 |

| | | |
|---|---|---|
| | SLC5A3 | solute carrier family 5 member 3 |
| | DAAM1 | dishevelled associated activator of morphogenesis 1 |
| | AGFG1 | ArfGAP with FG repeats 1 |
| | NR2F1 | nuclear receptor subfamily 2 group F member 1 |
| | IGF1 | insulin like growth factor 1 |
| | EOGT | EGF domain specific O-linked N-acetylglucosamine transferase |
| | ZNF451 | zinc finger protein 451 |
| | SUPT7L | SPT7 like, STAGA complex gamma subunit |
| | SNX10 | sorting nexin 10 |
| | TFDP1 | transcription factor Dp-1 |
| | HEY2 | hes related family bHLH transcription factor with YRPW motif 2 |
| | YWHAG | tyrosine 3-monooxygenase/tryptophan 5-monooxygenase activation protein gamma |
| | MGST2 | microsomal glutathione S-transferase 2 |
| | MRAS | muscle RAS oncogene homolog |
| | MRPL17 | mitochondrial ribosomal protein L17 |
| | ZDHHC15 | zinc finger DHHC-type containing 15 |
| | MS4A12 | membrane spanning 4-domains A12 |
| | DSTYK | dual serine/threonine and tyrosine protein kinase |
| | TFAP2A | transcription factor AP-2 alpha |
| | RSBN1L | round spermatid basic protein 1 like |
| | PPTC7 | PTC7 protein phosphatase homolog |
| | PKP2 | plakophilin 2 |
| | CDH11 | cadherin 11 |
| | MAGT1 | magnesium transporter 1 |
| | MSL2 | MSL complex subunit 2 |
| | UBE2E2 | ubiquitin conjugating enzyme E2 E2 |
| | ZNFX1 | zinc finger NFX1-type containing 1 |
| | SPTY2D1 | SPT2 chromatin protein domain containing 1 |
| | RGS7BP | regulator of G protein signaling 7 binding protein |
| | ALS2 | ALS2, alsin Rho guanine nucleotide exchange factor |
| | LOC101927322 | uncharacterized LOC101927322 |
| | CDK14 | cyclin dependent kinase 14 |
| | FBXO3 | F-box protein 3 |
| | RASA2 | RAS p21 protein activator 2 |
| | PIK3AP1 | phosphoinositide-3-kinase adaptor protein 1 |
| | AKT3 | AKT serine/threonine kinase 3 |
| | DDX4 | DEAD-box helicase 4 |
| | PRKCI | protein kinase C iota |
| | MAGI2 | membrane associated guanylate kinase, WW and PDZ domain containing 2 |
| | NIPAL1 | NIPA like domain containing 1 |

| | | |
|---|---|---|
| | CDKL5 | cyclin dependent kinase like 5 |
| | SSR1 | signal sequence receptor subunit 1 |
| | ZNF558 | zinc finger protein 558 |
| | PTCHD1 | patched domain containing 1 |
| | CCL17 | C-C motif chemokine ligand 17 |
| | BAAT | bile acid-CoA:amino acid N-acyltransferase |
| | LIMA1 | LIM domain and actin binding 1 |
| | DIDO1 | death inducer-obliterator 1 |
| | TRMT11 | tRNA methyltransferase 11 homolog |
| | CXCL5 | C-X-C motif chemokine ligand 5 |
| | ZSWIM7 | zinc finger SWIM-type containing 7 |
| | CEP63 | centrosomal protein 63 |
| | SLIT2 | slit guidance ligand 2 |
| | COL24A1 | collagen type XXIV alpha 1 chain |
| | NUP98 | nucleoporin 98 |
| | TRIM13 | tripartite motif containing 13 |
| | NAPG | NSF attachment protein gamma |
| | NOTCH3 | notch 3 |
| | NAP1L1 | nucleosome assembly protein 1 like 1 |
| | LIN28B | lin-28 homolog B |
| | SLC39A3 | solute carrier family 39 member 3 |
| | CCER1 | coiled-coil glutamate rich protein 1 |
| | CNR1 | cannabinoid receptor 1 |
| | USP11 | ubiquitin specific peptidase 11 |
| | LEPROT | leptin receptor overlapping transcript |
| | RER1 | retention in endoplasmic reticulum sorting receptor 1 |
| | KLHL2 | kelch like family member 2 |
| | KBTBD6 | kelch repeat and BTB domain containing 6 |
| | LPAR6 | lysophosphatidic acid receptor 6 |
| | C11orf58 | chromosome 11 open reading frame 58 |
| | ZNF41 | zinc finger protein 41 |
| | WDFY4 | WDFY family member 4 |
| | AHR | aryl hydrocarbon receptor |
| | CTBP2 | C-terminal binding protein 2 |
| | TBPL1 | TATA-box binding protein like 1 |
| | HYAL4 | hyaluronidase 4 |
| | ETV6 | ETS variant 6 |
| | ZNF680 | zinc finger protein 680 |
| | DICER1 | dicer 1, ribonuclease III |
| | DDX3X | DEAD-box helicase 3 X-linked |
| | UBE2W | ubiquitin conjugating enzyme E2 W |
| | PRRC1 | proline rich coiled-coil 1 |

| | | |
|---|---|---|
| | KATNBL1 | katanin regulatory subunit B1 like 1 |
| | RSF1 | remodeling and spacing factor 1 |
| | MAGEB4 | MAGE family member B4 |
| | NHLH2 | nescient helix-loop-helix 2 |
| | NCOR1 | nuclear receptor corepressor 1 |

| | | |
|---|---|---|
| Segment-2-PB1- MR17 5P | BCL11A | BCL11A, BAF complex component |
| | LPP | LIM domain containing preferred translocation partner in lipoma |
| | CD99 | CD99 molecule (Xg blood group) |
| | ZFHX3 | zinc finger homeobox 3 |
| | ELK4 | ELK4, ETS transcription factor |
| | GRM5 | glutamate metabotropic receptor 5 |
| | TBL1XR1 | transducin beta like 1 X-linked receptor 1 |
| | GTF2A1 | general transcription factor IIA subunit 1 |
| | CTNNA2 | catenin alpha 2 |
| | MMP16 | matrix metallopeptidase 16 |
| | TFAP4 | transcription factor AP-4 |
| | B3GALT1 | beta-1,3-galactosyltransferase 1 |
| | NTRK2 | neurotrophic receptor tyrosine kinase 2 |
| | DDX6 | DEAD-box helicase 6 |
| | FAM91A1 | family with sequence similarity 91 member A1 |
| | DIP2C | disco interacting protein 2 homolog C |
| | RAB21 | RAB21, member RAS oncogene family |
| | INO80D | INO80 complex subunit D |
| | DENND5A | DENN domain containing 5A |
| | TMEM170B | transmembrane protein 170B |
| | FBXW11 | F-box and WD repeat domain containing 11 |
| | FEM1C | fem-1 homolog C |
| | DICER1 | dicer 1, ribonuclease III |
| | AHR | aryl hydrocarbon receptor |
| | ATXN3 | ataxin 3 |
| | RDX | radixin |
| | MYADM | myeloid associated differentiation marker |
| | PIGR | polymeric immunoglobulin receptor |
| | CCND2 | cyclin D2 |
| | LCLAT1 | lysocardiolipin acyltransferase 1 |
| | SBNO1 | strawberry notch homolog 1 |
| | TXLNA | taxilin alpha |
| | CACNB4 | calcium voltage-gated channel auxiliary subunit beta 4 |
| | PDS5B | PDS5 cohesin associated factor B |
| | FAM76A | family with sequence similarity 76 member A |
| | KLHL8 | kelch like family member 8 |

| | | |
|---|---|---|
| | RALGPS2 | Ral GEF with PH domain and SH3 binding motif 2 |
| | MSI2 | musashi RNA binding protein 2 |
| | PAK2 | p21 (RAC1) activated kinase 2 |
| | PHF6 | PHD finger protein 6 |
| | FRY | FRY microtubule binding protein |
| | MDFIC | MyoD family inhibitor domain containing |
| | LMAN1 | lectin, mannose binding 1 |
| | RFX3 | regulatory factor X3 |
| | BTBD3 | BTB domain containing 3 |
| | ZNF570 | zinc finger protein 570 |
| | TMEM170A | transmembrane protein 170A |
| | POU2F1 | POU class 2 homeobox 1 |
| | SCAMP1 | secretory carrier membrane protein 1 |
| | UBE2W | ubiquitin conjugating enzyme E2 W |
| | GSK3B | glycogen synthase kinase 3 beta |
| | FZD3 | frizzled class receptor 3 |
| | PARP11 | poly(ADP-ribose) polymerase family member 11 |
| | TBX18 | T-box 18 |
| | PLEKHA5 | pleckstrin homology domain containing A5 |
| | MPZL2 | myelin protein zero like 2 |
| | BROX | BRO1 domain and CAAX motif containing |
| | CANX | calnexin |
| | NFAT5 | nuclear factor of activated T cells 5 |
| | PTPRD | protein tyrosine phosphatase, receptor type D |
| | CDK6 | cyclin dependent kinase 6 |
| | NEUROD6 | neuronal differentiation 6 |
| | KHDRBS2 | KH RNA binding domain containing, signal transduction associated 2 |
| | ZFAND4 | zinc finger AN1-type containing 4 |
| | HECA | hdc homolog, cell cycle regulator |
| | SCAI | suppressor of cancer cell invasion |
| | PXDC1 | PX domain containing 1 |
| | CPS1 | carbamoyl-phosphate synthase 1 |
| | ASPH | aspartate beta-hydroxylase |
| | MEF2C | myocyte enhancer factor 2C |
| | TOP1 | DNA topoisomerase I |
| | CCNY | cyclin Y |
| | MANEA | mannosidase endo-alpha |
| | SDHA | succinate dehydrogenase complex flavoprotein subunit A |
| | GRAMD4 | GRAM domain containing 4 |
| | KIAA1217 | KIAA1217 |
| | KCNJ16 | potassium voltage-gated channel subfamily J member 16 |

| | | |
|---|---|---|
| | SEMA4D | semaphorin 4D |
| | TENT5A | terminal nucleotidyltransferase 5A |
| | TIAM1 | T cell lymphoma invasion and metastasis 1 |
| | PTBP3 | polypyrimidine tract binding protein 3 |
| | ZFX | zinc finger protein X-linked |
| | TMEM33 | transmembrane protein 33 |
| | PDE4D | phosphodiesterase 4D |
| | TMF1 | TATA element modulatory factor 1 |
| | ERBB4 | erb-b2 receptor tyrosine kinase 4 |
| | NEXMIF | neurite extension and migration factor |
| | ZXDB | zinc finger X-linked duplicated B |
| | RAP2A | RAP2A, member of RAS oncogene family |
| | FIGN | fidgetin, microtubule severing factor |
| | PKD2 | polycystin 2, transient receptor potential cation channel |
| | ELAVL2 | ELAV like RNA binding protein 2 |
| | COP1 | COP1, E3 ubiquitin ligase |
| | ZMAT3 | zinc finger matrin-type 3 |
| | HOOK3 | hook microtubule tethering protein 3 |
| | ANO5 | anoctamin 5 |
| | B3GALT2 | beta-1,3-galactosyltransferase 2 |
| | ADGRL3 | adhesion G protein-coupled receptor L3 |
| | STK4 | serine/threonine kinase 4 |
| | ZBTB20 | zinc finger and BTB domain containing 20 |
| | FSD1L | fibronectin type III and SPRY domain containing 1 like |
| | SOCS4 | suppressor of cytokine signaling 4 |
| | EMP1 | epithelial membrane protein 1 |
| | RAB30 | RAB30, member RAS oncogene family |
| | RBM12 | RNA binding motif protein 12 |
| | EBF1 | EBF transcription factor 1 |
| | OSBPL8 | oxysterol binding protein like 8 |
| | MIER3 | MIER family member 3 |
| | LSM12 | LSM12 homolog |
| | PDE4A | phosphodiesterase 4A |
| | CCNT2 | cyclin T2 |
| | TOB2 | transducer of ERBB2, 2 |
| | OPHN1 | oligophrenin 1 |
| | DDX17 | DEAD-box helicase 17 |
| | CADM2 | cell adhesion molecule 2 |
| | UBE2H | ubiquitin conjugating enzyme E2 H |
| | EPHA3 | EPH receptor A3 |
| | GABRA4 | gamma-aminobutyric acid type A receptor alpha4 subunit |
| | FBXO30 | F-box protein 30 |

| | | |
|---|---|---|
| | PEX13 | peroxisomal biogenesis factor 13 |
| | TCP11L1 | t-complex 11 like 1 |
| | FBXO45 | F-box protein 45 |
| | EIF4G3 | eukaryotic translation initiation factor 4 gamma 3 |
| | TENT4B | terminal nucleotidyltransferase 4B |
| | DOK5 | docking protein 5 |
| | ATL2 | atlastin GTPase 2 |
| | ZNF454 | zinc finger protein 454 |
| | RAB2A | RAB2A, member RAS oncogene family |
| | LONRF1 | LON peptidase N-terminal domain and ring finger 1 |
| | YAF2 | YY1 associated factor 2 |
| | SH3TC2 | SH3 domain and tetratricopeptide repeats 2 |
| | PICALM | phosphatidylinositol binding clathrin assembly protein |
| | TLCD2 | TLC domain containing 2 |
| | KPNA4 | karyopherin subunit alpha 4 |
| | TULP4 | tubby like protein 4 |
| | CYP20A1 | cytochrome P450 family 20 subfamily A member 1 |
| | ZBTB44 | zinc finger and BTB domain containing 44 |
| | OPCML | opioid binding protein/cell adhesion molecule like |
| | TRIM33 | tripartite motif containing 33 |
| | KDSR | 3-ketodihydrosphingosine reductase |
| | ZZZ3 | zinc finger ZZ-type containing 3 |
| | SGIP1 | SH3 domain GRB2 like endophilin interacting protein 1 |
| | UBE4A | ubiquitination factor E4A |
| | ANO6 | anoctamin 6 |
| | ACVR1C | activin A receptor type 1C |
| | CHMP2B | charged multivesicular body protein 2B |
| | PTAR1 | protein prenyltransferase alpha subunit repeat containing 1 |
| | ZMYM2 | zinc finger MYM-type containing 2 |
| | ALDH6A1 | aldehyde dehydrogenase 6 family member A1 |
| | COX15 | cytochrome c oxidase assembly homolog COX15 |
| | ZNF714 | zinc finger protein 714 |
| | SEL1L | SEL1L, ERAD E3 ligase adaptor subunit |
| | NOTCH2 | notch 2 |
| | ARID2 | AT-rich interaction domain 2 |
| | FAM160B1 | family with sequence similarity 160 member B1 |
| | NCAM1 | neural cell adhesion molecule 1 |
| | ATP6V1C1 | ATPase H+ transporting V1 subunit C1 |
| | CYB5B | cytochrome b5 type B |
| | ZKSCAN8 | zinc finger with KRAB and SCAN domains 8 |
| | ARIH1 | ariadne RBR E3 ubiquitin protein ligase 1 |
| | FRS2 | fibroblast growth factor receptor substrate 2 |

| | | |
|---|---|---|
| | REV1 | REV1, DNA directed polymerase |
| | PSMD5 | proteasome 26S subunit, non-ATPase 5 |
| | ZHX1 | zinc fingers and homeoboxes 1 |
| | VPS13A | vacuolar protein sorting 13 homolog A |
| | IKZF2 | IKAROS family zinc finger 2 |
| | TLE4 | TLE family member 4, transcriptional corepressor |
| | SNX27 | sorting nexin family member 27 |
| | GPR37 | G protein-coupled receptor 37 |
| | SCN1A | sodium voltage-gated channel alpha subunit 1 |
| | SLC15A1 | solute carrier family 15 member 1 |
| | PRDM8 | PR/SET domain 8 |
| | LEF1 | lymphoid enhancer binding factor 1 |
| | MDM4 | MDM4, p53 regulator |
| | MPPED2 | metallophosphoesterase domain containing 2 |
| | ACER3 | alkaline ceramidase 3 |
| | KLF7 | Kruppel like factor 7 |
| | ADAMDEC1 | ADAM like decysin 1 |
| | GFRAL | GDNF family receptor alpha like |
| | DLGAP1 | DLG associated protein 1 |
| | HDAC2 | histone deacetylase 2 |
| | NPEPPS | aminopeptidase puromycin sensitive |
| | IQCJ | IQ motif containing J |
| | PAPOLA | poly(A) polymerase alpha |
| | CEP350 | centrosomal protein 350 |
| | PTPDC1 | protein tyrosine phosphatase domain containing 1 |
| | USP15 | ubiquitin specific peptidase 15 |
| | LILRA5 | leukocyte immunoglobulin like receptor A5 |
| | KCNJ14 | potassium voltage-gated channel subfamily J member 14 |
| | NFIA | nuclear factor I A |
| | ZDHHC21 | zinc finger DHHC-type containing 21 |
| | TMEM65 | transmembrane protein 65 |
| | DCAF10 | DDB1 and CUL4 associated factor 10 |
| | PGR | progesterone receptor |
| | UBA6 | ubiquitin like modifier activating enzyme 6 |
| | CREB5 | cAMP responsive element binding protein 5 |
| | UPF1 | UPF1, RNA helicase and ATPase |
| | PPP1R1C | protein phosphatase 1 regulatory inhibitor subunit 1C |
| | PNPO | pyridoxamine 5'-phosphate oxidase |
| | PBX1 | PBX homeobox 1 |
| | CPEB4 | cytoplasmic polyadenylation element binding protein 4 |
| | UBQLN2 | ubiquilin 2 |
| | RBMS3 | RNA binding motif single stranded interacting protein 3 |

| | | |
|---|---|---|
| | ZNF148 | zinc finger protein 148 |
| | CELF2 | CUGBP Elav-like family member 2 |
| | ATL1 | atlastin GTPase 1 |
| | CNOT8 | CCR4-NOT transcription complex subunit 8 |
| | CKAP5 | cytoskeleton associated protein 5 |
| | RABL3 | RAB, member of RAS oncogene family like 3 |
| | HEY1 | hes related family bHLH transcription factor with YRPW motif 1 |
| | NFIC | nuclear factor I C |
| | BTBD1 | BTB domain containing 1 |
| | TJP1 | tight junction protein 1 |
| | CORIN | corin, serine peptidase |
| | RBFOX2 | RNA binding fox-1 homolog 2 |
| | CEP41 | centrosomal protein 41 |
| | LAMP2 | lysosomal associated membrane protein 2 |
| | ZFP36L1 | ZFP36 ring finger protein like 1 |
| | ABHD18 | abhydrolase domain containing 18 |
| | CT55 | cancer/testis antigen 55 |
| | OTUD4 | OTU deubiquitinase 4 |
| | NUP93 | nucleoporin 93 |
| | BCLAF3 | BCLAF1 and THRAP3 family member 3 |
| | FBXW7 | F-box and WD repeat domain containing 7 |
| | ARHGAP42 | Rho GTPase activating protein 42 |
| | PPP4R2 | protein phosphatase 4 regulatory subunit 2 |
| | TRERF1 | transcriptional regulating factor 1 |
| | PPARGC1A | PPARG coactivator 1 alpha |
| | MORC3 | MORC family CW-type zinc finger 3 |
| | VGLL3 | vestigial like family member 3 |
| | ZNF483 | zinc finger protein 483 |
| | PPP4C | protein phosphatase 4 catalytic subunit |
| | BRAF | B-Raf proto-oncogene, serine/threonine kinase |
| | TBC1D4 | TBC1 domain family member 4 |
| | DYRK1A | dual specificity tyrosine phosphorylation regulated kinase 1A |
| | KCTD1 | potassium channel tetramerization domain containing 1 |
| | MAN1A1 | mannosidase alpha class 1A member 1 |
| | RHOBTB3 | Rho related BTB domain containing 3 |
| | FLVCR1 | feline leukemia virus subgroup C cellular receptor 1 |
| | CEP97 | centrosomal protein 97 |
| | LTN1 | listerin E3 ubiquitin protein ligase 1 |
| | FGL2 | fibrinogen like 2 |
| | BRWD1 | bromodomain and WD repeat domain containing 1 |
| | MYT1 | myelin transcription factor 1 |

| | | |
|---|---|---|
| | IMPAD1 | inositol monophosphatase domain containing 1 |
| | NUFIP2 | nuclear FMR1 interacting protein 2 |
| | SNTB2 | syntrophin beta 2 |
| | GPBP1L1 | GC-rich promoter binding protein 1 like 1 |
| | CAMK2D | calcium/calmodulin dependent protein kinase II delta |
| | GLI3 | GLI family zinc finger 3 |
| | MBNL2 | muscleblind like splicing regulator 2 |
| | ING2 | inhibitor of growth family member 2 |
| | C9orf3 | chromosome 9 open reading frame 3 |
| | GUCY1A2 | guanylate cyclase 1 soluble subunit alpha 2 |
| | GNAQ | G protein subunit alpha q |
| | PPP1CB | protein phosphatase 1 catalytic subunit beta |
| | CSRNP3 | cysteine and serine rich nuclear protein 3 |
| | TM9SF3 | transmembrane 9 superfamily member 3 |
| | DCAF17 | DDB1 and CUL4 associated factor 17 |
| | BCL11B | BCL11B, BAF complex component |
| | TAT | tyrosine aminotransferase |
| | SLC44A5 | solute carrier family 44 member 5 |
| | MAN2A1 | mannosidase alpha class 2A member 1 |
| | B3GALNT2 | beta-1,3-N-acetylgalactosaminyltransferase 2 |
| | PCNX3 | pecanex 3 |
| | RBPJ | recombination signal binding protein for immunoglobulin kappa J region |
| | C3orf70 | chromosome 3 open reading frame 70 |
| | ARHGAP5 | Rho GTPase activating protein 5 |
| | ZBTB10 | zinc finger and BTB domain containing 10 |
| | PURB | purine rich element binding protein B |
| | LIN7C | lin-7 homolog C, crumbs cell polarity complex component |
| | EEF2K | eukaryotic elongation factor 2 kinase |
| | EPHA5 | EPH receptor A5 |
| | BRWD3 | bromodomain and WD repeat domain containing 3 |
| | SPIN1 | spindlin 1 |
| | STC1 | stanniocalcin 1 |
| | PHACTR2 | phosphatase and actin regulator 2 |
| | XPR1 | xenotropic and polytropic retrovirus receptor 1 |
| | TRPM7 | transient receptor potential cation channel subfamily M member 7 |
| | UBE4B | ubiquitination factor E4B |
| | KATNBL1 | katanin regulatory subunit B1 like 1 |
| | SLITRK2 | SLIT and NTRK like family member 2 |
| | LRRTM3 | leucine rich repeat transmembrane neuronal 3 |
| | HIPK2 | homeodomain interacting protein kinase 2 |
| | AGFG1 | ArfGAP with FG repeats 1 |

| | | |
|---|---|---|
| | PPP3R1 | protein phosphatase 3 regulatory subunit B, alpha |
| | UCK2 | uridine-cytidine kinase 2 |
| | FBXL5 | F-box and leucine rich repeat protein 5 |
| | GRIA3 | glutamate ionotropic receptor AMPA type subunit 3 |
| | TMEM183A | transmembrane protein 183A |
| | DUSP3 | dual specificity phosphatase 3 |
| | LRCH2 | leucine rich repeats and calponin homology domain containing 2 |
| | ATP2B1 | ATPase plasma membrane Ca2+ transporting 1 |
| | FOXN2 | forkhead box N2 |
| | PTPRK | protein tyrosine phosphatase, receptor type K |
| | CCSAP | centriole, cilia and spindle associated protein |
| | APLP2 | amyloid beta precursor like protein 2 |
| | CDH11 | cadherin 11 |
| | GLDN | gliomedin |
| | RIC1 | RIC1 homolog, RAB6A GEF complex partner 1 |
| | ABCB7 | ATP binding cassette subfamily B member 7 |
| | ZNF273 | zinc finger protein 273 |
| | BDP1 | B double prime 1, subunit of RNA polymerase III transcription initiation factor IIIB |
| | DUSP19 | dual specificity phosphatase 19 |
| | HLA-DPB1 | major histocompatibility complex, class II, DP beta 1 |
| | NCAPG2 | non-SMC condensin II complex subunit G2 |
| | PRCP | prolylcarboxypeptidase |
| | MAVS | mitochondrial antiviral signaling protein |
| | C2orf49 | chromosome 2 open reading frame 49 |
| | ONECUT2 | one cut homeobox 2 |
| | UNC5C | unc-5 netrin receptor C |
| | TLK2 | tousled like kinase 2 |
| | TBCEL | tubulin folding cofactor E like |
| | STYX | serine/threonine/tyrosine interacting protein |
| | PDE6A | phosphodiesterase 6A |
| | TEAD1 | TEA domain transcription factor 1 |
| | CD200R1 | CD200 receptor 1 |
| | C19orf12 | chromosome 19 open reading frame 12 |
| | TRPS1 | transcriptional repressor GATA binding 1 |
| | PLS3 | plastin 3 |
| | L3MBTL3 | L3MBTL3, histone methyl-lysine binding protein |
| | ASB14 | ankyrin repeat and SOCS box containing 14 |
| | DCLK1 | doublecortin like kinase 1 |
| | SLC36A4 | solute carrier family 36 member 4 |
| | NR2E1 | nuclear receptor subfamily 2 group E member 1 |
| | SPATS2L | spermatogenesis associated serine rich 2 like |

| | | |
|---|---|---|
| | LSAMP | limbic system associated membrane protein |
| | ENAH | ENAH, actin regulator |
| | CNST | consortin, connexin sorting protein |
| | SOS1 | SOS Ras/Rac guanine nucleotide exchange factor 1 |
| | ATL3 | atlastin GTPase 3 |
| | MYSM1 | Myb like, SWIRM and MPN domains 1 |
| | PNMA2 | PNMA family member 2 |
| | ALYREF | Aly/REF export factor |
| | SESTD1 | SEC14 and spectrin domain containing 1 |
| | RNF11 | ring finger protein 11 |
| | THRAP3 | thyroid hormone receptor associated protein 3 |
| | HOXD9 | homeobox D9 |
| | LRRN1 | leucine rich repeat neuronal 1 |
| | PCNP | PEST proteolytic signal containing nuclear protein |
| | SDE2 | SDE2 telomere maintenance homolog |
| | SHPRH | SNF2 histone linker PHD RING helicase |
| | VCPKMT | valosin containing protein lysine methyltransferase |
| | EPM2AIP1 | EPM2A interacting protein 1 |
| | PKP4 | plakophilin 4 |
| | GUCY1A1 | guanylate cyclase 1 soluble subunit alpha 1 |
| | CAND1 | cullin associated and neddylation dissociated 1 |
| | UBE2K | ubiquitin conjugating enzyme E2 K |
| | ZNF704 | zinc finger protein 704 |
| | LRRC39 | leucine rich repeat containing 39 |
| | AFF4 | AF4/FMR2 family member 4 |
| | SAMD12 | sterile alpha motif domain containing 12 |
| | BTBD11 | BTB domain containing 11 |
| | ANKRD28 | ankyrin repeat domain 28 |
| | PHF12 | PHD finger protein 12 |
| | SF3B3 | splicing factor 3b subunit 3 |
| | PRDM15 | PR/SET domain 15 |
| | APELA | apelin receptor early endogenous ligand |
| | ACOT13 | acyl-CoA thioesterase 13 |
| | RASGEF1B | RasGEF domain family member 1B |
| | SRGAP2C | SLIT-ROBO Rho GTPase activating protein 2C |
| | HNRNPU | heterogeneous nuclear ribonucleoprotein U |
| | ZMAT4 | zinc finger matrin-type 4 |
| | ATXN1 | ataxin 1 |
| | MAP7D3 | MAP7 domain containing 3 |
| | LIN9 | lin-9 DREAM MuvB core complex component |
| | HSPH1 | heat shock protein family H (Hsp110) member 1 |
| | ZC3H12C | zinc finger CCCH-type containing 12C |

| | | |
|---|---|---|
| | ZNF501 | zinc finger protein 501 |
| | TDGF1 | teratocarcinoma-derived growth factor 1 |
| | KCNJ6 | potassium voltage-gated channel subfamily J member 6 |
| | GLIPR1L2 | GLIPR1 like 2 |
| | NIPAL1 | NIPA like domain containing 1 |
| | C18orf25 | chromosome 18 open reading frame 25 |
| | PGM2L1 | phosphoglucomutase 2 like 1 |
| | SULT1E1 | sulfotransferase family 1E member 1 |
| | SUCLA2 | succinate-CoA ligase ADP-forming beta subunit |
| | TSHZ1 | teashirt zinc finger homeobox 1 |
| | CDCP1 | CUB domain containing protein 1 |
| | TMEM161B | transmembrane protein 161B |
| | CDH12 | cadherin 12 |
| | ERCC6 | ERCC excision repair 6, chromatin remodeling factor |
| | NCBP3 | nuclear cap binding subunit 3 |
| | CDH20 | cadherin 20 |
| | MAP4 | microtubule associated protein 4 |
| | NRXN1 | neurexin 1 |
| | FBXO21 | F-box protein 21 |
| | GMCL1 | germ cell-less, spermatogenesis associated 1 |
| | CSNK1G3 | casein kinase 1 gamma 3 |
| | FAXC | failed axon connections homolog |
| | TRIM71 | tripartite motif containing 71 |
| | CTSC | cathepsin C |
| | SLC48A1 | solute carrier family 48 member 1 |
| | C5orf24 | chromosome 5 open reading frame 24 |
| | SDK1 | sidekick cell adhesion molecule 1 |
| | CLSPN | claspin |
| | DENND1B | DENN domain containing 1B |
| | PARG | poly(ADP-ribose) glycohydrolase |
| | SNRPF | small nuclear ribonucleoprotein polypeptide F |
| | HOMER1 | homer scaffold protein 1 |
| | BMPR2 | bone morphogenetic protein receptor type 2 |
| | PRTG | protogenin |
| | IL33 | interleukin 33 |
| | AMER2 | APC membrane recruitment protein 2 |
| | SNRNP40 | small nuclear ribonucleoprotein U5 subunit 40 |
| | TRUB1 | TruB pseudouridine synthase family member 1 |
| | SPAST | spastin |
| | COQ7 | coenzyme Q7, hydroxylase |
| | DVL3 | dishevelled segment polarity protein 3 |
| | NAA30 | N(alpha)-acetyltransferase 30, NatC catalytic subunit |

| | | |
|---|---|---|
| | SOX11 | SRY-box 11 |
| | SHANK2 | SH3 and multiple ankyrin repeat domains 2 |
| | FSBP | fibrinogen silencer binding protein |
| | CPLX2 | complexin 2 |
| | KLHL24 | kelch like family member 24 |
| | ZYG11B | zyg-11 family member B, cell cycle regulator |
| | RORA | RAR related orphan receptor A |
| | CLCN3 | chloride voltage-gated channel 3 |
| | LUC7L3 | LUC7 like 3 pre-mRNA splicing factor |
| | GABPB1 | GA binding protein transcription factor subunit beta 1 |
| | CDC27 | cell division cycle 27 |
| | ABCA5 | ATP binding cassette subfamily A member 5 |
| | FXR1 | FMR1 autosomal homolog 1 |
| | WNK3 | WNK lysine deficient protein kinase 3 |
| | QRICH1 | glutamine rich 1 |
| | ATRX | ATRX, chromatin remodeler |
| | LMO3 | LIM domain only 3 |
| | CELF1 | CUGBP Elav-like family member 1 |
| | PRKACA | protein kinase cAMP-activated catalytic subunit alpha |
| | COL25A1 | collagen type XXV alpha 1 chain |
| | UBR2 | ubiquitin protein ligase E3 component n-recognin 2 |
| | NKX3-1 | NK3 homeobox 1 |
| | SIX4 | SIX homeobox 4 |
| | MAGI2 | membrane associated guanylate kinase, WW and PDZ domain containing 2 |
| | SOX4 | SRY-box 4 |
| | TRDMT1 | tRNA aspartic acid methyltransferase 1 |
| | TGFBR2 | transforming growth factor beta receptor 2 |
| | FBXO42 | F-box protein 42 |
| | BCOR | BCL6 corepressor |
| | SERPINB1 | serpin family B member 1 |
| | ADAM22 | ADAM metallopeptidase domain 22 |
| | VCAN | versican |
| | RBM48 | RNA binding motif protein 48 |
| | SH3BP2 | SH3 domain binding protein 2 |
| | FGF12 | fibroblast growth factor 12 |
| | SERTM1 | serine rich and transmembrane domain containing 1 |
| | SLC25A37 | solute carrier family 25 member 37 |
| | P4HA1 | prolyl 4-hydroxylase subunit alpha 1 |
| | AP1S3 | adaptor related protein complex 1 subunit sigma 3 |
| | MKLN1 | muskelin 1 |
| | RNF19A | ring finger protein 19A, RBR E3 ubiquitin protein ligase |

| | | |
|---|---|---|
| | RERG | RAS like estrogen regulated growth inhibitor |
| | TP53 | tumor protein p53 |
| | FCF1 | FCF1, rRNA-processing protein |
| | THEM4 | thioesterase superfamily member 4 |
| | CFAP53 | cilia and flagella associated protein 53 |
| | ZBTB18 | zinc finger and BTB domain containing 18 |
| | PCDHB11 | protocadherin beta 11 |
| | DAPK1 | death associated protein kinase 1 |
| | TMEM135 | transmembrane protein 135 |
| | XPO4 | exportin 4 |
| | OR11A1 | olfactory receptor family 11 subfamily A member 1 |
| | SOGA1 | suppressor of glucose, autophagy associated 1 |
| | LBR | lamin B receptor |
| | MSANTD3 | Myb/SANT DNA binding domain containing 3 |
| | ABHD13 | abhydrolase domain containing 13 |
| | VEZT | vezatin, adherens junctions transmembrane protein |
| | ERP44 | endoplasmic reticulum protein 44 |
| | ACBD7 | acyl-CoA binding domain containing 7 |
| | AKAP11 | A-kinase anchoring protein 11 |
| | CLINT1 | clathrin interactor 1 |
| | TMEM45A | transmembrane protein 45A |
| | HS3ST3B1 | heparan sulfate-glucosamine 3-sulfotransferase 3B1 |
| | SMG1 | SMG1, nonsense mediated mRNA decay associated PI3K related kinase |
| | ATF2 | activating transcription factor 2 |
| | MYNN | myoneurin |
| | HECTD2 | HECT domain E3 ubiquitin protein ligase 2 |
| | TMED2 | transmembrane p24 trafficking protein 2 |
| | SACM1L | SAC1 like phosphatidylinositide phosphatase |
| | HNRNPA3 | heterogeneous nuclear ribonucleoprotein A3 |
| | HGF | hepatocyte growth factor |
| | FAM20B | FAM20B, glycosaminoglycan xylosylkinase |
| | TMEM74 | transmembrane protein 74 |
| | ZNF814 | zinc finger protein 814 |
| | C1orf115 | chromosome 1 open reading frame 115 |
| | TSHZ3 | teashirt zinc finger homeobox 3 |
| | NLGN1 | neuroligin 1 |
| | GOLT1B | golgi transport 1B |
| | PABPC4L | poly(A) binding protein cytoplasmic 4 like |
| | SLC27A6 | solute carrier family 27 member 6 |
| | NAA50 | N(alpha)-acetyltransferase 50, NatE catalytic subunit |
| | RNF125 | ring finger protein 125 |

| | | |
|---|---|---|
| | SGPL1 | sphingosine-1-phosphate lyase 1 |
| | SAMHD1 | SAM and HD domain containing deoxynucleoside triphosphate triphosphohydrolase 1 |
| | E2F7 | E2F transcription factor 7 |
| | RNF144A | ring finger protein 144A |
| | SH3GLB1 | SH3 domain containing GRB2 like, endophilin B1 |
| | CHD2 | chromodomain helicase DNA binding protein 2 |
| | ID4 | inhibitor of DNA binding 4, HLH protein |
| | RALGAPB | Ral GTPase activating protein non-catalytic beta subunit |
| | TBC1D9 | TBC1 domain family member 9 |
| | KPNA6 | karyopherin subunit alpha 6 |
| | IRF2BP2 | interferon regulatory factor 2 binding protein 2 |
| | PPP1R9A | protein phosphatase 1 regulatory subunit 9A |
| | USP37 | ubiquitin specific peptidase 37 |
| | C18orf54 | chromosome 18 open reading frame 54 |
| | PTPN11 | protein tyrosine phosphatase, non-receptor type 11 |
| | FAM126B | family with sequence similarity 126 member B |
| | DNAJA2 | DnaJ heat shock protein family (Hsp40) member A2 |
| | GRIP1 | glutamate receptor interacting protein 1 |
| | AEBP2 | AE binding protein 2 |
| | FMR1 | fragile X mental retardation 1 |
| | TTPAL | alpha tocopherol transfer protein like |
| | ESCO2 | establishment of sister chromatid cohesion N-acetyltransferase 2 |
| | FRMD4B | FERM domain containing 4B |
| | WEE1 | WEE1 G2 checkpoint kinase |
| | PIGA | phosphatidylinositol glycan anchor biosynthesis class A |
| | GOLIM4 | golgi integral membrane protein 4 |
| | GDAP2 | ganglioside induced differentiation associated protein 2 |
| | MN1 | MN1 proto-oncogene, transcriptional regulator |
| | ICK | intestinal cell kinase |
| | CEBPZOS | CEBPZ opposite strand |
| | RBMS2 | RNA binding motif single stranded interacting protein 2 |
| | ELAVL4 | ELAV like RNA binding protein 4 |
| | STAM | signal transducing adaptor molecule |
| | ZNHIT6 | zinc finger HIT-type containing 6 |
| | MITF | melanocyte inducing transcription factor |
| | GTF2B | general transcription factor IIB |
| | CCDC158 | coiled-coil domain containing 158 |
| | FUBP3 | far upstream element binding protein 3 |
| | RANBP3L | RAN binding protein 3 like |
| | FEZF1 | FEZ family zinc finger 1 |
| | DMD | dystrophin |

| | | |
|---|---|---|
| | RBM27 | RNA binding motif protein 27 |
| | SETBP1 | SET binding protein 1 |
| | LIFR | LIF receptor alpha |
| | ATP8B2 | ATPase phospholipid transporting 8B2 |
| | S1PR5 | sphingosine-1-phosphate receptor 5 |
| | DPP9 | dipeptidyl peptidase 9 |
| | ZDHHC14 | zinc finger DHHC-type containing 14 |
| | CDC42SE2 | CDC42 small effector 2 |
| | ZNF354C | zinc finger protein 354C |
| | ALX1 | ALX homeobox 1 |
| | TFCP2L1 | transcription factor CP2 like 1 |
| | ZNF813 | zinc finger protein 813 |
| | FBXL18 | F-box and leucine rich repeat protein 18 |
| | EIF2AK2 | eukaryotic translation initiation factor 2 alpha kinase 2 |
| | GPC6 | glypican 6 |
| | RBM26 | RNA binding motif protein 26 |
| | USP38 | ubiquitin specific peptidase 38 |
| | PLEKHB2 | pleckstrin homology domain containing B2 |
| | GCFC2 | GC-rich sequence DNA-binding factor 2 |
| | SNRPD1 | small nuclear ribonucleoprotein D1 polypeptide |
| | STK26 | serine/threonine kinase 26 |
| | QKI | QKI, KH domain containing RNA binding |
| | WASL | Wiskott-Aldrich syndrome like |
| | CASK | calcium/calmodulin dependent serine protein kinase |
| | TYW1B | tRNA-yW synthesizing protein 1 homolog B |
| | AK4 | adenylate kinase 4 |
| | EPHB1 | EPH receptor B1 |
| | RAB3C | RAB3C, member RAS oncogene family |
| | GPATCH2L | G-patch domain containing 2 like |
| | KCNMA1 | potassium calcium-activated channel subfamily M alpha 1 |
| | SPG21 | SPG21, maspardin |
| | DNAL1 | dynein axonemal light chain 1 |
| | PITPNA | phosphatidylinositol transfer protein alpha |
| | ANKRD42 | ankyrin repeat domain 42 |
| | KCNK2 | potassium two pore domain channel subfamily K member 2 |
| | HELB | DNA helicase B |
| | DPH6 | diphthamine biosynthesis 6 |
| | ATP2C1 | ATPase secretory pathway Ca2+ transporting 1 |
| | ZBTB39 | zinc finger and BTB domain containing 39 |
| | LURAP1L | leucine rich adaptor protein 1 like |
| | BEND4 | BEN domain containing 4 |
| | RAB22A | RAB22A, member RAS oncogene family |

| | | |
|---|---|---|
| | BOLL | boule homolog, RNA binding protein |
| | HAUS6 | HAUS augmin like complex subunit 6 |
| | PSMD11 | proteasome 26S subunit, non-ATPase 11 |
| | HSD3B7 | hydroxy-delta-5-steroid dehydrogenase, 3 beta- and steroid delta-isomerase 7 |
| | IRF6 | interferon regulatory factor 6 |
| | RSL24D1 | ribosomal L24 domain containing 1 |
| | RAB33B | RAB33B, member RAS oncogene family |
| | SREK1 | splicing regulatory glutamic acid and lysine rich protein 1 |
| | PRKAR2A | protein kinase cAMP-dependent type II regulatory subunit alpha |
| | ST7 | suppression of tumorigenicity 7 |
| | MCF2 | MCF.2 cell line derived transforming sequence |
| | BTF3L4 | basic transcription factor 3 like 4 |
| | TRAF3IP1 | TRAF3 interacting protein 1 |
| | IP6K2 | inositol hexakisphosphate kinase 2 |
| | KMT2E | lysine methyltransferase 2E |
| | TMED10 | transmembrane p24 trafficking protein 10 |
| | ABLIM1 | actin binding LIM protein 1 |
| | APAF1 | apoptotic peptidase activating factor 1 |
| | UTP25 | UTP25, small subunit processor component |
| | SDR16C5 | short chain dehydrogenase/reductase family 16C member 5 |
| | TSHZ2 | teashirt zinc finger homeobox 2 |
| | C17orf75 | chromosome 17 open reading frame 75 |
| | MED23 | mediator complex subunit 23 |
| | AHCYL2 | adenosylhomocysteinase like 2 |
| | BNC2 | basonuclin 2 |
| | CCAR1 | cell division cycle and apoptosis regulator 1 |
| | PPP1R14C | protein phosphatase 1 regulatory inhibitor subunit 14C |
| | KIAA0895 | KIAA0895 |
| | BACH2 | BTB domain and CNC homolog 2 |
| | DLG1 | discs large MAGUK scaffold protein 1 |
| | RNF170 | ring finger protein 170 |
| | POU2F3 | POU class 2 homeobox 3 |
| | ST3GAL2 | ST3 beta-galactoside alpha-2,3-sialyltransferase 2 |
| | CPB1 | carboxypeptidase B1 |
| | EPS15 | epidermal growth factor receptor pathway substrate 15 |
| | CHD1 | chromodomain helicase DNA binding protein 1 |
| | TRIM2 | tripartite motif containing 2 |
| | ANKS1B | ankyrin repeat and sterile alpha motif domain containing 1B |
| | ZNF391 | zinc finger protein 391 |
| | SYNJ2BP | synaptojanin 2 binding protein |
| | SMPDL3A | sphingomyelin phosphodiesterase acid like 3A |

| | | |
|---|---|---|
| | SOGA3 | SOGA family member 3 |
| | XYLT2 | xylosyltransferase 2 |
| | MACF1 | microtubule-actin crosslinking factor 1 |
| | GRIK1 | glutamate ionotropic receptor kainate type subunit 1 |
| | SEPTIN7 | septin 7 |
| | KIF1B | kinesin family member 1B |
| | TGDS | TDP-glucose 4,6-dehydratase |
| | SLC30A7 | solute carrier family 30 member 7 |
| | AKAP5 | A-kinase anchoring protein 5 |
| | SLC7A2 | solute carrier family 7 member 2 |
| | REEP1 | receptor accessory protein 1 |
| | SCLT1 | sodium channel and clathrin linker 1 |
| | NKAIN2 | sodium/potassium transporting ATPase interacting 2 |
| | ASXL3 | ASXL transcriptional regulator 3 |
| | NUDT11 | nudix hydrolase 11 |
| | IDH1 | isocitrate dehydrogenase (NADP(+)) 1, cytosolic |
| | ESR1 | estrogen receptor 1 |
| | FAM118A | family with sequence similarity 118 member A |
| | JCAD | junctional cadherin 5 associated |
| | MGP | matrix Gla protein |
| | GPR180 | G protein-coupled receptor 180 |
| | PPP1R15B | protein phosphatase 1 regulatory subunit 15B |
| | EEF1AKMT3 | EEF1A lysine methyltransferase 3 |
| | POLI | DNA polymerase iota |
| | MTF2 | metal response element binding transcription factor 2 |
| | PPP1R3B | protein phosphatase 1 regulatory subunit 3B |
| | KLHL32 | kelch like family member 32 |
| | KIAA0408 | KIAA0408 |
| | PDE12 | phosphodiesterase 12 |
| | CHIC1 | cysteine rich hydrophobic domain 1 |
| | ZC3H4 | zinc finger CCCH-type containing 4 |
| | ZFP36L2 | ZFP36 ring finger protein like 2 |
| | ZFP42 | ZFP42 zinc finger protein |
| | NHLRC2 | NHL repeat containing 2 |
| | EIF2S1 | eukaryotic translation initiation factor 2 subunit alpha |
| | UBN2 | ubinuclein 2 |
| | SNRNP27 | small nuclear ribonucleoprotein U4/U6.U5 subunit 27 |
| | RNF2 | ring finger protein 2 |
| | TENM1 | teneurin transmembrane protein 1 |
| | AKAIN1 | A-kinase anchor inhibitor 1 |
| | PLIN1 | perilipin 1 |
| | JMY | junction mediating and regulatory protein, p53 cofactor |

| | | |
|---|---|---|
| | FOXN3 | forkhead box N3 |
| | CREB3L2 | cAMP responsive element binding protein 3 like 2 |
| | OSBPL11 | oxysterol binding protein like 11 |
| | MOB3B | MOB kinase activator 3B |
| | DIPK2A | divergent protein kinase domain 2A |
| | PPM1A | protein phosphatase, Mg2+/Mn2+ dependent 1A |
| | TWSG1 | twisted gastrulation BMP signaling modulator 1 |
| | PM20D2 | peptidase M20 domain containing 2 |
| | MMGT1 | membrane magnesium transporter 1 |
| | PPM1K | protein phosphatase, Mg2+/Mn2+ dependent 1K |
| | LIN54 | lin-54 DREAM MuvB core complex component |
| | THUMPD1 | THUMP domain containing 1 |
| | EXOC8 | exocyst complex component 8 |
| | SMU1 | SMU1, DNA replication regulator and spliceosomal factor |
| | MARCHF7 | membrane associated ring-CH-type finger 7 |
| | EXOSC6 | exosome component 6 |
| | TTC26 | tetratricopeptide repeat domain 26 |
| | JPH1 | junctophilin 1 |
| | UBL3 | ubiquitin like 3 |
| | CREB1 | cAMP responsive element binding protein 1 |
| | NRAS | NRAS proto-oncogene, GTPase |
| | ARRDC3 | arrestin domain containing 3 |
| | SELL | selectin L |
| | RBM8A | RNA binding motif protein 8A |
| | GCM1 | glial cells missing transcription factor 1 |
| | SPOCK3 | SPARC (osteonectin), cwcv and kazal like domains proteoglycan 3 |
| | ERAP1 | endoplasmic reticulum aminopeptidase 1 |
| | PIK3CG | phosphatidylinositol-4,5-bisphosphate 3-kinase catalytic subunit gamma |
| | TSPYL6 | TSPY like 6 |
| | ATRAID | all-trans retinoic acid induced differentiation factor |
| | AS3MT | arsenite methyltransferase |
| | ANKRD12 | ankyrin repeat domain 12 |
| | PDE8A | phosphodiesterase 8A |
| | C16orf74 | chromosome 16 open reading frame 74 |
| | AKAP2 | A-kinase anchoring protein 2 |
| | EDEM1 | ER degradation enhancing alpha-mannosidase like protein 1 |
| | TEDDM1 | transmembrane epididymal protein 1 |
| | TRIM5 | tripartite motif containing 5 |
| | EIF5A2 | eukaryotic translation initiation factor 5A2 |
| | SNX18 | sorting nexin 18 |
| | ERGIC1 | endoplasmic reticulum-golgi intermediate compartment 1 |

| | | |
|---|---|---|
| | MMAA | metabolism of cobalamin associated A |
| | RAB2B | RAB2B, member RAS oncogene family |
| | MBNL3 | muscleblind like splicing regulator 3 |
| | SUCO | SUN domain containing ossification factor |
| | TAS2R20 | taste 2 receptor member 20 |
| | PBRM1 | polybromo 1 |
| | MAP3K2 | mitogen-activated protein kinase kinase kinase 2 |
| | MYO6 | myosin VI |
| | LRP11 | LDL receptor related protein 11 |
| | PELI1 | pellino E3 ubiquitin protein ligase 1 |
| | PYGO1 | pygopus family PHD finger 1 |
| | FAM107B | family with sequence similarity 107 member B |
| | CAPRIN1 | cell cycle associated protein 1 |
| | RELA | RELA proto-oncogene, NF-kB subunit |
| | KIAA1841 | KIAA1841 |
| | MID1 | midline 1 |
| | NUP35 | nucleoporin 35 |
| | VAMP2 | vesicle associated membrane protein 2 |
| | FOXK1 | forkhead box K1 |
| | VAMP4 | vesicle associated membrane protein 4 |
| | FGF14 | fibroblast growth factor 14 |
| | RBM17 | RNA binding motif protein 17 |
| | KIAA0930 | KIAA0930 |
| | DPYSL2 | dihydropyrimidinase like 2 |
| | GOLGA7 | golgin A7 |
| | IDE | insulin degrading enzyme |
| | FERMT1 | fermitin family member 1 |
| | PXDN | peroxidasin |
| | PLA2G4A | phospholipase A2 group IVA |
| | CAMK2A | calcium/calmodulin dependent protein kinase II alpha |
| | TRIM7 | tripartite motif containing 7 |
| | DOCK4 | dedicator of cytokinesis 4 |
| | NRG1 | neuregulin 1 |
| | SMIM15 | small integral membrane protein 15 |
| | MED21 | mediator complex subunit 21 |
| | ZNF736 | zinc finger protein 736 |
| | ESRRG | estrogen related receptor gamma |
| | EHD1 | EH domain containing 1 |
| | STARD3NL | STARD3 N-terminal like |
| | LIMD1 | LIM domains containing 1 |
| | HYKK | hydroxylysine kinase |
| | RHOBTB1 | Rho related BTB domain containing 1 |

| | | |
|---|---|---|
| | OGFRL1 | opioid growth factor receptor like 1 |
| | BAZ2B | bromodomain adjacent to zinc finger domain 2B |
| | SYAP1 | synapse associated protein 1 |
| | WNT5A | Wnt family member 5A |
| | BMP3 | bone morphogenetic protein 3 |
| | LDB2 | LIM domain binding 2 |
| | NUP155 | nucleoporin 155 |
| | HAS2 | hyaluronan synthase 2 |
| | TPRG1 | tumor protein p63 regulated 1 |
| | TMEM263 | transmembrane protein 263 |
| | SCUBE3 | signal peptide, CUB domain and EGF like domain containing 3 |
| | CAPZA1 | capping actin protein of muscle Z-line subunit alpha 1 |
| | VMA21 | VMA21, vacuolar ATPase assembly factor |
| | CHMP7 | charged multivesicular body protein 7 |
| | FAM120A | family with sequence similarity 120A |
| | NOL11 | nucleolar protein 11 |
| | CORO2B | coronin 2B |
| | CSDE1 | cold shock domain containing E1 |
| | SPRED1 | sprouty related EVH1 domain containing 1 |
| | GLS | glutaminase |
| | REXO2 | RNA exonuclease 2 |
| | ELAVL1 | ELAV like RNA binding protein 1 |
| | RNF217 | ring finger protein 217 |
| | ANKUB1 | ankyrin repeat and ubiquitin domain containing 1 |
| | HOXA2 | homeobox A2 |
| | SPRYD7 | SPRY domain containing 7 |
| | RAB8A | RAB8A, member RAS oncogene family |
| | OSGIN2 | oxidative stress induced growth inhibitor family member 2 |
| | COL22A1 | collagen type XXII alpha 1 chain |
| | DSCAM | DS cell adhesion molecule |
| | ST8SIA1 | ST8 alpha-N-acetyl-neuraminide alpha-2,8-sialyltransferase 1 |
| | CYB5R4 | cytochrome b5 reductase 4 |
| | BPTF | bromodomain PHD finger transcription factor |
| | SEC22C | SEC22 homolog C, vesicle trafficking protein |
| | WAC | WW domain containing adaptor with coiled-coil |
| | NDN | necdin, MAGE family member |
| | XIRP2 | xin actin binding repeat containing 2 |
| | LBH | limb bud and heart development |
| | FANCF | FA complementation group F |
| | MYOCD | myocardin |
| | TYW1 | tRNA-yW synthesizing protein 1 homolog |
| | RSPO4 | R-spondin 4 |

| | | |
|---|---|---|
| | RBL1 | RB transcriptional corepressor like 1 |
| | RBFOX1 | RNA binding fox-1 homolog 1 |
| | SUMO1 | small ubiquitin-like modifier 1 |
| | ASZ1 | ankyrin repeat, SAM and basic leucine zipper domain containing 1 |
| | MRPL19 | mitochondrial ribosomal protein L19 |
| | ZNF224 | zinc finger protein 224 |
| | ZEB1 | zinc finger E-box binding homeobox 1 |
| | GTPBP10 | GTP binding protein 10 |
| | BRPF3 | bromodomain and PHD finger containing 3 |
| | SMIM17 | small integral membrane protein 17 |
| | DERL2 | derlin 2 |
| | PROX1 | prospero homeobox 1 |
| | TIGD6 | tigger transposable element derived 6 |
| | AMER1 | APC membrane recruitment protein 1 |
| | PTPRJ | protein tyrosine phosphatase, receptor type J |
| | ZNF493 | zinc finger protein 493 |
| | PAX5 | paired box 5 |
| | DENND4A | DENN domain containing 4A |
| | BUD13 | BUD13 homolog |
| | UBE3A | ubiquitin protein ligase E3A |
| | PPP3CB | protein phosphatase 3 catalytic subunit beta |
| | LARP4B | La ribonucleoprotein domain family member 4B |
| | TRIM13 | tripartite motif containing 13 |
| | PHC3 | polyhomeotic homolog 3 |
| | FBXO40 | F-box protein 40 |
| | MYBL1 | MYB proto-oncogene like 1 |
| | PDE7B | phosphodiesterase 7B |
| | OSBPL3 | oxysterol binding protein like 3 |
| | NR2C2 | nuclear receptor subfamily 2 group C member 2 |
| | REL | REL proto-oncogene, NF-kB subunit |
| | ATAD1 | ATPase family, AAA domain containing 1 |
| | AFTPH | aftiphilin |
| | MARF1 | meiosis regulator and mRNA stability factor 1 |
| | TCF20 | transcription factor 20 |
| | USF1 | upstream transcription factor 1 |
| | SMG7 | SMG7, nonsense mediated mRNA decay factor |
| | DUSP10 | dual specificity phosphatase 10 |
| | LRRCC1 | leucine rich repeat and coiled-coil centrosomal protein 1 |
| | OPA1 | OPA1, mitochondrial dynamin like GTPase |
| | ZMYM4 | zinc finger MYM-type containing 4 |
| | USF3 | upstream transcription factor family member 3 |

| | | |
|---|---|---|
| | SLC43A2 | solute carrier family 43 member 2 |
| | MOB1A | MOB kinase activator 1A |
| | ARHGEF9 | Cdc42 guanine nucleotide exchange factor 9 |
| | TAL1 | TAL bHLH transcription factor 1, erythroid differentiation factor |
| | SBSPON | somatomedin B and thrombospondin type 1 domain containing |
| | MED13 | mediator complex subunit 13 |
| | CCDC88A | coiled-coil domain containing 88A |
| | IL26 | interleukin 26 |
| | FNIP2 | folliculin interacting protein 2 |
| | ENDOD1 | endonuclease domain containing 1 |
| | NME7 | NME/NM23 family member 7 |
| | SSBP2 | single stranded DNA binding protein 2 |
| | PDCD10 | programmed cell death 10 |
| | CAMSAP2 | calmodulin regulated spectrin associated protein family member 2 |
| | DYNC1I2 | dynein cytoplasmic 1 intermediate chain 2 |
| | NCOA2 | nuclear receptor coactivator 2 |
| | ASB5 | ankyrin repeat and SOCS box containing 5 |
| | CAMLG | calcium modulating ligand |
| | FOXP1 | forkhead box P1 |
| | DGKH | diacylglycerol kinase eta |
| | MED13L | mediator complex subunit 13 like |
| | ALDH1A2 | aldehyde dehydrogenase 1 family member A2 |
| | CAMK2N1 | calcium/calmodulin dependent protein kinase II inhibitor 1 |
| | OSMR | oncostatin M receptor |
| | TAOK1 | TAO kinase 1 |
| | PIK3R1 | phosphoinositide-3-kinase regulatory subunit 1 |
| | GTF3C3 | general transcription factor IIIC subunit 3 |
| | TERF2 | telomeric repeat binding factor 2 |
| | VNN1 | vanin 1 |
| | C1QTNF7 | C1q and TNF related 7 |
| | SEMA6D | semaphorin 6D |
| | SNX13 | sorting nexin 13 |
| | IDUA | iduronidase, alpha-L- |
| | PRPF4B | pre-mRNA processing factor 4B |
| | CAPZA2 | capping actin protein of muscle Z-line subunit alpha 2 |
| | SLC51B | solute carrier family 51 beta subunit |
| | TLR10 | toll like receptor 10 |
| | CLSTN1 | calsyntenin 1 |
| | MYLIP | myosin regulatory light chain interacting protein |
| | SATB1 | SATB homeobox 1 |
| | SERPINB9 | serpin family B member 9 |

| | | |
|---|---|---|
| | ZBTB8A | zinc finger and BTB domain containing 8A |
| | GIT2 | GIT ArfGAP 2 |
| | ARID1B | AT-rich interaction domain 1B |
| | ELL2 | elongation factor for RNA polymerase II 2 |
| | PIP4P2 | phosphatidylinositol-4,5-bisphosphate 4-phosphatase 2 |
| | KIF14 | kinesin family member 14 |
| | ANO2 | anoctamin 2 |
| | AREL1 | apoptosis resistant E3 ubiquitin protein ligase 1 |
| | EIF4E | eukaryotic translation initiation factor 4E |
| | IFT80 | intraflagellar transport 80 |
| | SLX4 | SLX4 structure-specific endonuclease subunit |
| | SPTSSB | serine palmitoyltransferase small subunit B |
| | PCDH19 | protocadherin 19 |
| | BARD1 | BRCA1 associated RING domain 1 |
| | WDR33 | WD repeat domain 33 |
| | ARMC1 | armadillo repeat containing 1 |
| | XRN2 | 5'-3' exoribonuclease 2 |
| | TCEA1 | transcription elongation factor A1 |
| | PRKAA2 | protein kinase AMP-activated catalytic subunit alpha 2 |
| | TMEM183B | transmembrane protein 183B |
| | FAM210A | family with sequence similarity 210 member A |
| | ZNF22 | zinc finger protein 22 |
| | DOCK9 | dedicator of cytokinesis 9 |
| | KLHL26 | kelch like family member 26 |
| | KAT6B | lysine acetyltransferase 6B |
| | LRP12 | LDL receptor related protein 12 |
| | BBS7 | Bardet-Biedl syndrome 7 |
| | PRKG2 | protein kinase cGMP-dependent 2 |
| | KCNT2 | potassium sodium-activated channel subfamily T member 2 |
| | SLC6A14 | solute carrier family 6 member 14 |
| | DACH1 | dachshund family transcription factor 1 |
| | FKTN | fukutin |
| | ERI1 | exoribonuclease 1 |
| | FBXO28 | F-box protein 28 |
| | AAK1 | AP2 associated kinase 1 |
| | RASA1 | RAS p21 protein activator 1 |
| | IYD | iodotyrosine deiodinase |
| | SFPQ | splicing factor proline and glutamine rich |
| | KIF5B | kinesin family member 5B |
| | LRIG1 | leucine rich repeats and immunoglobulin like domains 1 |
| | EIF4EBP2 | eukaryotic translation initiation factor 4E binding protein 2 |
| | ZNF250 | zinc finger protein 250 |

| | | |
|---|---|---|
| | TRIM10 | tripartite motif containing 10 |
| | DR1 | down-regulator of transcription 1 |
| | MID1IP1 | MID1 interacting protein 1 |
| | CLOCK | clock circadian regulator |
| | MIDN | midnolin |
| | CPSF7 | cleavage and polyadenylation specific factor 7 |
| | MIGA1 | mitoguardin 1 |
| | LDHAL6A | lactate dehydrogenase A like 6A |
| | BCLAF1 | BCL2 associated transcription factor 1 |
| | SLC17A6 | solute carrier family 17 member 6 |
| | FAM126A | family with sequence similarity 126 member A |
| | GLUL | glutamate-ammonia ligase |
| | LMOD2 | leiomodin 2 |
| | MAGI3 | membrane associated guanylate kinase, WW and PDZ domain containing 3 |
| | TLK1 | tousled like kinase 1 |
| | IGFBP7 | insulin like growth factor binding protein 7 |
| | YLPM1 | YLP motif containing 1 |
| | ARHGEF38 | Rho guanine nucleotide exchange factor 38 |
| | CCBE1 | collagen and calcium binding EGF domains 1 |
| | C17orf102 | chromosome 17 open reading frame 102 |
| | SHISA9 | shisa family member 9 |
| | VPS35L | VPS35 endosomal protein sorting factor like |
| | API5 | apoptosis inhibitor 5 |
| | HSPE1 | heat shock protein family E (Hsp10) member 1 |
| | IPO7 | importin 7 |
| | QTRT2 | queuine tRNA-ribosyltransferase accessory subunit 2 |
| | ARAP2 | ArfGAP with RhoGAP domain, ankyrin repeat and PH domain 2 |
| | STK38L | serine/threonine kinase 38 like |
| | RETREG3 | reticulophagy regulator family member 3 |
| | TMX3 | thioredoxin related transmembrane protein 3 |
| | NEUROD1 | neuronal differentiation 1 |
| | SLC12A6 | solute carrier family 12 member 6 |
| | PHF21B | PHD finger protein 21B |
| | ALDH1B1 | aldehyde dehydrogenase 1 family member B1 |
| | NR1D2 | nuclear receptor subfamily 1 group D member 2 |
| | USP31 | ubiquitin specific peptidase 31 |
| | KIF13A | kinesin family member 13A |
| | WDR92 | WD repeat domain 92 |
| | PCDHB16 | protocadherin beta 16 |
| | FAM120C | family with sequence similarity 120C |
| | PAX6 | paired box 6 |

| | | |
|---|---|---|
| | TINAGL1 | tubulointerstitial nephritis antigen like 1 |
| | GNG12 | G protein subunit gamma 12 |
| | KAZN | kazrin, periplakin interacting protein |
| | AHCYL1 | adenosylhomocysteinase like 1 |
| | ENTPD7 | ectonucleoside triphosphate diphosphohydrolase 7 |
| | PEAK3 | PEAK family member 3 |
| | TGIF2 | TGFB induced factor homeobox 2 |
| | FREM2 | FRAS1 related extracellular matrix protein 2 |
| | CBLB | Cbl proto-oncogene B |
| | TRMT9B | tRNA methyltransferase 9B (putative) |
| | WTAP | WT1 associated protein |
| | IGFBP5 | insulin like growth factor binding protein 5 |
| | NKX2-1 | NK2 homeobox 1 |
| | C1QL3 | complement C1q like 3 |
| | MGAT2 | mannosyl (alpha-1,6-)-glycoprotein beta-1,2-N-acetylglucosaminyltransferase |
| | EMCN | endomucin |
| | PLSCR4 | phospholipid scramblase 4 |
| | EDEM3 | ER degradation enhancing alpha-mannosidase like protein 3 |
| | SAMD4A | sterile alpha motif domain containing 4A |
| | PRR3 | proline rich 3 |
| | OMG | oligodendrocyte myelin glycoprotein |
| | SMAD5 | SMAD family member 5 |
| | PDE3B | phosphodiesterase 3B |
| | SH3PXD2A | SH3 and PX domains 2A |
| | NAA25 | N(alpha)-acetyltransferase 25, NatB auxiliary subunit |
| | CALB1 | calbindin 1 |
| | LATS1 | large tumor suppressor kinase 1 |
| | CMIP | c-Maf inducing protein |
| | PCF11 | PCF11, cleavage and polyadenylation factor subunit |
| | ALDH1A3 | aldehyde dehydrogenase 1 family member A3 |
| | TPH1 | tryptophan hydroxylase 1 |
| | REST | RE1 silencing transcription factor |
| | PRELID2 | PRELI domain containing 2 |
| | DEK | DEK proto-oncogene |
| | RLIM | ring finger protein, LIM domain interacting |
| | IGF2BP2 | insulin like growth factor 2 mRNA binding protein 2 |
| | CNGB3 | cyclic nucleotide gated channel beta 3 |
| | SSH1 | slingshot protein phosphatase 1 |
| | SLCO5A1 | solute carrier organic anion transporter family member 5A1 |
| | NFASC | neurofascin |
| | FAM84A | family with sequence similarity 84 member A |

| | | |
|---|---|---|
| | ANKFY1 | ankyrin repeat and FYVE domain containing 1 |
| | ERO1B | endoplasmic reticulum oxidoreductase 1 beta |
| | NRXN3 | neurexin 3 |
| | NFYC | nuclear transcription factor Y subunit gamma |
| | DNMBP | dynamin binding protein |
| | TRIB2 | tribbles pseudokinase 2 |
| | U2SURP | U2 snRNP associated SURP domain containing |
| | THOC1 | THO complex 1 |
| | PAX9 | paired box 9 |
| | MIB1 | mindbomb E3 ubiquitin protein ligase 1 |
| | TMEM248 | transmembrane protein 248 |
| | PLEKHA2 | pleckstrin homology domain containing A2 |
| | TDRP | testis development related protein |
| | TP53INP1 | tumor protein p53 inducible nuclear protein 1 |
| | KLHL4 | kelch like family member 4 |
| | SPAG1 | sperm associated antigen 1 |
| | SLC2A12 | solute carrier family 2 member 12 |
| | KLF15 | Kruppel like factor 15 |
| | ZNF468 | zinc finger protein 468 |
| | PRAMEF19 | PRAME family member 19 |
| | IQCB1 | IQ motif containing B1 |
| | ARFGEF3 | ARFGEF family member 3 |
| | PDE11A | phosphodiesterase 11A |
| | GLIPR1 | GLI pathogenesis related 1 |
| | IRS1 | insulin receptor substrate 1 |
| | ZNF260 | zinc finger protein 260 |
| | SHC1 | SHC adaptor protein 1 |
| | LZIC | leucine zipper and CTNNBIP1 domain containing |
| | KDM3B | lysine demethylase 3B |
| | CTR9 | CTR9 homolog, Paf1/RNA polymerase II complex component |
| | CD2AP | CD2 associated protein |
| | AFF2 | AF4/FMR2 family member 2 |
| | PCDH11Y | protocadherin 11 Y-linked |
| | BRK1 | BRICK1, SCAR/WAVE actin nucleating complex subunit |
| | ASTN2 | astrotactin 2 |
| | PTPRE | protein tyrosine phosphatase, receptor type E |
| | ZDBF2 | zinc finger DBF-type containing 2 |
| | RALYL | RALY RNA binding protein like |
| | RANBP1 | RAN binding protein 1 |
| | CAB39 | calcium binding protein 39 |
| | DLST | dihydrolipoamide S-succinyltransferase |
| | BAG4 | BCL2 associated athanogene 4 |

| | | |
|---|---|---|
| | LIMS1 | LIM zinc finger domain containing 1 |
| | ZFYVE28 | zinc finger FYVE-type containing 28 |
| | RNF168 | ring finger protein 168 |
| | SLC39A10 | solute carrier family 39 member 10 |
| | ZNF366 | zinc finger protein 366 |
| | SSR3 | signal sequence receptor subunit 3 |
| | PTPN2 | protein tyrosine phosphatase, non-receptor type 2 |
| | EDIL3 | EGF like repeats and discoidin domains 3 |
| | BRD3 | bromodomain containing 3 |
| | LUZP2 | leucine zipper protein 2 |
| | CNOT6L | CCR4-NOT transcription complex subunit 6 like |
| | SCD | stearoyl-CoA desaturase |
| | EN2 | engrailed homeobox 2 |
| | MRPL42 | mitochondrial ribosomal protein L42 |
| | HLF | HLF, PAR bZIP transcription factor |
| | RC3H1 | ring finger and CCCH-type domains 1 |
| | NT5DC1 | 5'-nucleotidase domain containing 1 |
| | ERCC6L2 | ERCC excision repair 6 like 2 |
| | SELENOP | selenoprotein P |
| | SMAP1 | small ArfGAP 1 |
| | PRDM16 | PR/SET domain 16 |
| | ERC1 | ELKS/RAB6-interacting/CAST family member 1 |
| | CHD6 | chromodomain helicase DNA binding protein 6 |
| | FOXJ2 | forkhead box J2 |
| | ZIC3 | Zic family member 3 |
| | MAPK1 | mitogen-activated protein kinase 1 |
| | RIMKLB | ribosomal modification protein rimK like family member B |
| | CWF19L2 | CWF19 like cell cycle control factor 2 |
| | EDNRA | endothelin receptor type A |
| | WDR45B | WD repeat domain 45B |
| | GOSR1 | golgi SNAP receptor complex member 1 |
| | STMP1 | short transmembrane mitochondrial protein 1 |
| | ZSCAN22 | zinc finger and SCAN domain containing 22 |
| | UHRF2 | ubiquitin like with PHD and ring finger domains 2 |
| | REEP3 | receptor accessory protein 3 |
| | PPHLN1 | periphilin 1 |
| | ZNF787 | zinc finger protein 787 |
| | KLHL31 | kelch like family member 31 |
| | PCYT1B | phosphate cytidylyltransferase 1, choline, beta |
| | CREBRF | CREB3 regulatory factor |
| | DOCK3 | dedicator of cytokinesis 3 |
| | STXBP5 | syntaxin binding protein 5 |

| | | |
|---|---|---|
| | NKTR | natural killer cell triggering receptor |
| | TACC1 | transforming acidic coiled-coil containing protein 1 |
| | PRMT2 | protein arginine methyltransferase 2 |
| | RP2 | RP2, ARL3 GTPase activating protein |
| | DUSP18 | dual specificity phosphatase 18 |
| | STK35 | serine/threonine kinase 35 |
| | NR2C1 | nuclear receptor subfamily 2 group C member 1 |
| | TMEM132B | transmembrane protein 132B |
| | PRAMEF18 | PRAME family member 18 |
| | ENOSF1 | enolase superfamily member 1 |
| | CEP170 | centrosomal protein 170 |
| | ZNF264 | zinc finger protein 264 |
| | TNRC6B | trinucleotide repeat containing 6B |
| | CHAMP1 | chromosome alignment maintaining phosphoprotein 1 |
| | ITGAX | integrin subunit alpha X |
| | TGFBRAP1 | transforming growth factor beta receptor associated protein 1 |
| | YIPF6 | Yip1 domain family member 6 |
| | FAM199X | family with sequence similarity 199, X-linked |
| | ITPRIPL2 | ITPRIP like 2 |
| | PADI2 | peptidyl arginine deiminase 2 |
| | ZMYND11 | zinc finger MYND-type containing 11 |
| | WHAMM | WAS protein homolog associated with actin, golgi membranes and microtubules |
| | RAB6D | RAB6D, member RAS oncogene family |
| | IRGQ | immunity related GTPase Q |
| | DIAPH1 | diaphanous related formin 1 |
| | CBL | Cbl proto-oncogene |
| | MTSS1 | MTSS1, I-BAR domain containing |
| | PPP3CC | protein phosphatase 3 catalytic subunit gamma |
| | TNFRSF11A | TNF receptor superfamily member 11a |
| | SLC25A53 | solute carrier family 25 member 53 |
| | BCL10 | BCL10, immune signaling adaptor |
| | ADRB1 | adrenoceptor beta 1 |
| | AQP4 | aquaporin 4 |
| | ZNF850 | zinc finger protein 850 |
| | KLHL15 | kelch like family member 15 |
| | PALM2 | paralemmin 2 |
| | ZEB2 | zinc finger E-box binding homeobox 2 |
| | ELF1 | E74 like ETS transcription factor 1 |
| | STXBP6 | syntaxin binding protein 6 |
| | IFT88 | intraflagellar transport 88 |
| | EIF1AX | eukaryotic translation initiation factor 1A X-linked |

| | | |
|---|---|---|
| | FCGR1B | Fc fragment of IgG receptor Ib |
| | TXNIP | thioredoxin interacting protein |
| | SSR1 | signal sequence receptor subunit 1 |
| | ZNF124 | zinc finger protein 124 |
| | SPINK13 | serine peptidase inhibitor, Kazal type 13 (putative) |
| | BRINP2 | BMP/retinoic acid inducible neural specific 2 |
| | SOWAHB | sosondowah ankyrin repeat domain family member B |
| | SEC22A | SEC22 homolog A, vesicle trafficking protein |
| | FPR2 | formyl peptide receptor 2 |
| | GTDC1 | glycosyltransferase like domain containing 1 |
| | CDCA8 | cell division cycle associated 8 |
| | FAM3C | family with sequence similarity 3 member C |
| | MBTD1 | mbt domain containing 1 |
| | LRCH1 | leucine rich repeats and calponin homology domain containing 1 |
| | GTF2IRD2B | GTF2I repeat domain containing 2B |
| | TIMMDC1 | translocase of inner mitochondrial membrane domain containing 1 |
| | UBD | ubiquitin D |
| | ABLIM2 | actin binding LIM protein family member 2 |
| | ZBTB41 | zinc finger and BTB domain containing 41 |
| | MOBP | myelin-associated oligodendrocyte basic protein |
| | TFDP3 | transcription factor Dp family member 3 |
| | IFT81 | intraflagellar transport 81 |
| | SLC7A8 | solute carrier family 7 member 8 |
| | FAM122A | family with sequence similarity 122A |
| | TM9SF1 | transmembrane 9 superfamily member 1 |
| | OSBPL6 | oxysterol binding protein like 6 |
| | MLLT6 | MLLT6, PHD finger containing |
| | DCAF16 | DDB1 and CUL4 associated factor 16 |
| | VAPA | VAMP associated protein A |
| | MAPK9 | mitogen-activated protein kinase 9 |
| | MC2R | melanocortin 2 receptor |
| | PIK3CA | phosphatidylinositol-4,5-bisphosphate 3-kinase catalytic subunit alpha |
| | BRD4 | bromodomain containing 4 |
| | LEKR1 | leucine, glutamate and lysine rich 1 |
| | SZRD1 | SUZ RNA binding domain containing 1 |
| | CCNDBP1 | cyclin D1 binding protein 1 |
| | ATP1B1 | ATPase Na+/K+ transporting subunit beta 1 |
| | ANKRD17 | ankyrin repeat domain 17 |
| | PRKAA1 | protein kinase AMP-activated catalytic subunit alpha 1 |
| | HAND1 | heart and neural crest derivatives expressed 1 |

| | | |
|---|---|---|
| | CCNG2 | cyclin G2 |
| | FKBP5 | FKBP prolyl isomerase 5 |
| | STX11 | syntaxin 11 |
| | PCID2 | PCI domain containing 2 |
| | SOX8 | SRY-box 8 |
| | SLC16A7 | solute carrier family 16 member 7 |
| | CFLAR | CASP8 and FADD like apoptosis regulator |
| | CTNNA3 | catenin alpha 3 |
| | PSD3 | pleckstrin and Sec7 domain containing 3 |
| | CTBS | chitobiase |
| | MARCHF8 | membrane associated ring-CH-type finger 8 |
| | AES | amino-terminal enhancer of split |
| | STT3B | STT3B, catalytic subunit of the oligosaccharyltransferase complex |
| | SRCIN1 | SRC kinase signaling inhibitor 1 |
| | PTPN14 | protein tyrosine phosphatase, non-receptor type 14 |
| | USP4 | ubiquitin specific peptidase 4 |
| | CYBRD1 | cytochrome b reductase 1 |
| | B3GLCT | beta 3-glucosyltransferase |
| | CSNK2A1 | casein kinase 2 alpha 1 |
| | IGSF3 | immunoglobulin superfamily member 3 |
| | IGFBP4 | insulin like growth factor binding protein 4 |
| | CNTNAP2 | contactin associated protein like 2 |
| | PFKFB2 | 6-phosphofructo-2-kinase/fructose-2,6-biphosphatase 2 |
| | VXN | vexin |
| | TRIM45 | tripartite motif containing 45 |
| | SIRT1 | sirtuin 1 |
| | MED4 | mediator complex subunit 4 |
| | CEP44 | centrosomal protein 44 |
| | KLHL28 | kelch like family member 28 |
| | RHOQ | ras homolog family member Q |
| | EHF | ETS homologous factor |
| | SKA1 | spindle and kinetochore associated complex subunit 1 |
| | FAM84B | family with sequence similarity 84 member B |
| | WDR26 | WD repeat domain 26 |
| | CAST | calpastatin |
| | STARD4 | StAR related lipid transfer domain containing 4 |
| | GCA | grancalcin |
| | EXPH5 | exophilin 5 |
| | EGR3 | early growth response 3 |
| | MMS22L | MMS22 like, DNA repair protein |
| | PIAS4 | protein inhibitor of activated STAT 4 |

| | | |
|---|---|---|
| | PTEN | phosphatase and tensin homolog |
| | C10orf111 | chromosome 10 open reading frame 111 |
| | GNB1 | G protein subunit beta 1 |
| | PAXIP1 | PAX interacting protein 1 |
| | RBM14 | RNA binding motif protein 14 |
| | ETNK1 | ethanolamine kinase 1 |
| | GNAI1 | G protein subunit alpha i1 |
| | FBLN2 | fibulin 2 |
| | RCAN1 | regulator of calcineurin 1 |
| | ACVR1 | activin A receptor type 1 |
| | OR7D2 | olfactory receptor family 7 subfamily D member 2 |
| | GABRA1 | gamma-aminobutyric acid type A receptor alpha1 subunit |
| | ARL17A | ADP ribosylation factor like GTPase 17A |
| | ADGRG7 | adhesion G protein-coupled receptor G7 |
| | ST18 | ST18, C2H2C-type zinc finger |
| | STON2 | stonin 2 |
| | FAM135A | family with sequence similarity 135 member A |
| | C2orf69 | chromosome 2 open reading frame 69 |
| | NPPC | natriuretic peptide C |
| | CNTLN | centlein |
| | ARL6IP6 | ADP ribosylation factor like GTPase 6 interacting protein 6 |
| | LENG8 | leukocyte receptor cluster member 8 |
| | PDE1A | phosphodiesterase 1A |
| | SPATA22 | spermatogenesis associated 22 |
| | CDH2 | cadherin 2 |
| | ANKRD10 | ankyrin repeat domain 10 |
| | MLF1 | myeloid leukemia factor 1 |
| | PTPN21 | protein tyrosine phosphatase, non-receptor type 21 |
| | PUM2 | pumilio RNA binding family member 2 |
| | RBM46 | RNA binding motif protein 46 |
| | AIG1 | androgen induced 1 |
| | TRIL | TLR4 interactor with leucine rich repeats |
| | SLC2A13 | solute carrier family 2 member 13 |
| | PHF2 | PHD finger protein 2 |
| | RER1 | retention in endoplasmic reticulum sorting receptor 1 |
| | CASP10 | caspase 10 |
| | KIAA1210 | KIAA1210 |
| | FBXO44 | F-box protein 44 |
| | SLC44A1 | solute carrier family 44 member 1 |
| | KCNJ4 | potassium voltage-gated channel subfamily J member 4 |
| | CCDC141 | coiled-coil domain containing 141 |
| | PARP8 | poly(ADP-ribose) polymerase family member 8 |

| | | |
|---|---|---|
| | XRCC2 | X-ray repair cross complementing 2 |
| | NDEL1 | nudE neurodevelopment protein 1 like 1 |
| | TAB2 | TGF-beta activated kinase 1 (MAP3K7) binding protein 2 |
| | RALA | RAS like proto-oncogene A |
| | KHDRBS3 | KH RNA binding domain containing, signal transduction associated 3 |
| | WDR48 | WD repeat domain 48 |
| | TMEM260 | transmembrane protein 260 |
| | ABL2 | ABL proto-oncogene 2, non-receptor tyrosine kinase |
| | RBMS1 | RNA binding motif single stranded interacting protein 1 |
| | PPP1CA | protein phosphatase 1 catalytic subunit alpha |
| | ABCC9 | ATP binding cassette subfamily C member 9 |
| | CDCA3 | cell division cycle associated 3 |
| | AKAP10 | A-kinase anchoring protein 10 |
| | STK11 | serine/threonine kinase 11 |
| | RUNX2 | runt related transcription factor 2 |
| | GABPA | GA binding protein transcription factor subunit alpha |
| | ID2 | inhibitor of DNA binding 2 |
| | ODF2L | outer dense fiber of sperm tails 2 like |
| | EML2 | EMAP like 2 |
| | IKZF1 | IKAROS family zinc finger 1 |
| | ARFGEF1 | ADP ribosylation factor guanine nucleotide exchange factor 1 |
| | TNRC18 | trinucleotide repeat containing 18 |
| | ADAR | adenosine deaminase, RNA specific |
| | WDFY1 | WD repeat and FYVE domain containing 1 |
| | FBXO36 | F-box protein 36 |
| | CPED1 | cadherin like and PC-esterase domain containing 1 |
| | FRMD4A | FERM domain containing 4A |
| | FRMD6 | FERM domain containing 6 |
| | LSM8 | LSM8 homolog, U6 small nuclear RNA associated |
| | DCP1A | decapping mRNA 1A |
| | PPIC | peptidylprolyl isomerase C |
| | GLYATL3 | glycine-N-acyltransferase like 3 |
| | SLC35F5 | solute carrier family 35 member F5 |
| | SLC35D1 | solute carrier family 35 member D1 |
| | MED7 | mediator complex subunit 7 |
| | LARP4 | La ribonucleoprotein domain family member 4 |
| | CREBBP | CREB binding protein |
| | PRKD3 | protein kinase D3 |
| | C1orf109 | chromosome 1 open reading frame 109 |
| | C5orf22 | chromosome 5 open reading frame 22 |
| | SLC9A4 | solute carrier family 9 member A4 |

| | | |
|---|---|---|
| | MRPL50 | mitochondrial ribosomal protein L50 |
| | CETN2 | centrin 2 |
| | CLVS2 | clavesin 2 |
| | TGFB2 | transforming growth factor beta 2 |
| | ACSL4 | acyl-CoA synthetase long chain family member 4 |
| | BTAF1 | B-TFIID TATA-box binding protein associated factor 1 |
| | WWC3 | WWC family member 3 |
| | ZNF460 | zinc finger protein 460 |
| | GRID2 | glutamate ionotropic receptor delta type subunit 2 |
| | TLR5 | toll like receptor 5 |
| | ZNF106 | zinc finger protein 106 |
| | FAM102B | family with sequence similarity 102 member B |
| | ZNF516 | zinc finger protein 516 |
| | MAP2 | microtubule associated protein 2 |
| | SLC4A7 | solute carrier family 4 member 7 |
| | CYP4V2 | cytochrome P450 family 4 subfamily V member 2 |
| | CCDC146 | coiled-coil domain containing 146 |
| | RGS7BP | regulator of G protein signaling 7 binding protein |
| | PSME3 | proteasome activator subunit 3 |
| | ICE2 | interactor of little elongation complex ELL subunit 2 |
| | THAP10 | THAP domain containing 10 |
| | CSN1S1 | casein alpha s1 |
| | SLC5A3 | solute carrier family 5 member 3 |
| | KATNAL1 | katanin catalytic subunit A1 like 1 |
| | FASTKD2 | FAST kinase domains 2 |
| | NCMAP | non-compact myelin associated protein |
| | TMEM178B | transmembrane protein 178B |
| | LINGO2 | leucine rich repeat and Ig domain containing 2 |
| | SLC23A2 | solute carrier family 23 member 2 |
| | ZFP82 | ZFP82 zinc finger protein |
| | FOXI1 | forkhead box I1 |
| | DUSP5 | dual specificity phosphatase 5 |
| | SOSTDC1 | sclerostin domain containing 1 |
| | SLC30A9 | solute carrier family 30 member 9 |
| | TCEAL7 | transcription elongation factor A like 7 |
| | ZNF552 | zinc finger protein 552 |
| | DYTN | dystrotelin |
| | SLC16A9 | solute carrier family 16 member 9 |
| | RBM11 | RNA binding motif protein 11 |
| | LRRC2 | leucine rich repeat containing 2 |
| | RASGRF2 | Ras protein specific guanine nucleotide releasing factor 2 |
| | MAP3K21 | mitogen-activated protein kinase kinase kinase 21 |

| | | |
|---|---|---|
| | ASAH1 | N-acylsphingosine amidohydrolase 1 |
| | ZNF28 | zinc finger protein 28 |
| | TRIP11 | thyroid hormone receptor interactor 11 |
| | AHNAK | AHNAK nucleoprotein |
| | PPIL1 | peptidylprolyl isomerase like 1 |
| | MYLK4 | myosin light chain kinase family member 4 |
| | APOLD1 | apolipoprotein L domain containing 1 |
| | NDUFA4 | NDUFA4, mitochondrial complex associated |
| | GATAD2B | GATA zinc finger domain containing 2B |
| | SERAC1 | serine active site containing 1 |
| | TMEM106B | transmembrane protein 106B |
| | RAB8B | RAB8B, member RAS oncogene family |
| | CCPG1 | cell cycle progression 1 |
| | ADNP | activity dependent neuroprotector homeobox |
| | CALHM5 | calcium homeostasis modulator family member 5 |
| | FEM1B | fem-1 homolog B |
| | KCTD6 | potassium channel tetramerization domain containing 6 |
| | CMTM4 | CKLF like MARVEL transmembrane domain containing 4 |
| | CCDC80 | coiled-coil domain containing 80 |
| | SERPINB4 | serpin family B member 4 |
| | NSD1 | nuclear receptor binding SET domain protein 1 |
| | SFMBT1 | Scm like with four mbt domains 1 |
| | NUCKS1 | nuclear casein kinase and cyclin dependent kinase substrate 1 |
| | C7orf31 | chromosome 7 open reading frame 31 |
| | SERPINB3 | serpin family B member 3 |
| | RAB6A | RAB6A, member RAS oncogene family |
| | MAGEB16 | MAGE family member B16 |
| | SLC6A11 | solute carrier family 6 member 11 |
| | MYF5 | myogenic factor 5 |
| | PFDN4 | prefoldin subunit 4 |
| | CNOT6 | CCR4-NOT transcription complex subunit 6 |
| | SKIDA1 | SKI/DACH domain containing 1 |
| | ZNF761 | zinc finger protein 761 |
| | ELAVL3 | ELAV like RNA binding protein 3 |
| | DIO2 | iodothyronine deiodinase 2 |
| | SOCS6 | suppressor of cytokine signaling 6 |
| | CES5A | carboxylesterase 5A |
| | UBE2E2 | ubiquitin conjugating enzyme E2 E2 |
| | MED10 | mediator complex subunit 10 |
| | C5 | complement C5 |
| | ZNF792 | zinc finger protein 792 |
| | SGCZ | sarcoglycan zeta |

| | | |
|---|---|---|
| | ZNF292 | zinc finger protein 292 |
| | SLC25A26 | solute carrier family 25 member 26 |
| | CLDN18 | claudin 18 |
| | NCEH1 | neutral cholesterol ester hydrolase 1 |
| | LRRC8D | leucine rich repeat containing 8 VRAC subunit D |
| | ESM1 | endothelial cell specific molecule 1 |
| | LDLRAD3 | low density lipoprotein receptor class A domain containing 3 |
| | GFPT1 | glutamine--fructose-6-phosphate transaminase 1 |
| | KCTD14 | potassium channel tetramerization domain containing 14 |
| | MAN1A2 | mannosidase alpha class 1A member 2 |
| | TBC1D19 | TBC1 domain family member 19 |
| | EREG | epiregulin |
| | ZHX2 | zinc fingers and homeoboxes 2 |
| | APC | APC, WNT signaling pathway regulator |
| | GPR65 | G protein-coupled receptor 65 |
| | KLF6 | Kruppel like factor 6 |
| | ME3 | malic enzyme 3 |
| | BBX | BBX, HMG-box containing |
| | ANK3 | ankyrin 3 |
| | PLRG1 | pleiotropic regulator 1 |
| | RAD54B | RAD54 homolog B |
| | EMP2 | epithelial membrane protein 2 |
| | TIMM17A | translocase of inner mitochondrial membrane 17A |
| | KLHL42 | kelch like family member 42 |
| | KIAA0232 | KIAA0232 |
| | PPP2R2A | protein phosphatase 2 regulatory subunit Balpha |
| | EIF4ENIF1 | eukaryotic translation initiation factor 4E nuclear import factor 1 |
| | CLIC4 | chloride intracellular channel 4 |
| | NHLRC3 | NHL repeat containing 3 |
| | TXNRD2 | thioredoxin reductase 2 |
| | SAMD8 | sterile alpha motif domain containing 8 |
| | SLC17A5 | solute carrier family 17 member 5 |
| | TMX1 | thioredoxin related transmembrane protein 1 |
| | MRPL11 | mitochondrial ribosomal protein L11 |
| | CRLS1 | cardiolipin synthase 1 |
| | WDR72 | WD repeat domain 72 |
| | SERPINB12 | serpin family B member 12 |
| | LRBA | LPS responsive beige-like anchor protein |
| | ZNF772 | zinc finger protein 772 |
| | ITCH | itchy E3 ubiquitin protein ligase |
| | GPATCH2 | G-patch domain containing 2 |

| | | |
|---|---|---|
| | GRIK4 | glutamate ionotropic receptor kainate type subunit 4 |
| | SET | SET nuclear proto-oncogene |
| | MPP6 | membrane palmitoylated protein 6 |
| | POU2AF1 | POU class 2 associating factor 1 |
| | GABBR2 | gamma-aminobutyric acid type B receptor subunit 2 |
| | APPBP2 | amyloid beta precursor protein binding protein 2 |
| | B4GALT6 | beta-1,4-galactosyltransferase 6 |
| | NAA16 | N(alpha)-acetyltransferase 16, NatA auxiliary subunit |
| | NFYB | nuclear transcription factor Y subunit beta |
| | RASL11B | RAS like family 11 member B |
| | FAM169A | family with sequence similarity 169 member A |
| | SAV1 | salvador family WW domain containing protein 1 |
| | FOXL2 | forkhead box L2 |
| | BNC1 | basonuclin 1 |
| | GTF2IRD2 | GTF2I repeat domain containing 2 |
| | SPRTN | SprT-like N-terminal domain |
| | ANXA13 | annexin A13 |
| | ZDHHC6 | zinc finger DHHC-type containing 6 |
| | LIMS4 | LIM zinc finger domain containing 4 |
| | RWDD4 | RWD domain containing 4 |
| | ECPAS | Ecm29 proteasome adaptor and scaffold |
| | MFSD14A | major facilitator superfamily domain containing 14A |
| | HOXA11 | homeobox A11 |
| | PRH1-TAS2R14 | PRH1-TAS2R14 readthrough |
| | SRP54 | signal recognition particle 54 |
| | KRBOX1 | KRAB box domain containing 1 |
| | GID8 | GID complex subunit 8 homolog |
| | NPAS3 | neuronal PAS domain protein 3 |
| | SRSF2 | serine and arginine rich splicing factor 2 |
| | TBC1D7 | TBC1 domain family member 7 |
| | PXT1 | peroxisomal testis enriched protein 1 |
| | ICA1L | islet cell autoantigen 1 like |
| | SEPHS1 | selenophosphate synthetase 1 |
| | MAGEB18 | MAGE family member B18 |
| | GCC1 | GRIP and coiled-coil domain containing 1 |
| | C2CD6 | C2 calcium dependent domain containing 6 |
| | AMOTL2 | angiomotin like 2 |
| | FLRT2 | fibronectin leucine rich transmembrane protein 2 |
| | CHRNB4 | cholinergic receptor nicotinic beta 4 subunit |
| | RAB6C | RAB6C, member RAS oncogene family |
| | DCAF4L1 | DDB1 and CUL4 associated factor 4 like 1 |
| | G2E3 | G2/M-phase specific E3 ubiquitin protein ligase |

| | | |
|---|---|---|
| | NLK | nemo like kinase |
| | SF3B1 | splicing factor 3b subunit 1 |
| | CSRP2 | cysteine and glycine rich protein 2 |
| | ZNF558 | zinc finger protein 558 |
| | LZTS1 | leucine zipper tumor suppressor 1 |
| | NAMPT | nicotinamide phosphoribosyltransferase |
| | PLAC8 | placenta specific 8 |
| | METTL21A | methyltransferase like 21A |
| | MCRS1 | microspherule protein 1 |
| | C14orf39 | chromosome 14 open reading frame 39 |
| | SKIL | SKI like proto-oncogene |
| | C14orf28 | chromosome 14 open reading frame 28 |
| | XRN1 | 5'-3' exoribonuclease 1 |
| | TUBGCP3 | tubulin gamma complex associated protein 3 |
| | ITGB8 | integrin subunit beta 8 |
| | GEN1 | GEN1, Holliday junction 5' flap endonuclease |
| | RPS6KA5 | ribosomal protein S6 kinase A5 |
| | POLA2 | DNA polymerase alpha 2, accessory subunit |
| | DCAKD | dephospho-CoA kinase domain containing |
| | LYRM9 | LYR motif containing 9 |
| | MAST3 | microtubule associated serine/threonine kinase 3 |
| | LSM6 | LSM6 homolog, U6 small nuclear RNA and mRNA degradation associated |
| | BRCA2 | BRCA2, DNA repair associated |
| | ZNF322 | zinc finger protein 322 |
| | TMED7 | transmembrane p24 trafficking protein 7 |
| | MAPRE1 | microtubule associated protein RP/EB family member 1 |
| | GCLM | glutamate-cysteine ligase modifier subunit |
| | RAD51B | RAD51 paralog B |
| | TC2N | tandem C2 domains, nuclear |
| | ARHGEF12 | Rho guanine nucleotide exchange factor 12 |
| | DUSP7 | dual specificity phosphatase 7 |
| | CEACAM5 | carcinoembryonic antigen related cell adhesion molecule 5 |
| | CEBPG | CCAAT enhancer binding protein gamma |
| | PTPRG | protein tyrosine phosphatase, receptor type G |
| | CCNJ | cyclin J |
| | KARS | lysyl-tRNA synthetase |
| | ASF1A | anti-silencing function 1A histone chaperone |
| | KIT | KIT proto-oncogene receptor tyrosine kinase |
| | XIAP | X-linked inhibitor of apoptosis |
| | ACAP2 | ArfGAP with coiled-coil, ankyrin repeat and PH domains 2 |
| | USP3 | ubiquitin specific peptidase 3 |

| | | |
|---|---|---|
| | RNPS1 | RNA binding protein with serine rich domain 1 |
| | WDR1 | WD repeat domain 1 |
| | DLX6 | distal-less homeobox 6 |
| | TEX12 | testis expressed 12 |
| | ATP1B2 | ATPase Na+/K+ transporting subunit beta 2 |
| | TCP10L2 | t-complex 10 like 2 |
| | ZDHHC15 | zinc finger DHHC-type containing 15 |
| | MED12L | mediator complex subunit 12 like |
| | TMPO | thymopoietin |
| | MARCKS | myristoylated alanine rich protein kinase C substrate |
| | ST8SIA4 | ST8 alpha-N-acetyl-neuraminide alpha-2,8-sialyltransferase 4 |
| | DNAJB14 | DnaJ heat shock protein family (Hsp40) member B14 |
| | CA13 | carbonic anhydrase 13 |
| | SH3BGR | SH3 domain binding glutamate rich protein |
| | WRB-SH3BGR | WRB-SH3BGR readthrough |
| | COL11A1 | collagen type XI alpha 1 chain |
| | SOCS7 | suppressor of cytokine signaling 7 |
| | ERI2 | ERI1 exoribonuclease family member 2 |
| | SHOC2 | SHOC2, leucine rich repeat scaffold protein |
| | C6orf89 | chromosome 6 open reading frame 89 |
| | RAI2 | retinoic acid induced 2 |
| | DPYD | dihydropyrimidine dehydrogenase |
| | AP3S1 | adaptor related protein complex 3 subunit sigma 1 |
| | SORL1 | sortilin related receptor 1 |
| | USP40 | ubiquitin specific peptidase 40 |
| | ITGA2 | integrin subunit alpha 2 |
| | C8orf37 | chromosome 8 open reading frame 37 |
| | SRGAP3 | SLIT-ROBO Rho GTPase activating protein 3 |
| | RICTOR | RPTOR independent companion of MTOR complex 2 |
| | KDM5B | lysine demethylase 5B |
| | OLFML1 | olfactomedin like 1 |
| | WWP2 | WW domain containing E3 ubiquitin protein ligase 2 |
| | UPF2 | UPF2, regulator of nonsense mediated mRNA decay |
| | PELI2 | pellino E3 ubiquitin protein ligase family member 2 |
| | RRS1 | ribosome biogenesis regulator homolog |
| | RASSF3 | Ras association domain family member 3 |
| | ZNF765 | zinc finger protein 765 |
| | ZC3H6 | zinc finger CCCH-type containing 6 |
| | PRELID3B | PRELI domain containing 3B |
| | BEST3 | bestrophin 3 |
| | MOCS1 | molybdenum cofactor synthesis 1 |
| | NOS1 | nitric oxide synthase 1 |

| | | |
|---|---|---|
| | GSTCD | glutathione S-transferase C-terminal domain containing |
| | AKIRIN1 | akirin 1 |
| | CWC15 | CWC15 spliceosome associated protein homolog |
| | RNF182 | ring finger protein 182 |
| | TNFSF10 | TNF superfamily member 10 |
| | GPR83 | G protein-coupled receptor 83 |
| | BICRAL | BRD4 interacting chromatin remodeling complex associated protein like |
| | EMSY | EMSY, BRCA2 interacting transcriptional repressor |
| | CPNE5 | copine 5 |
| | ZIC1 | Zic family member 1 |
| | IKZF3 | IKAROS family zinc finger 3 |
| | RELL1 | RELT like 1 |
| | EP300 | E1A binding protein p300 |
| | PCMTD2 | protein-L-isoaspartate (D-aspartate) O-methyltransferase domain containing 2 |
| | CUX2 | cut like homeobox 2 |
| | ST6GAL2 | ST6 beta-galactoside alpha-2,6-sialyltransferase 2 |
| | STIL | STIL, centriolar assembly protein |
| | CALN1 | calneuron 1 |
| | MBNL1 | muscleblind like splicing regulator 1 |
| | YAP1 | Yes associated protein 1 |
| | CHST11 | carbohydrate sulfotransferase 11 |
| | ANKRD52 | ankyrin repeat domain 52 |
| | FBXO33 | F-box protein 33 |
| | TFPI | tissue factor pathway inhibitor |
| | EIF5 | eukaryotic translation initiation factor 5 |
| | HOXA13 | homeobox A13 |
| | GINS1 | GINS complex subunit 1 |
| | BTG2 | BTG anti-proliferation factor 2 |
| | NCOA7 | nuclear receptor coactivator 7 |
| | RAPGEF6 | Rap guanine nucleotide exchange factor 6 |
| | CYB5A | cytochrome b5 type A |
| | LRRC34 | leucine rich repeat containing 34 |
| | GSDMA | gasdermin A |
| | ZNF700 | zinc finger protein 700 |
| | KDM7A | lysine demethylase 7A |
| | TGFA | transforming growth factor alpha |
| | H2AFY | H2A histone family member Y |
| | LRP6 | LDL receptor related protein 6 |
| | UBFD1 | ubiquitin family domain containing 1 |
| | KIN | Kin17 DNA and RNA binding protein |
| | SIAH1 | siah E3 ubiquitin protein ligase 1 |

| | | |
|---|---|---|
| | TENT2 | terminal nucleotidyltransferase 2 |
| | YPEL5 | yippee like 5 |
| | TTC16 | tetratricopeptide repeat domain 16 |
| | WDR91 | WD repeat domain 91 |
| | AZIN1 | antizyme inhibitor 1 |
| | GCC2 | GRIP and coiled-coil domain containing 2 |
| | ATP2B4 | ATPase plasma membrane Ca2+ transporting 4 |
| | GPR158 | G protein-coupled receptor 158 |
| | DACT3 | dishevelled binding antagonist of beta catenin 3 |
| | ARHGEF35 | Rho guanine nucleotide exchange factor 35 |
| | ADGRG2 | adhesion G protein-coupled receptor G2 |
| | EEA1 | early endosome antigen 1 |
| | IL1RAP | interleukin 1 receptor accessory protein |
| | NDUFC2-KCTD14 | NDUFC2-KCTD14 readthrough |
| | SHROOM4 | shroom family member 4 |
| | ASB7 | ankyrin repeat and SOCS box containing 7 |
| | AHSA1 | activator of HSP90 ATPase activity 1 |
| | FAM241A | family with sequence similarity 241 member A |
| | TBC1D12 | TBC1 domain family member 12 |
| | KNSTRN | kinetochore localized astrin (SPAG5) binding protein |
| | FAT3 | FAT atypical cadherin 3 |
| | F5 | coagulation factor V |
| | TARBP1 | TAR (HIV-1) RNA binding protein 1 |
| | CELF4 | CUGBP Elav-like family member 4 |
| | MFHAS1 | malignant fibrous histiocytoma amplified sequence 1 |
| | CBX5 | chromobox 5 |
| | SPDYE6 | speedy/RINGO cell cycle regulator family member E6 |
| | MPZL3 | myelin protein zero like 3 |
| | FAM83H | family with sequence similarity 83 member H |
| | ARHGEF26 | Rho guanine nucleotide exchange factor 26 |
| | TP63 | tumor protein p63 |
| | CBX6 | chromobox 6 |
| | ZNF268 | zinc finger protein 268 |
| | MYOM3 | myomesin 3 |
| | GPR52 | G protein-coupled receptor 52 |
| | GOLPH3 | golgi phosphoprotein 3 |
| | EPC1 | enhancer of polycomb homolog 1 |
| | ITGA6 | integrin subunit alpha 6 |
| | IGSF21 | immunoglobin superfamily member 21 |
| | SEPTIN14 | septin 14 |
| | OPN3 | opsin 3 |

| | | |
|---|---|---|
| | DSTYK | dual serine/threonine and tyrosine protein kinase |
| | ANKRD29 | ankyrin repeat domain 29 |
| | BMP2K | BMP2 inducible kinase |
| | GK5 | glycerol kinase 5 |
| | ADAT2 | adenosine deaminase, tRNA specific 2 |
| | ARHGAP26 | Rho GTPase activating protein 26 |
| | SASH1 | SAM and SH3 domain containing 1 |
| | GATA3 | GATA binding protein 3 |
| | TMEM241 | transmembrane protein 241 |
| | HS6ST3 | heparan sulfate 6-O-sulfotransferase 3 |
| | CXADR | CXADR, Ig-like cell adhesion molecule |
| | AGPS | alkylglycerone phosphate synthase |
| | LNPK | lunapark, ER junction formation factor |
| | SPRY2 | sprouty RTK signaling antagonist 2 |
| | OSTM1 | osteoclastogenesis associated transmembrane protein 1 |
| | SLC7A1 | solute carrier family 7 member 1 |
| | GLRX | glutaredoxin |
| | KAT6A | lysine acetyltransferase 6A |
| | TMEM243 | transmembrane protein 243 |
| | SMIM13 | small integral membrane protein 13 |
| | FUT9 | fucosyltransferase 9 |
| | GATA6 | GATA binding protein 6 |
| | MGAM2 | maltase-glucoamylase 2 (putative) |
| | SLC30A5 | solute carrier family 30 member 5 |
| | SIX1 | SIX homeobox 1 |
| | CXXC4 | CXXC finger protein 4 |
| | MAP3K20 | mitogen-activated protein kinase kinase kinase 20 |
| | LOX | lysyl oxidase |
| | P4HA3 | prolyl 4-hydroxylase subunit alpha 3 |
| | TRPC5 | transient receptor potential cation channel subfamily C member 5 |
| | MAPK13 | mitogen-activated protein kinase 13 |
| | ANKRD50 | ankyrin repeat domain 50 |
| | THEMIS | thymocyte selection associated |
| | SLTM | SAFB like transcription modulator |
| | MAP3K8 | mitogen-activated protein kinase kinase kinase 8 |
| | SP110 | SP110 nuclear body protein |
| | USP46 | ubiquitin specific peptidase 46 |
| | LIN28B | lin-28 homolog B |
| | CUL3 | cullin 3 |
| | ADSS | adenylosuccinate synthase |
| | ZNF860 | zinc finger protein 860 |

| | | |
|---|---|---|
| | TMEM123 | transmembrane protein 123 |
| | ZNF544 | zinc finger protein 544 |
| | TMEM56 | transmembrane protein 56 |
| | DDX43 | DEAD-box helicase 43 |
| | ZNF805 | zinc finger protein 805 |
| | COPG2 | coatomer protein complex subunit gamma 2 |
| | HDX | highly divergent homeobox |
| | KIF27 | kinesin family member 27 |
| | SETD5 | SET domain containing 5 |
| | MORF4L1 | mortality factor 4 like 1 |
| | ENAM | enamelin |
| | TXNRD1 | thioredoxin reductase 1 |
| | CTTNBP2NL | CTTNBP2 N-terminal like |
| | ZNF662 | zinc finger protein 662 |
| | FOXL2NB | FOXL2 neighbor |
| | SLC38A4 | solute carrier family 38 member 4 |
| | INPP5B | inositol polyphosphate-5-phosphatase B |
| | GLT8D2 | glycosyltransferase 8 domain containing 2 |
| | RNF6 | ring finger protein 6 |
| | CCDC186 | coiled-coil domain containing 186 |
| | TRIM22 | tripartite motif containing 22 |
| | SRCAP | Snf2 related CREBBP activator protein |
| | EEF1AKMT2 | EEF1A lysine methyltransferase 2 |
| | ACTR2 | ARP2 actin related protein 2 homolog |
| | GSR | glutathione-disulfide reductase |
| | NMNAT2 | nicotinamide nucleotide adenylyltransferase 2 |
| | GALNT4 | polypeptide N-acetylgalactosaminyltransferase 4 |
| | POC1B-GALNT4 | POC1B-GALNT4 readthrough |
| | CLTRN | collectrin, amino acid transport regulator |
| | HS3ST5 | heparan sulfate-glucosamine 3-sulfotransferase 5 |
| | DOCK7 | dedicator of cytokinesis 7 |
| | SLC7A7 | solute carrier family 7 member 7 |
| | USP49 | ubiquitin specific peptidase 49 |
| | GCG | glucagon |
| | EFNB1 | ephrin B1 |
| | DKK2 | dickkopf WNT signaling pathway inhibitor 2 |
| | UBXN2B | UBX domain protein 2B |
| | TKTL2 | transketolase like 2 |
| | POLD3 | DNA polymerase delta 3, accessory subunit |
| | FRK | fyn related Src family tyrosine kinase |
| | CTTN | cortactin |

| | | |
|---|---|---|
| | ASB16 | ankyrin repeat and SOCS box containing 16 |
| | C3orf14 | chromosome 3 open reading frame 14 |
| | ALG10 | ALG10, alpha-1,2-glucosyltransferase |
| | PCDH11X | protocadherin 11 X-linked |
| | ATP8A1 | ATPase phospholipid transporting 8A1 |
| | LRRC49 | leucine rich repeat containing 49 |
| | MSH4 | mutS homolog 4 |
| | PLEKHA1 | pleckstrin homology domain containing A1 |
| | LMLN | leishmanolysin like peptidase |
| | SNF8 | SNF8, ESCRT-II complex subunit |
| | RAB3B | RAB3B, member RAS oncogene family |
| | IER5L | immediate early response 5 like |
| | MIER1 | MIER1 transcriptional regulator |
| | TP53I11 | tumor protein p53 inducible protein 11 |
| | CYSLTR1 | cysteinyl leukotriene receptor 1 |
| | LTBP1 | latent transforming growth factor beta binding protein 1 |
| | BMT2 | base methyltransferase of 25S rRNA 2 homolog |
| | IL21R | interleukin 21 receptor |
| | CHRD | chordin |
| | KLF11 | Kruppel like factor 11 |
| | UGT8 | UDP glycosyltransferase 8 |
| | PPFIA2 | PTPRF interacting protein alpha 2 |
| | KCTD8 | potassium channel tetramerization domain containing 8 |
| | FTO | FTO, alpha-ketoglutarate dependent dioxygenase |
| | RRAGD | Ras related GTP binding D |
| | DCX | doublecortin |
| | NAPEPLD | N-acyl phosphatidylethanolamine phospholipase D |
| | SNX20 | sorting nexin 20 |
| | PAPOLG | poly(A) polymerase gamma |
| | KLF8 | Kruppel like factor 8 |
| | ZNF550 | zinc finger protein 550 |
| | IL2 | interleukin 2 |
| | PDCD6IP | programmed cell death 6 interacting protein |
| | ABCC1 | ATP binding cassette subfamily C member 1 |
| | CCN3 | cellular communication network factor 3 |
| | SEPTIN11 | septin 11 |
| | SRGAP2B | SLIT-ROBO Rho GTPase activating protein 2B |
| | PPM1L | protein phosphatase, Mg2+/Mn2+ dependent 1L |
| | KIF3A | kinesin family member 3A |
| | CYP7B1 | cytochrome P450 family 7 subfamily B member 1 |
| | SCAF8 | SR-related CTD associated factor 8 |
| | MGARP | mitochondria localized glutamic acid rich protein |

| | | |
|---|---|---|
| | TTC21B | tetratricopeptide repeat domain 21B |
| | C5orf30 | chromosome 5 open reading frame 30 |
| | SIGLEC6 | sialic acid binding Ig like lectin 6 |
| | ANKRD40 | ankyrin repeat domain 40 |
| | MSL2 | MSL complex subunit 2 |
| | PTGS2 | prostaglandin-endoperoxide synthase 2 |
| | SPTY2D1 | SPT2 chromatin protein domain containing 1 |
| | CLK4 | CDC like kinase 4 |
| | COL24A1 | collagen type XXIV alpha 1 chain |
| | TIMM23B | translocase of inner mitochondrial membrane 23 homolog B |
| | TOR1B | torsin family 1 member B |
| | DNAJC27 | DnaJ heat shock protein family (Hsp40) member C27 |
| | GOLGA4 | golgin A4 |
| | MARVELD3 | MARVEL domain containing 3 |
| | CMTR2 | cap methyltransferase 2 |
| | ANKRD34C | ankyrin repeat domain 34C |
| | ADGRL2 | adhesion G protein-coupled receptor L2 |
| | VWA2 | von Willebrand factor A domain containing 2 |
| | TMEM150C | transmembrane protein 150C |
| | VPS4B | vacuolar protein sorting 4 homolog B |
| | ANKRD62 | ankyrin repeat domain 62 |
| | NEGR1 | neuronal growth regulator 1 |
| | CDK17 | cyclin dependent kinase 17 |
| | PLEKHH2 | pleckstrin homology, MyTH4 and FERM domain containing H2 |
| | DLC1 | DLC1 Rho GTPase activating protein |
| | ZNF600 | zinc finger protein 600 |
| | PRPS2 | phosphoribosyl pyrophosphate synthetase 2 |
| | CA5B | carbonic anhydrase 5B |
| | KLLN | killin, p53 regulated DNA replication inhibitor |
| | TRAPPC6B | trafficking protein particle complex 6B |
| | ZBTB14 | zinc finger and BTB domain containing 14 |
| | CTDSPL | CTD small phosphatase like |
| | HECW1 | HECT, C2 and WW domain containing E3 ubiquitin protein ligase 1 |
| | LHFPL6 | LHFPL tetraspan subfamily member 6 |
| | ATF7IP | activating transcription factor 7 interacting protein |
| | FAM217B | family with sequence similarity 217 member B |
| | CMTM6 | CKLF like MARVEL transmembrane domain containing 6 |
| | SAR1B | secretion associated Ras related GTPase 1B |
| | ELL | elongation factor for RNA polymerase II |
| | CADM1 | cell adhesion molecule 1 |
| | TMEM192 | transmembrane protein 192 |

| | | |
|---|---|---|
| | LRRC1 | leucine rich repeat containing 1 |
| | ITPR2 | inositol 1,4,5-trisphosphate receptor type 2 |
| | MNX1 | motor neuron and pancreas homeobox 1 |
| | TTC39C | tetratricopeptide repeat domain 39C |
| | NANP | N-acetylneuraminic acid phosphatase |
| | TRIO | trio Rho guanine nucleotide exchange factor |
| | EPPIN | epididymal peptidase inhibitor |
| | ITGAM | integrin subunit alpha M |
| | USO1 | USO1 vesicle transport factor |
| | DIP2A | disco interacting protein 2 homolog A |
| | SESN3 | sestrin 3 |
| | ISOC1 | isochorismatase domain containing 1 |
| | NEK2 | NIMA related kinase 2 |
| | TES | testin LIM domain protein |
| | CETN3 | centrin 3 |
| | MXD1 | MAX dimerization protein 1 |
| | SLC10A7 | solute carrier family 10 member 7 |
| | TPM3 | tropomyosin 3 |
| | NOC2L | NOC2 like nucleolar associated transcriptional repressor |
| | THRB | thyroid hormone receptor beta |
| | AK6 | adenylate kinase 6 |
| | MOSPD1 | motile sperm domain containing 1 |
| | FBXL20 | F-box and leucine rich repeat protein 20 |
| | HIC2 | HIC ZBTB transcriptional repressor 2 |
| | OSR1 | odd-skipped related transcription factor 1 |
| | RAB6B | RAB6B, member RAS oncogene family |
| | DCUN1D1 | defective in cullin neddylation 1 domain containing 1 |
| | TACR1 | tachykinin receptor 1 |
| | FERMT3 | fermitin family member 3 |
| | ZC3H15 | zinc finger CCCH-type containing 15 |
| | RGS9BP | regulator of G protein signaling 9 binding protein |
| | DNAJC8 | DnaJ heat shock protein family (Hsp40) member C8 |
| | NDP | NDP, norrin cystine knot growth factor |
| | ETV1 | ETS variant 1 |
| | LRRC8C | leucine rich repeat containing 8 VRAC subunit C |
| | MRPS10 | mitochondrial ribosomal protein S10 |
| | HDAC9 | histone deacetylase 9 |
| | PRRG3 | proline rich and Gla domain 3 |
| | GABRG1 | gamma-aminobutyric acid type A receptor gamma1 subunit |
| | ARHGAP31 | Rho GTPase activating protein 31 |
| | TMEM87B | transmembrane protein 87B |
| | DGKB | diacylglycerol kinase beta |

| | | |
|---|---|---|
| | MORC1 | MORC family CW-type zinc finger 1 |
| | NOL9 | nucleolar protein 9 |
| | RAB27B | RAB27B, member RAS oncogene family |
| | LRRC66 | leucine rich repeat containing 66 |
| | PGBD5 | piggyBac transposable element derived 5 |
| | ESCO1 | establishment of sister chromatid cohesion N-acetyltransferase 1 |
| | LYRM2 | LYR motif containing 2 |
| | SEMA3E | semaphorin 3E |
| | ATP5PF | ATP synthase peripheral stalk subunit F6 |
| | WDR36 | WD repeat domain 36 |
| | ERH | ERH, mRNA splicing and mitosis factor |
| | CDC37L1 | cell division cycle 37 like 1 |
| | TNKS2 | tankyrase 2 |
| | TBX21 | T-box 21 |
| | FUT2 | fucosyltransferase 2 |
| | SLC41A2 | solute carrier family 41 member 2 |
| | C1QTNF1 | C1q and TNF related 1 |
| | CIRBP | cold inducible RNA binding protein |
| | HLTF | helicase like transcription factor |
| | INSR | insulin receptor |
| | MAT2B | methionine adenosyltransferase 2B |
| | MIS18BP1 | MIS18 binding protein 1 |
| | MCFD2 | multiple coagulation factor deficiency 2 |
| | SUPT3H | SPT3 homolog, SAGA and STAGA complex component |
| | ZNF326 | zinc finger protein 326 |
| | KCNB1 | potassium voltage-gated channel subfamily B member 1 |
| | POU3F2 | POU class 3 homeobox 2 |
| | MAP2K6 | mitogen-activated protein kinase kinase 6 |
| | DPP10 | dipeptidyl peptidase like 10 |
| | WWOX | WW domain containing oxidoreductase |
| | CARF | calcium responsive transcription factor |
| | SPON1 | spondin 1 |
| | PCSK5 | proprotein convertase subtilisin/kexin type 5 |
| | SLC20A1 | solute carrier family 20 member 1 |
| | SLC27A1 | solute carrier family 27 member 1 |
| | GSPT2 | G1 to S phase transition 2 |
| | NSL1 | NSL1, MIS12 kinetochore complex component |
| | PAIP2B | poly(A) binding protein interacting protein 2B |
| | CALM2 | calmodulin 2 |
| | ZC3H18 | zinc finger CCCH-type containing 18 |
| | SOX1 | SRY-box 1 |
| | SCN3A | sodium voltage-gated channel alpha subunit 3 |

| | | |
|---|---|---|
| | GNRHR | gonadotropin releasing hormone receptor |
| | LHFPL4 | LHFPL tetraspan subfamily member 4 |
| | MINDY2 | MINDY lysine 48 deubiquitinase 2 |
| | IDS | iduronate 2-sulfatase |
| | TRIP12 | thyroid hormone receptor interactor 12 |
| | DMGDH | dimethylglycine dehydrogenase |
| | APP | amyloid beta precursor protein |
| | MDM1 | Mdm1 nuclear protein |
| | RAN | RAN, member RAS oncogene family |
| | SYT11 | synaptotagmin 11 |
| | AR | androgen receptor |
| | UNC5D | unc-5 netrin receptor D |
| | ST14 | suppression of tumorigenicity 14 |
| | RAB3D | RAB3D, member RAS oncogene family |
| | SEC24B | SEC24 homolog B, COPII coat complex component |
| | PDP1 | pyruvate dehyrogenase phosphatase catalytic subunit 1 |
| | PTPRC | protein tyrosine phosphatase, receptor type C |
| | DCAF7 | DDB1 and CUL4 associated factor 7 |
| | HNRNPR | heterogeneous nuclear ribonucleoprotein R |
| | ARHGAP19 | Rho GTPase activating protein 19 |
| | EPB41L2 | erythrocyte membrane protein band 4.1 like 2 |
| | COL5A2 | collagen type V alpha 2 chain |
| | KRR1 | KRR1, small subunit processome component homolog |
| | FOXC1 | forkhead box C1 |
| | PRKCE | protein kinase C epsilon |
| | SOX21 | SRY-box 21 |
| | CDKN1B | cyclin dependent kinase inhibitor 1B |
| | RNPEP | arginyl aminopeptidase |
| | NOTCH3 | notch 3 |
| | BCAT1 | branched chain amino acid transaminase 1 |
| | EYA1 | EYA transcriptional coactivator and phosphatase 1 |
| | TMEM71 | transmembrane protein 71 |
| | SEZ6L | seizure related 6 homolog like |
| | VDAC1 | voltage dependent anion channel 1 |
| | TSPYL5 | TSPY like 5 |
| | AAGAB | alpha and gamma adaptin binding protein |
| | PARPBP | PARP1 binding protein |
| | FAM122B | family with sequence similarity 122B |
| | COL5A1 | collagen type V alpha 1 chain |
| | CMPK2 | cytidine/uridine monophosphate kinase 2 |
| | ACTR3 | ARP3 actin related protein 3 homolog |
| | SLC35C2 | solute carrier family 35 member C2 |

| | | |
|---|---|---|
| | HAND2 | heart and neural crest derivatives expressed 2 |
| | PARM1 | prostate androgen-regulated mucin-like protein 1 |
| | MPDZ | multiple PDZ domain crumbs cell polarity complex component |
| | IFFO2 | intermediate filament family orphan 2 |
| | CBX1 | chromobox 1 |
| | RBM33 | RNA binding motif protein 33 |
| | CAVIN2 | caveolae associated protein 2 |
| | ADGRG1 | adhesion G protein-coupled receptor G1 |
| | TMEM196 | transmembrane protein 196 |
| | MCF2L | MCF.2 cell line derived transforming sequence like |
| | PARVA | parvin alpha |
| | ETFBKMT | electron transfer flavoprotein subunit beta lysine methyltransferase |
| | NXPE3 | neurexophilin and PC-esterase domain family member 3 |
| | ZNF780B | zinc finger protein 780B |
| | ANKRD45 | ankyrin repeat domain 45 |
| | IFIT5 | interferon induced protein with tetratricopeptide repeats 5 |
| | PLS1 | plastin 1 |
| | PURA | purine rich element binding protein A |
| | PDK1 | pyruvate dehydrogenase kinase 1 |
| | RCHY1 | ring finger and CHY zinc finger domain containing 1 |
| | PUM1 | pumilio RNA binding family member 1 |
| | ERMN | ermin |
| | MYCBP | MYC binding protein |
| | ZFAND3 | zinc finger AN1-type containing 3 |
| | DCBLD2 | discoidin, CUB and LCCL domain containing 2 |
| | ACSL6 | acyl-CoA synthetase long chain family member 6 |
| | RCSD1 | RCSD domain containing 1 |
| | OPA3 | OPA3, outer mitochondrial membrane lipid metabolism regulator |
| | ARSK | arylsulfatase family member K |
| | NCR1 | natural cytotoxicity triggering receptor 1 |
| | ZFP91 | ZFP91 zinc finger protein |
| | PTP4A1 | protein tyrosine phosphatase type IVA, member 1 |
| | EZH1 | enhancer of zeste 1 polycomb repressive complex 2 subunit |
| | C2CD2 | C2 calcium dependent domain containing 2 |
| | MSR1 | macrophage scavenger receptor 1 |
| | PIGN | phosphatidylinositol glycan anchor biosynthesis class N |
| | MXRA7 | matrix remodeling associated 7 |
| | NRK | Nik related kinase |
| | PRDM2 | PR/SET domain 2 |
| | PATE4 | prostate and testis expressed 4 |
| | NAP1L3 | nucleosome assembly protein 1 like 3 |

| | | |
|---|---|---|
| | SOCS3 | suppressor of cytokine signaling 3 |
| | ENPP1 | ectonucleotide pyrophosphatase/phosphodiesterase 1 |
| | NET1 | neuroepithelial cell transforming 1 |
| | ZCCHC14 | zinc finger CCHC-type containing 14 |
| | TCF24 | transcription factor 24 |
| | PPP2R5E | protein phosphatase 2 regulatory subunit B'epsilon |
| | ACVR2A | activin A receptor type 2A |
| | NRIP1 | nuclear receptor interacting protein 1 |
| | PANK1 | pantothenate kinase 1 |
| | DOCK1 | dedicator of cytokinesis 1 |
| Segment-2-PB1- MR17 3P | MAP3K2 | mitogen-activated protein kinase kinase kinase 2 |
| | ABCC9 | ATP binding cassette subfamily C member 9 |
| | VEGFA | vascular endothelial growth factor A |
| | TULP3 | tubby like protein 3 |
| | AHR | aryl hydrocarbon receptor |
| | BCL6 | BCL6, transcription repressor |
| | OSBP2 | oxysterol binding protein 2 |
| | CRISPLD1 | cysteine rich secretory protein LCCL domain containing 1 |
| | PCSK2 | proprotein convertase subtilisin/kexin type 2 |
| | ROCK2 | Rho associated coiled-coil containing protein kinase 2 |
| | SLC24A2 | solute carrier family 24 member 2 |
| | MEF2C | myocyte enhancer factor 2C |
| | ATL2 | atlastin GTPase 2 |
| | LIN7A | lin-7 homolog A, crumbs cell polarity complex component |
| | C5orf24 | chromosome 5 open reading frame 24 |
| | GRIA2 | glutamate ionotropic receptor AMPA type subunit 2 |
| | MARCHF7 | membrane associated ring-CH-type finger 7 |
| | SUZ12 | SUZ12, polycomb repressive complex 2 subunit |
| | TRDN | triadin |
| | CACNB4 | calcium voltage-gated channel auxiliary subunit beta 4 |
| | OTUD4 | OTU deubiquitinase 4 |
| | PRG4 | proteoglycan 4 |
| | CYSLTR1 | cysteinyl leukotriene receptor 1 |
| | LYRM7 | LYR motif containing 7 |
| | SNX1 | sorting nexin 1 |
| | VANGL2 | VANGL planar cell polarity protein 2 |
| | NEK7 | NIMA related kinase 7 |
| | PLPP3 | phospholipid phosphatase 3 |
| | NPAS2 | neuronal PAS domain protein 2 |
| | HDAC1 | histone deacetylase 1 |
| | RAB11FIP2 | RAB11 family interacting protein 2 |

| | | |
|---|---|---|
| | MINDY2 | MINDY lysine 48 deubiquitinase 2 |
| | TMEM178B | transmembrane protein 178B |
| | ATP10A | ATPase phospholipid transporting 10A (putative) |
| | ACKR3 | atypical chemokine receptor 3 |
| | GRHL1 | grainyhead like transcription factor 1 |
| | KPNA4 | karyopherin subunit alpha 4 |
| | NECAB1 | N-terminal EF-hand calcium binding protein 1 |
| | CREBZF | CREB/ATF bZIP transcription factor |
| | LOX | lysyl oxidase |
| | SEMA3F | semaphorin 3F |
| | SLC33A1 | solute carrier family 33 member 1 |
| | NFAT5 | nuclear factor of activated T cells 5 |
| | POGZ | pogo transposable element derived with ZNF domain |
| | SLC30A5 | solute carrier family 30 member 5 |
| | MPP6 | membrane palmitoylated protein 6 |
| | SGCD | sarcoglycan delta |
| | ACSL6 | acyl-CoA synthetase long chain family member 6 |
| | SENP3 | SUMO specific peptidase 3 |
| | LOXL3 | lysyl oxidase like 3 |
| | ACBD5 | acyl-CoA binding domain containing 5 |
| | CCAR1 | cell division cycle and apoptosis regulator 1 |
| | IQSEC1 | IQ motif and Sec7 domain 1 |
| | ARHGEF11 | Rho guanine nucleotide exchange factor 11 |
| | SETD3 | SET domain containing 3, actin histidine methyltransferase |
| | EP300 | E1A binding protein p300 |
| | TMEM33 | transmembrane protein 33 |
| | MKRN3 | makorin ring finger protein 3 |
| | SGPP1 | sphingosine-1-phosphate phosphatase 1 |
| | CDC73 | cell division cycle 73 |
| | PIK3CB | phosphatidylinositol-4,5-bisphosphate 3-kinase catalytic subunit beta |
| | PYROXD1 | pyridine nucleotide-disulphide oxidoreductase domain 1 |
| | CCDC88A | coiled-coil domain containing 88A |
| | ZNF655 | zinc finger protein 655 |
| | PCNX1 | pecanex 1 |
| | HIPK3 | homeodomain interacting protein kinase 3 |
| | TVP23B | trans-golgi network vesicle protein 23 homolog B |
| | ZBTB18 | zinc finger and BTB domain containing 18 |
| | GTPBP10 | GTP binding protein 10 |
| | HMCN1 | hemicentin 1 |
| | ZNF208 | zinc finger protein 208 |
| | XPOT | exportin for tRNA |

| | | |
|---|---|---|
| | RNF19A | ring finger protein 19A, RBR E3 ubiquitin protein ligase |
| | CHRNA5 | cholinergic receptor nicotinic alpha 5 subunit |
| | ZBTB6 | zinc finger and BTB domain containing 6 |
| | KIAA1217 | KIAA1217 |
| | BCL9L | BCL9 like |
| | SPEN | spen family transcriptional repressor |
| | KAT6A | lysine acetyltransferase 6A |
| | MICU3 | mitochondrial calcium uptake family member 3 |
| | PTPRK | protein tyrosine phosphatase, receptor type K |
| | LRP6 | LDL receptor related protein 6 |
| | FAM126B | family with sequence similarity 126 member B |
| | ZYG11B | zyg-11 family member B, cell cycle regulator |
| | RNF170 | ring finger protein 170 |
| | MOSMO | modulator of smoothened |
| | CLEC2D | C-type lectin domain family 2 member D |
| | MGAT4A | alpha-1,3-mannosyl-glycoprotein 4-beta-N-acetylglucosaminyltransferase A |
| | KHDRBS1 | KH RNA binding domain containing, signal transduction associated 1 |
| | CEP97 | centrosomal protein 97 |
| | C11orf54 | chromosome 11 open reading frame 54 |
| | MAP9 | microtubule associated protein 9 |
| | PMEPA1 | prostate transmembrane protein, androgen induced 1 |
| | TWSG1 | twisted gastrulation BMP signaling modulator 1 |
| | CNIH1 | cornichon family AMPA receptor auxiliary protein 1 |
| | PCGF3 | polycomb group ring finger 3 |
| | EEA1 | early endosome antigen 1 |
| | PCDH10 | protocadherin 10 |
| | TNRC18 | trinucleotide repeat containing 18 |
| | AEBP2 | AE binding protein 2 |
| | SCUBE1 | signal peptide, CUB domain and EGF like domain containing 1 |
| | LPCAT4 | lysophosphatidylcholine acyltransferase 4 |
| | SMAD7 | SMAD family member 7 |
| | PRNP | prion protein |
| | PAPPA | pappalysin 1 |
| | ZBTB41 | zinc finger and BTB domain containing 41 |
| | DLG1 | discs large MAGUK scaffold protein 1 |
| | TFAM | transcription factor A, mitochondrial |
| | GOLPH3 | golgi phosphoprotein 3 |
| | DNA2 | DNA replication helicase/nuclease 2 |
| | TMPRSS11B | transmembrane serine protease 11B |
| | MBNL3 | muscleblind like splicing regulator 3 |
| | INPP5A | inositol polyphosphate-5-phosphatase A |

| | | |
|---|---|---|
| | FIGN | fidgetin, microtubule severing factor |
| | NAA15 | N(alpha)-acetyltransferase 15, NatA auxiliary subunit |
| | PABPC1 | poly(A) binding protein cytoplasmic 1 |
| | UGCG | UDP-glucose ceramide glucosyltransferase |
| | FRMPD4 | FERM and PDZ domain containing 4 |
| | BOD1L1 | biorientation of chromosomes in cell division 1 like 1 |
| | PRDM5 | PR/SET domain 5 |
| | SPRY3 | sprouty RTK signaling antagonist 3 |
| | SLC4A7 | solute carrier family 4 member 7 |
| | NEGR1 | neuronal growth regulator 1 |
| | IKZF2 | IKAROS family zinc finger 2 |
| | GABRA4 | gamma-aminobutyric acid type A receptor alpha4 subunit |
| | PDS5B | PDS5 cohesin associated factor B |
| | RYBP | RING1 and YY1 binding protein |
| | ELAVL4 | ELAV like RNA binding protein 4 |
| | BTBD1 | BTB domain containing 1 |
| | ATP8B2 | ATPase phospholipid transporting 8B2 |
| | CACNA2D1 | calcium voltage-gated channel auxiliary subunit alpha2delta 1 |
| | CADM2 | cell adhesion molecule 2 |
| | DIO2 | iodothyronine deiodinase 2 |
| | EXTL2 | exostosin like glycosyltransferase 2 |
| | RMC1 | regulator of MON1-CCZ1 |
| | ERBIN | erbb2 interacting protein |
| | CNTLN | centlein |
| | SLMAP | sarcolemma associated protein |
| | GPR155 | G protein-coupled receptor 155 |
| | PSMC3IP | PSMC3 interacting protein |
| | WNT5B | Wnt family member 5B |
| | KIAA0825 | KIAA0825 |
| | PTGER4 | prostaglandin E receptor 4 |
| | CFLAR | CASP8 and FADD like apoptosis regulator |
| | LRP1B | LDL receptor related protein 1B |
| | NR4A3 | nuclear receptor subfamily 4 group A member 3 |
| | CHD1 | chromodomain helicase DNA binding protein 1 |
| | PPIL4 | peptidylprolyl isomerase like 4 |
| | OPA1 | OPA1, mitochondrial dynamin like GTPase |
| | TET1 | tet methylcytosine dioxygenase 1 |
| | PI15 | peptidase inhibitor 15 |
| | PRDM11 | PR/SET domain 11 |
| | MYCBP2 | MYC binding protein 2, E3 ubiquitin protein ligase |
| | ENTPD1 | ectonucleoside triphosphate diphosphohydrolase 1 |
| | SPOP | speckle type BTB/POZ protein |

| | | |
|---|---|---|
| | USP42 | ubiquitin specific peptidase 42 |
| | CALCRL | calcitonin receptor like receptor |
| | RFX7 | regulatory factor X7 |
| | LGR4 | leucine rich repeat containing G protein-coupled receptor 4 |
| | HTRA2 | HtrA serine peptidase 2 |
| | SMCHD1 | structural maintenance of chromosomes flexible hinge domain containing 1 |
| | ASNSD1 | asparagine synthetase domain containing 1 |
| | RNF144A | ring finger protein 144A |
| | ANKRD17 | ankyrin repeat domain 17 |
| | TCF4 | transcription factor 4 |
| | KIAA0355 | KIAA0355 |
| | NOL7 | nucleolar protein 7 |
| | ZSWIM2 | zinc finger SWIM-type containing 2 |
| | DLC1 | DLC1 Rho GTPase activating protein |
| | COL4A4 | collagen type IV alpha 4 chain |
| | SLK | STE20 like kinase |
| | ZFAND5 | zinc finger AN1-type containing 5 |
| | FAM122B | family with sequence similarity 122B |
| | ADAMTS5 | ADAM metallopeptidase with thrombospondin type 1 motif 5 |
| | KCNT2 | potassium sodium-activated channel subfamily T member 2 |
| | LMO7 | LIM domain 7 |
| | CABP7 | calcium binding protein 7 |
| | RABGAP1 | RAB GTPase activating protein 1 |
| | PPP3CC | protein phosphatase 3 catalytic subunit gamma |
| | TMTC1 | transmembrane and tetratricopeptide repeat containing 1 |
| | ETNK1 | ethanolamine kinase 1 |
| | NEXMIF | neurite extension and migration factor |
| | NRXN1 | neurexin 1 |
| | DCP1B | decapping mRNA 1B |
| | JRKL | JRK like |
| | PLXDC2 | plexin domain containing 2 |
| | NFIB | nuclear factor I B |
| | PRRC2C | proline rich coiled-coil 2C |
| | CDC14A | cell division cycle 14A |
| | VPS54 | VPS54, GARP complex subunit |
| | TRHDE | thyrotropin releasing hormone degrading enzyme |
| | MYO1E | myosin IE |
| | RICTOR | RPTOR independent companion of MTOR complex 2 |
| | EXOC7 | exocyst complex component 7 |
| | FAM117B | family with sequence similarity 117 member B |
| | SHOC2 | SHOC2, leucine rich repeat scaffold protein |

| | | |
|---|---|---|
| | CXCR4 | C-X-C motif chemokine receptor 4 |
| | OSBPL6 | oxysterol binding protein like 6 |
| | MKX | mohawk homeobox |
| | FZD3 | frizzled class receptor 3 |
| | CEP126 | centrosomal protein 126 |
| | NTF3 | neurotrophin 3 |
| | TNFRSF11B | TNF receptor superfamily member 11b |
| | PLCL1 | phospholipase C like 1 (inactive) |
| | IPO7 | importin 7 |
| | BRWD3 | bromodomain and WD repeat domain containing 3 |
| | KLHL5 | kelch like family member 5 |
| | NFATC3 | nuclear factor of activated T cells 3 |
| | ELP4 | elongator acetyltransferase complex subunit 4 |
| | PHTF2 | putative homeodomain transcription factor 2 |
| | ATP11C | ATPase phospholipid transporting 11C |
| | TBX4 | T-box 4 |
| | LIN7C | lin-7 homolog C, crumbs cell polarity complex component |
| | KIAA1211 | KIAA1211 |
| | CPEB3 | cytoplasmic polyadenylation element binding protein 3 |
| | EPS8 | epidermal growth factor receptor pathway substrate 8 |
| | FH | fumarate hydratase |
| | PWWP3B | PWWP domain containing 3B |
| | TRPM6 | transient receptor potential cation channel subfamily M member 6 |
| | PHF12 | PHD finger protein 12 |
| | SASS6 | SAS-6 centriolar assembly protein |
| | PIGW | phosphatidylinositol glycan anchor biosynthesis class W |
| | PURA | purine rich element binding protein A |
| | CBX3 | chromobox 3 |
| | CACNA1G | calcium voltage-gated channel subunit alpha1 G |
| | MAGI1 | membrane associated guanylate kinase, WW and PDZ domain containing 1 |
| | DUSP10 | dual specificity phosphatase 10 |
| | CAAP1 | caspase activity and apoptosis inhibitor 1 |
| | ADCYAP1 | adenylate cyclase activating polypeptide 1 |
| | TMEM170B | transmembrane protein 170B |
| | L3MBTL3 | L3MBTL3, histone methyl-lysine binding protein |
| | GPCPD1 | glycerophosphocholine phosphodiesterase 1 |
| | COL4A5 | collagen type IV alpha 5 chain |
| | MAD2L1BP | MAD2L1 binding protein |
| | BTBD7 | BTB domain containing 7 |
| | AKAP7 | A-kinase anchoring protein 7 |
| | DCUN1D1 | defective in cullin neddylation 1 domain containing 1 |

| | | |
|---|---|---|
| | SELENOF | selenoprotein F |
| | CCDC50 | coiled-coil domain containing 50 |
| | MDK | midkine |
| | NAALAD2 | N-acetylated alpha-linked acidic dipeptidase 2 |
| | ATF2 | activating transcription factor 2 |
| | UBE2E1 | ubiquitin conjugating enzyme E2 E1 |
| | KAT6B | lysine acetyltransferase 6B |
| | LIN54 | lin-54 DREAM MuvB core complex component |
| | NBEA | neurobeachin |
| | PHIP | pleckstrin homology domain interacting protein |
| | LZTS3 | leucine zipper tumor suppressor family member 3 |
| | QKI | QKI, KH domain containing RNA binding |
| | C21orf91 | chromosome 21 open reading frame 91 |
| | PUM3 | pumilio RNA binding family member 3 |
| | RSPO3 | R-spondin 3 |
| | RBMS3 | RNA binding motif single stranded interacting protein 3 |
| | RNF128 | ring finger protein 128, E3 ubiquitin protein ligase |
| | SCAI | suppressor of cancer cell invasion |
| | IDE | insulin degrading enzyme |
| | ZNF626 | zinc finger protein 626 |
| | CFAP97 | cilia and flagella associated protein 97 |
| | BRIP1 | BRCA1 interacting protein C-terminal helicase 1 |
| | SGIP1 | SH3 domain GRB2 like endophilin interacting protein 1 |
| | RANBP9 | RAN binding protein 9 |
| | PLA2R1 | phospholipase A2 receptor 1 |
| | USP24 | ubiquitin specific peptidase 24 |
| | UCHL5 | ubiquitin C-terminal hydrolase L5 |
| | DEK | DEK proto-oncogene |
| | SHPRH | SNF2 histone linker PHD RING helicase |
| | PHYHIPL | phytanoyl-CoA 2-hydroxylase interacting protein like |
| | BCLAF1 | BCL2 associated transcription factor 1 |
| | BAZ2B | bromodomain adjacent to zinc finger domain 2B |
| | REEP3 | receptor accessory protein 3 |
| | ERCC3 | ERCC excision repair 3, TFIIH core complex helicase subunit |
| | SP4 | Sp4 transcription factor |
| | NETO1 | neuropilin and tolloid like 1 |
| | CNOT6L | CCR4-NOT transcription complex subunit 6 like |
| | IRF1 | interferon regulatory factor 1 |
| | WDR17 | WD repeat domain 17 |
| | PRPF40A | pre-mRNA processing factor 40 homolog A |
| | RNF141 | ring finger protein 141 |
| | AKTIP | AKT interacting protein |

| | | |
|---|---|---|
| | MCMDC2 | minichromosome maintenance domain containing 2 |
| | SLC38A11 | solute carrier family 38 member 11 |
| | FOXF1 | forkhead box F1 |
| | RUFY3 | RUN and FYVE domain containing 3 |
| | SLAIN2 | SLAIN motif family member 2 |
| | MAP4K4 | mitogen-activated protein kinase kinase kinase kinase 4 |
| | PTPN3 | protein tyrosine phosphatase, non-receptor type 3 |
| | PJA2 | praja ring finger ubiquitin ligase 2 |
| | NOM1 | nucleolar protein with MIF4G domain 1 |
| | NAA25 | N(alpha)-acetyltransferase 25, NatB auxiliary subunit |
| | RALYL | RALY RNA binding protein like |
| | TMEM18 | transmembrane protein 18 |
| | CHL1 | cell adhesion molecule L1 like |
| | MBLAC2 | metallo-beta-lactamase domain containing 2 |
| | ZZZ3 | zinc finger ZZ-type containing 3 |
| | MYRF | myelin regulatory factor |
| | FEM1B | fem-1 homolog B |
| | EPS15L1 | epidermal growth factor receptor pathway substrate 15 like 1 |
| | ITGBL1 | integrin subunit beta like 1 |
| | LSM8 | LSM8 homolog, U6 small nuclear RNA associated |
| | GABRB3 | gamma-aminobutyric acid type A receptor beta3 subunit |
| | SLC17A6 | solute carrier family 17 member 6 |
| | EBI3 | Epstein-Barr virus induced 3 |
| | SYNPO2 | synaptopodin 2 |
| | STXBP1 | syntaxin binding protein 1 |
| | DNAL4 | dynein axonemal light chain 4 |
| | FNIP2 | folliculin interacting protein 2 |
| | GRAMD1B | GRAM domain containing 1B |
| | PTPRJ | protein tyrosine phosphatase, receptor type J |
| | CNOT4 | CCR4-NOT transcription complex subunit 4 |
| | NRARP | NOTCH regulated ankyrin repeat protein |
| | SATB1 | SATB homeobox 1 |
| | MRPL57 | mitochondrial ribosomal protein L57 |
| | CHORDC1 | cysteine and histidine rich domain containing 1 |
| | HSD17B7 | hydroxysteroid 17-beta dehydrogenase 7 |
| | BCAT1 | branched chain amino acid transaminase 1 |
| | MANBA | mannosidase beta |
| | ASPA | aspartoacylase |
| | DIP2C | disco interacting protein 2 homolog C |
| | AJAP1 | adherens junctions associated protein 1 |
| | ATP2B4 | ATPase plasma membrane Ca2+ transporting 4 |
| | FRAS1 | Fraser extracellular matrix complex subunit 1 |

| | | |
|---|---|---|
| | GPN3 | GPN-loop GTPase 3 |
| | DNAJB4 | DnaJ heat shock protein family (Hsp40) member B4 |
| | TMED5 | transmembrane p24 trafficking protein 5 |
| | RPS6KA5 | ribosomal protein S6 kinase A5 |
| | CCL18 | C-C motif chemokine ligand 18 |
| | ASB7 | ankyrin repeat and SOCS box containing 7 |
| | KCNV1 | potassium voltage-gated channel modifier subfamily V member 1 |
| | ATN1 | atrophin 1 |
| | F9 | coagulation factor IX |
| | ODR4 | odr-4 GPCR localization factor homolog |
| | FAT3 | FAT atypical cadherin 3 |
| | C11orf87 | chromosome 11 open reading frame 87 |
| | RAB3C | RAB3C, member RAS oncogene family |
| | EN2 | engrailed homeobox 2 |
| | MEF2A | myocyte enhancer factor 2A |
| | FAXC | failed axon connections homolog |
| | ZC4H2 | zinc finger C4H2-type containing |
| | NCK1 | NCK adaptor protein 1 |
| | OSMR | oncostatin M receptor |
| | FBXW7 | F-box and WD repeat domain containing 7 |
| | TPK1 | thiamin pyrophosphokinase 1 |
| | PIAS1 | protein inhibitor of activated STAT 1 |
| | HAUS2 | HAUS augmin like complex subunit 2 |
| | KDM4C | lysine demethylase 4C |
| | SLC6A1 | solute carrier family 6 member 1 |
| | VCAN | versican |
| | PRR27 | proline rich 27 |
| | FGF7 | fibroblast growth factor 7 |
| | BCAP29 | B cell receptor associated protein 29 |
| | PRRG4 | proline rich and Gla domain 4 |
| | BPTF | bromodomain PHD finger transcription factor |
| | ARMC8 | armadillo repeat containing 8 |
| | SPAG9 | sperm associated antigen 9 |
| | CREBRF | CREB3 regulatory factor |
| | CSPP1 | centrosome and spindle pole associated protein 1 |
| | WNT11 | Wnt family member 11 |
| | RAB27B | RAB27B, member RAS oncogene family |
| | SKIL | SKI like proto-oncogene |
| | ZNF384 | zinc finger protein 384 |
| | UBE2U | ubiquitin conjugating enzyme E2 U |
| | DENND4A | DENN domain containing 4A |

| | | |
|---|---|---|
| | WDR73 | WD repeat domain 73 |
| | ZBTB14 | zinc finger and BTB domain containing 14 |
| | DCUN1D3 | defective in cullin neddylation 1 domain containing 3 |
| | CBX6 | chromobox 6 |
| | RSRC2 | arginine and serine rich coiled-coil 2 |
| | ARAF | A-Raf proto-oncogene, serine/threonine kinase |
| | QSER1 | glutamine and serine rich 1 |
| | FABP2 | fatty acid binding protein 2 |
| | RBM41 | RNA binding motif protein 41 |
| | DLL1 | delta like canonical Notch ligand 1 |
| | OSBPL3 | oxysterol binding protein like 3 |
| | GSK3B | glycogen synthase kinase 3 beta |
| | MKRN2OS | MKRN2 opposite strand |
| | DACT1 | dishevelled binding antagonist of beta catenin 1 |
| | LEMD3 | LEM domain containing 3 |
| | RPGRIP1L | RPGRIP1 like |
| | CHST2 | carbohydrate sulfotransferase 2 |
| | WNT5A | Wnt family member 5A |
| | ASF1A | anti-silencing function 1A histone chaperone |
| | HOOK1 | hook microtubule tethering protein 1 |
| | CFAP44 | cilia and flagella associated protein 44 |
| | ZNF366 | zinc finger protein 366 |
| | BACH2 | BTB domain and CNC homolog 2 |
| | SNTG1 | syntrophin gamma 1 |
| | WDR92 | WD repeat domain 92 |
| | AVPR1A | arginine vasopressin receptor 1A |
| | RPS6KB1 | ribosomal protein S6 kinase B1 |
| | RNF6 | ring finger protein 6 |
| | NR1D2 | nuclear receptor subfamily 1 group D member 2 |
| | RAB9B | RAB9B, member RAS oncogene family |
| | AHCTF1 | AT-hook containing transcription factor 1 |
| | ESRP2 | epithelial splicing regulatory protein 2 |
| | LCOR | ligand dependent nuclear receptor corepressor |
| | FUT9 | fucosyltransferase 9 |
| | CD2AP | CD2 associated protein |
| | ABCE1 | ATP binding cassette subfamily E member 1 |
| | SCN9A | sodium voltage-gated channel alpha subunit 9 |
| | TBC1D4 | TBC1 domain family member 4 |
| | PCDH8 | protocadherin 8 |
| | TMEM87A | transmembrane protein 87A |
| | SIKE1 | suppressor of IKBKE 1 |
| | GUCY1A2 | guanylate cyclase 1 soluble subunit alpha 2 |

| | | |
|---|---|---|
| | TRIO | trio Rho guanine nucleotide exchange factor |
| | PHACTR4 | phosphatase and actin regulator 4 |
| | RAB30 | RAB30, member RAS oncogene family |
| | DDOST | dolichyl-diphosphooligosaccharide--protein glycosyltransferase non-catalytic subunit |
| | R3HDM2 | R3H domain containing 2 |
| | KCNQ3 | potassium voltage-gated channel subfamily Q member 3 |
| | PREX2 | phosphatidylinositol-3,4,5-trisphosphate dependent Rac exchange factor 2 |
| | RIOK2 | RIO kinase 2 |
| | PTEN | phosphatase and tensin homolog |
| | CDK13 | cyclin dependent kinase 13 |
| | PGM2L1 | phosphoglucomutase 2 like 1 |
| | PTPRZ1 | protein tyrosine phosphatase, receptor type Z1 |
| | NEUROD1 | neuronal differentiation 1 |
| | GFM2 | G elongation factor mitochondrial 2 |
| | PER2 | period circadian regulator 2 |
| | FAM118B | family with sequence similarity 118 member B |
| | PPP1R1C | protein phosphatase 1 regulatory inhibitor subunit 1C |
| | TMEM47 | transmembrane protein 47 |
| | ANKRD50 | ankyrin repeat domain 50 |
| | N4BP1 | NEDD4 binding protein 1 |
| | SLC40A1 | solute carrier family 40 member 1 |
| | CALN1 | calneuron 1 |
| | HERC3 | HECT and RLD domain containing E3 ubiquitin protein ligase 3 |
| | FAM177A1 | family with sequence similarity 177 member A1 |
| | RNF44 | ring finger protein 44 |
| | HAPLN1 | hyaluronan and proteoglycan link protein 1 |
| | RIPK2 | receptor interacting serine/threonine kinase 2 |
| | FAM19A4 | family with sequence similarity 19 member A4, C-C motif chemokine like |
| | MNX1 | motor neuron and pancreas homeobox 1 |
| | ABCD2 | ATP binding cassette subfamily D member 2 |
| | TMEM38B | transmembrane protein 38B |
| | NRF1 | nuclear respiratory factor 1 |
| | ZNF333 | zinc finger protein 333 |
| | HBS1L | HBS1 like translational GTPase |
| | PRAG1 | PEAK1 related, kinase-activating pseudokinase 1 |
| | PARD6G | par-6 family cell polarity regulator gamma |
| | ONECUT2 | one cut homeobox 2 |
| | SLC35F1 | solute carrier family 35 member F1 |
| | ARHGEF38 | Rho guanine nucleotide exchange factor 38 |
| | BAZ1A | bromodomain adjacent to zinc finger domain 1A |

| | | |
|---|---|---|
| | C8orf34 | chromosome 8 open reading frame 34 |
| | C1orf56 | chromosome 1 open reading frame 56 |
| | TBC1D23 | TBC1 domain family member 23 |
| | PTER | phosphotriesterase related |
| | SNRNP48 | small nuclear ribonucleoprotein U11/U12 subunit 48 |
| | FAM131A | family with sequence similarity 131 member A |
| | TRMT10A | tRNA methyltransferase 10A |
| | GABRP | gamma-aminobutyric acid type A receptor pi subunit |
| | ZKSCAN4 | zinc finger with KRAB and SCAN domains 4 |
| | EIF4G1 | eukaryotic translation initiation factor 4 gamma 1 |
| | AGMO | alkylglycerol monooxygenase |
| | AP1S1 | adaptor related protein complex 1 subunit sigma 1 |
| | PAH | phenylalanine hydroxylase |
| | SLITRK4 | SLIT and NTRK like family member 4 |
| | TFDP1 | transcription factor Dp-1 |
| | NDUFAF4 | NADH:ubiquinone oxidoreductase complex assembly factor 4 |
| | FBRS | fibrosin |
| | PLG | plasminogen |
| | C6orf120 | chromosome 6 open reading frame 120 |
| | ZNF454 | zinc finger protein 454 |
| | GAN | gigaxonin |
| | ILDR1 | immunoglobulin like domain containing receptor 1 |
| | GRTP1 | growth hormone regulated TBC protein 1 |
| | ARFGEF3 | ARFGEF family member 3 |
| | ANKRD39 | ankyrin repeat domain 39 |
| | GPR180 | G protein-coupled receptor 180 |
| | SPRED1 | sprouty related EVH1 domain containing 1 |
| | SRI | sorcin |
| | HCN1 | hyperpolarization activated cyclic nucleotide gated potassium channel 1 |
| | FAM19A2 | family with sequence similarity 19 member A2, C-C motif chemokine like |
| | DENND5B | DENN domain containing 5B |
| | ZNF737 | zinc finger protein 737 |
| | TNPO1 | transportin 1 |
| | MARK1 | microtubule affinity regulating kinase 1 |
| | ARAP1 | ArfGAP with RhoGAP domain, ankyrin repeat and PH domain 1 |
| | PWWP2A | PWWP domain containing 2A |
| | TRIM22 | tripartite motif containing 22 |
| | LMO4 | LIM domain only 4 |
| | UBAP2L | ubiquitin associated protein 2 like |
| | PTPRD | protein tyrosine phosphatase, receptor type D |

| | | |
|---|---|---|
| | EIF2AK3 | eukaryotic translation initiation factor 2 alpha kinase 3 |
| | ZFAND3 | zinc finger AN1-type containing 3 |
| | KREMEN1 | kringle containing transmembrane protein 1 |
| | GRIA4 | glutamate ionotropic receptor AMPA type subunit 4 |
| | COMMD2 | COMM domain containing 2 |
| | PLSCR4 | phospholipid scramblase 4 |
| | N6AMT1 | N-6 adenine-specific DNA methyltransferase 1 |
| | SLC25A36 | solute carrier family 25 member 36 |
| | SCN3A | sodium voltage-gated channel alpha subunit 3 |
| | BICDL1 | BICD family like cargo adaptor 1 |
| | SLC25A32 | solute carrier family 25 member 32 |
| | SPRY2 | sprouty RTK signaling antagonist 2 |
| | TBL1XR1 | transducin beta like 1 X-linked receptor 1 |
| | PPA2 | pyrophosphatase (inorganic) 2 |
| | EMC2 | ER membrane protein complex subunit 2 |
| | DSC2 | desmocollin 2 |
| | PITX2 | paired like homeodomain 2 |
| | ZNF728 | zinc finger protein 728 |
| | PCBP2 | poly(rC) binding protein 2 |
| | AMER2 | APC membrane recruitment protein 2 |
| | CABCOCO1 | ciliary associated calcium binding coiled-coil 1 |
| | LIN28A | lin-28 homolog A |
| | ARHGAP24 | Rho GTPase activating protein 24 |
| | CADM1 | cell adhesion molecule 1 |
| | ZBTB43 | zinc finger and BTB domain containing 43 |
| | THAP3 | THAP domain containing 3 |
| | ARHGAP29 | Rho GTPase activating protein 29 |
| | FAM122A | family with sequence similarity 122A |
| | SLIT2 | slit guidance ligand 2 |
| | FUBP3 | far upstream element binding protein 3 |
| | UBN2 | ubinuclein 2 |
| | SLC39A10 | solute carrier family 39 member 10 |
| | RUSC1 | RUN and SH3 domain containing 1 |
| | CDH19 | cadherin 19 |
| | MAPK3 | mitogen-activated protein kinase 3 |
| | LRG1 | leucine rich alpha-2-glycoprotein 1 |
| | PIK3R1 | phosphoinositide-3-kinase regulatory subunit 1 |
| | ZBTB10 | zinc finger and BTB domain containing 10 |
| | KIF3C | kinesin family member 3C |
| | RETREG1 | reticulophagy regulator 1 |
| | SP1 | Sp1 transcription factor |
| | UFM1 | ubiquitin fold modifier 1 |

| | | |
|---|---|---|
| | PKHD1L1 | PKHD1 like 1 |
| | MTHFS | methenyltetrahydrofolate synthetase |
| | PRDM1 | PR/SET domain 1 |
| | INO80D | INO80 complex subunit D |
| | SOCS4 | suppressor of cytokine signaling 4 |
| | TCF21 | transcription factor 21 |
| | SLC25A37 | solute carrier family 25 member 37 |
| | SMIM11A | small integral membrane protein 11A |
| | SMIM11B | small integral membrane protein 11B |
| | VPS37A | VPS37A, ESCRT-I subunit |
| | HS3ST3A1 | heparan sulfate-glucosamine 3-sulfotransferase 3A1 |
| | TPBG | trophoblast glycoprotein |
| | SH3YL1 | SH3 and SYLF domain containing 1 |
| | ITGAV | integrin subunit alpha V |
| | CMBL | carboxymethylenebutenolidase homolog |
| | ADIPOR2 | adiponectin receptor 2 |
| | ZNF99 | zinc finger protein 99 |
| | PCYT1B | phosphate cytidylyltransferase 1, choline, beta |
| | RFC2 | replication factor C subunit 2 |
| | RGS21 | regulator of G protein signaling 21 |
| | HRK | harakiri, BCL2 interacting protein |
| | ETV1 | ETS variant 1 |
| | PCDH7 | protocadherin 7 |
| | LMNB1 | lamin B1 |
| | MTCH2 | mitochondrial carrier 2 |
| | FAM227A | family with sequence similarity 227 member A |
| | SLC7A11 | solute carrier family 7 member 11 |
| | CRB3 | crumbs cell polarity complex component 3 |
| | ZNF483 | zinc finger protein 483 |
| | NPHS1 | NPHS1, nephrin |
| | HOXD10 | homeobox D10 |
| | APLP2 | amyloid beta precursor like protein 2 |
| | RTKN2 | rhotekin 2 |
| | ELK4 | ELK4, ETS transcription factor |
| | LAMA4 | laminin subunit alpha 4 |
| | ID2 | inhibitor of DNA binding 2 |
| | LRP8 | LDL receptor related protein 8 |
| | ADGRB3 | adhesion G protein-coupled receptor B3 |
| | TEAD1 | TEA domain transcription factor 1 |
| | FREM1 | FRAS1 related extracellular matrix 1 |
| | C14orf177 | chromosome 14 open reading frame 177 |
| | UCK2 | uridine-cytidine kinase 2 |

| | | |
|---|---|---|
| | CNTNAP1 | contactin associated protein 1 |
| | TMEM263 | transmembrane protein 263 |
| | NAALADL2 | N-acetylated alpha-linked acidic dipeptidase like 2 |
| | PLSCR1 | phospholipid scramblase 1 |
| | HOOK3 | hook microtubule tethering protein 3 |
| | LMO1 | LIM domain only 1 |
| | BBS7 | Bardet-Biedl syndrome 7 |
| | ING3 | inhibitor of growth family member 3 |
| | STK4 | serine/threonine kinase 4 |
| | NAA50 | N(alpha)-acetyltransferase 50, NatE catalytic subunit |
| | PLEKHG7 | pleckstrin homology and RhoGEF domain containing G7 |
| | MYO3B | myosin IIIB |
| | PNISR | PNN interacting serine and arginine rich protein |
| | LRRFIP1 | LRR binding FLII interacting protein 1 |
| | MDGA2 | MAM domain containing glycosylphosphatidylinositol anchor 2 |
| | GABRB2 | gamma-aminobutyric acid type A receptor beta2 subunit |
| | MNT | MAX network transcriptional repressor |
| | KIF16B | kinesin family member 16B |
| | GALNT6 | polypeptide N-acetylgalactosaminyltransferase 6 |
| | RORA | RAR related orphan receptor A |
| | CAT | catalase |
| | WWTR1 | WW domain containing transcription regulator 1 |
| | PEX3 | peroxisomal biogenesis factor 3 |
| | GABRA1 | gamma-aminobutyric acid type A receptor alpha1 subunit |
| | TRAF6 | TNF receptor associated factor 6 |
| | XIAP | X-linked inhibitor of apoptosis |
| | PLGLB1 | plasminogen-like B1 |
| | PLGLB2 | plasminogen-like B2 |
| | LUM | lumican |
| | CCSER1 | coiled-coil serine rich protein 1 |
| | PSMD5 | proteasome 26S subunit, non-ATPase 5 |
| | PIGM | phosphatidylinositol glycan anchor biosynthesis class M |
| | SHANK2 | SH3 and multiple ankyrin repeat domains 2 |
| | CRY1 | cryptochrome circadian regulator 1 |
| | ADH7 | alcohol dehydrogenase 7 (class IV), mu or sigma polypeptide |
| | TFCP2 | transcription factor CP2 |
| | SETD7 | SET domain containing 7, histone lysine methyltransferase |
| | AKAP8 | A-kinase anchoring protein 8 |
| | AIRE | autoimmune regulator |
| | ZNF704 | zinc finger protein 704 |
| | ZNF570 | zinc finger protein 570 |
| | B3GLCT | beta 3-glucosyltransferase |

| | | |
|---|---|---|
| | EIF2S1 | eukaryotic translation initiation factor 2 subunit alpha |
| | OPRM1 | opioid receptor mu 1 |
| | SYP | synaptophysin |
| | APOL6 | apolipoprotein L6 |
| | QRICH1 | glutamine rich 1 |
| | TMEM100 | transmembrane protein 100 |
| | LDLR | low density lipoprotein receptor |
| | TRIP4 | thyroid hormone receptor interactor 4 |
| | SIK3 | SIK family kinase 3 |
| | CNKSR2 | connector enhancer of kinase suppressor of Ras 2 |
| | CDKN2B | cyclin dependent kinase inhibitor 2B |
| | FBXO36 | F-box protein 36 |
| | TRAM1 | translocation associated membrane protein 1 |
| | PTAR1 | protein prenyltransferase alpha subunit repeat containing 1 |
| | ZNF451 | zinc finger protein 451 |
| | FCHO2 | FCH domain only 2 |
| | IBTK | inhibitor of Bruton tyrosine kinase |
| | CASC3 | CASC3, exon junction complex subunit |
| | LAMP1 | lysosomal associated membrane protein 1 |
| | PIK3AP1 | phosphoinositide-3-kinase adaptor protein 1 |
| | ATXN1 | ataxin 1 |
| | GIGYF2 | GRB10 interacting GYF protein 2 |
| | MAGEL2 | MAGE family member L2 |
| | SHROOM1 | shroom family member 1 |
| | SUSD5 | sushi domain containing 5 |
| | DNMT3A | DNA methyltransferase 3 alpha |
| | OXCT1 | 3-oxoacid CoA-transferase 1 |
| | SC5D | sterol-C5-desaturase |
| | TP63 | tumor protein p63 |
| | RBAK | RB associated KRAB zinc finger |
| | CWC15 | CWC15 spliceosome associated protein homolog |
| | C2CD2 | C2 calcium dependent domain containing 2 |
| | RRAS2 | RAS related 2 |
| | ZSWIM8 | zinc finger SWIM-type containing 8 |
| | ZDHHC3 | zinc finger DHHC-type containing 3 |
| | GOLGA1 | golgin A1 |
| | NIPBL | NIPBL, cohesin loading factor |
| | CEP44 | centrosomal protein 44 |
| | CCDC62 | coiled-coil domain containing 62 |
| | ZNF518A | zinc finger protein 518A |
| | WDTC1 | WD and tetratricopeptide repeats 1 |
| | POMP | proteasome maturation protein |

| | | |
|---|---|---|
| | SQLE | squalene epoxidase |
| | SEM1 | SEM1, 26S proteasome complex subunit |
| | ATRX | ATRX, chromatin remodeler |
| | HELZ | helicase with zinc finger |
| | KLHDC1 | kelch domain containing 1 |
| | RNF217 | ring finger protein 217 |
| | TRUB1 | TruB pseudouridine synthase family member 1 |
| | CTLA4 | cytotoxic T-lymphocyte associated protein 4 |
| | DEGS1 | delta 4-desaturase, sphingolipid 1 |
| | FBXO38 | F-box protein 38 |
| | ATP2C1 | ATPase secretory pathway Ca2+ transporting 1 |
| | XKR6 | XK related 6 |
| | ZNF100 | zinc finger protein 100 |
| | HDAC4 | histone deacetylase 4 |
| | ZNF713 | zinc finger protein 713 |
| | LARP4 | La ribonucleoprotein domain family member 4 |
| | MEIOC | meiosis specific with coiled-coil domain |
| | ELMSAN1 | ELM2 and Myb/SANT domain containing 1 |
| | CACYBP | calcyclin binding protein |
| | VAMP4 | vesicle associated membrane protein 4 |
| | KLC4 | kinesin light chain 4 |
| | ANGPT1 | angiopoietin 1 |
| | C20orf194 | chromosome 20 open reading frame 194 |
| | MAL2 | mal, T cell differentiation protein 2 (gene/pseudogene) |
| | ENY2 | ENY2, transcription and export complex 2 subunit |
| | SLC9A6 | solute carrier family 9 member A6 |
| | MTMR7 | myotubularin related protein 7 |
| | GJC1 | gap junction protein gamma 1 |
| | STRN3 | striatin 3 |
| | IRAK4 | interleukin 1 receptor associated kinase 4 |
| | C17orf77 | chromosome 17 open reading frame 77 |
| | TTC7A | tetratricopeptide repeat domain 7A |
| | FOPNL | FGFR1OP N-terminal like |
| | LILRB4 | leukocyte immunoglobulin like receptor B4 |
| | NPC1 | NPC intracellular cholesterol transporter 1 |
| | FAM241A | family with sequence similarity 241 member A |
| | EVI2B | ecotropic viral integration site 2B |
| | POLK | DNA polymerase kappa |
| | MSI2 | musashi RNA binding protein 2 |
| | KMT2E | lysine methyltransferase 2E |
| | PGAP1 | post-GPI attachment to proteins 1 |
| | FOXD3 | forkhead box D3 |

| | | |
|---|---|---|
| | TMEM165 | transmembrane protein 165 |
| | ZNF676 | zinc finger protein 676 |
| | GUCY1B1 | guanylate cyclase 1 soluble subunit beta 1 |
| | COL21A1 | collagen type XXI alpha 1 chain |
| | ATXN3 | ataxin 3 |
| | ACAP2 | ArfGAP with coiled-coil, ankyrin repeat and PH domains 2 |
| | BBX | BBX, HMG-box containing |
| | MON2 | MON2 homolog, regulator of endosome-to-Golgi trafficking |
| | TMX4 | thioredoxin related transmembrane protein 4 |
| | SNAP29 | synaptosome associated protein 29 |
| | RLIM | ring finger protein, LIM domain interacting |
| | PRPF18 | pre-mRNA processing factor 18 |
| | ATXN7L1 | ataxin 7 like 1 |
| | RBM12 | RNA binding motif protein 12 |
| | PATE1 | prostate and testis expressed 1 |
| | MARCHF2 | membrane associated ring-CH-type finger 2 |
| | SETD5 | SET domain containing 5 |
| | KLF4 | Kruppel like factor 4 |
| | PDS5A | PDS5 cohesin associated factor A |
| | EYS | eyes shut homolog |
| | CCR8 | C-C motif chemokine receptor 8 |
| | ERCC4 | ERCC excision repair 4, endonuclease catalytic subunit |
| | CENPP | centromere protein P |
| | TXN | thioredoxin |
| | DCAF16 | DDB1 and CUL4 associated factor 16 |
| | DHX40 | DEAH-box helicase 40 |
| | LIN28B | lin-28 homolog B |
| | PLEKHA2 | pleckstrin homology domain containing A2 |
| | TMEM108 | transmembrane protein 108 |
| | SVEP1 | sushi, von Willebrand factor type A, EGF and pentraxin domain containing 1 |
| | TPR | translocated promoter region, nuclear basket protein |
| | GLTP | glycolipid transfer protein |
| | FAM92A | family with sequence similarity 92 member A |
| | KIRREL1 | kirre like nephrin family adhesion molecule 1 |
| | IDH1 | isocitrate dehydrogenase (NADP(+)) 1, cytosolic |
| | RANBP10 | RAN binding protein 10 |
| | TCF3 | transcription factor 3 |
| | PLAGL2 | PLAG1 like zinc finger 2 |
| | HEATR5A | HEAT repeat containing 5A |
| | IRAK1BP1 | interleukin 1 receptor associated kinase 1 binding protein 1 |
| | FGD4 | FYVE, RhoGEF and PH domain containing 4 |

| | | |
|---|---|---|
| | TENT2 | terminal nucleotidyltransferase 2 |
| | OGFRL1 | opioid growth factor receptor like 1 |
| | FAT4 | FAT atypical cadherin 4 |
| | ZNF236 | zinc finger protein 236 |
| | RALGDS | ral guanine nucleotide dissociation stimulator |
| | SYNGAP1 | synaptic Ras GTPase activating protein 1 |
| | PCDHA9 | protocadherin alpha 9 |
| | WDR72 | WD repeat domain 72 |
| | POLR2G | RNA polymerase II subunit G |
| | DLG3 | discs large MAGUK scaffold protein 3 |
| | SLC30A10 | solute carrier family 30 member 10 |
| | TSHZ3 | teashirt zinc finger homeobox 3 |
| | CCDC18 | coiled-coil domain containing 18 |
| | ABHD18 | abhydrolase domain containing 18 |
| | PIAS2 | protein inhibitor of activated STAT 2 |
| | DNAJA1 | DnaJ heat shock protein family (Hsp40) member A1 |
| | TMEM218 | transmembrane protein 218 |
| | TRAT1 | T cell receptor associated transmembrane adaptor 1 |
| | CAPRIN1 | cell cycle associated protein 1 |
| | NANP | N-acetylneuraminic acid phosphatase |
| | UBA6 | ubiquitin like modifier activating enzyme 6 |
| | OSBPL8 | oxysterol binding protein like 8 |
| | PCF11 | PCF11, cleavage and polyadenylation factor subunit |
| | GPR171 | G protein-coupled receptor 171 |
| | SMAD6 | SMAD family member 6 |
| | CERS6 | ceramide synthase 6 |
| | COBLL1 | cordon-bleu WH2 repeat protein like 1 |
| | JAM3 | junctional adhesion molecule 3 |
| | SVIL | supervillin |
| | GDAP2 | ganglioside induced differentiation associated protein 2 |
| | ZDHHC17 | zinc finger DHHC-type containing 17 |
| | DENND4C | DENN domain containing 4C |
| | WIPF1 | WAS/WASL interacting protein family member 1 |
| | BBS2 | Bardet-Biedl syndrome 2 |
| | MEDAG | mesenteric estrogen dependent adipogenesis |
| | REV3L | REV3 like, DNA directed polymerase zeta catalytic subunit |
| | HVCN1 | hydrogen voltage gated channel 1 |
| | COMT | catechol-O-methyltransferase |
| | ME1 | malic enzyme 1 |
| | NPAS3 | neuronal PAS domain protein 3 |
| | IAH1 | isoamyl acetate hydrolyzing esterase 1 (putative) |
| | SET | SET nuclear proto-oncogene |

| | | |
|---|---|---|
| | IL7 | interleukin 7 |
| | VPS13C | vacuolar protein sorting 13 homolog C |
| | MAGT1 | magnesium transporter 1 |
| | ANKRD13A | ankyrin repeat domain 13A |
| | YTHDF3 | YTH N6-methyladenosine RNA binding protein 3 |
| | ZFP36L1 | ZFP36 ring finger protein like 1 |
| | HHIP | hedgehog interacting protein |
| | NKAIN2 | sodium/potassium transporting ATPase interacting 2 |
| | ETNPPL | ethanolamine-phosphate phospho-lyase |
| | RAB18 | RAB18, member RAS oncogene family |
| | KLHL28 | kelch like family member 28 |
| | SENP7 | SUMO specific peptidase 7 |
| | MDM4 | MDM4, p53 regulator |
| | MTPN | myotrophin |
| | LRRTM2 | leucine rich repeat transmembrane neuronal 2 |
| | MXD1 | MAX dimerization protein 1 |
| | NUP58 | nucleoporin 58 |
| | CTHRC1 | collagen triple helix repeat containing 1 |
| | PET117 | PET117 homolog |
| | ANKS1A | ankyrin repeat and sterile alpha motif domain containing 1A |
| | SOWAHC | sosondowah ankyrin repeat domain family member C |
| | FLT1 | fms related tyrosine kinase 1 |
| | PPM1E | protein phosphatase, Mg2+/Mn2+ dependent 1E |
| | ZDHHC2 | zinc finger DHHC-type containing 2 |
| | NR2C2 | nuclear receptor subfamily 2 group C member 2 |
| | TMEM248 | transmembrane protein 248 |
| | ZEB1 | zinc finger E-box binding homeobox 1 |
| | IKZF5 | IKAROS family zinc finger 5 |
| | UNC5C | unc-5 netrin receptor C |
| | SYT2 | synaptotagmin 2 |
| | MTR | 5-methyltetrahydrofolate-homocysteine methyltransferase |
| | MAP3K12 | mitogen-activated protein kinase kinase kinase 12 |
| | KDM2A | lysine demethylase 2A |
| | ZNF706 | zinc finger protein 706 |
| | TRAPPC8 | trafficking protein particle complex 8 |
| | MBD2 | methyl-CpG binding domain protein 2 |
| | PRKACB | protein kinase cAMP-activated catalytic subunit beta |
| | SEC62 | SEC62 homolog, preprotein translocation factor |
| | BRF2 | BRF2, RNA polymerase III transcription initiation factor subunit |
| | KIF3A | kinesin family member 3A |
| | TBC1D9 | TBC1 domain family member 9 |

| | | |
|---|---|---|
| | MLX | MLX, MAX dimerization protein |
| | CACNB1 | calcium voltage-gated channel auxiliary subunit beta 1 |
| | RAB11A | RAB11A, member RAS oncogene family |
| | C1GALT1 | core 1 synthase, glycoprotein-N-acetylgalactosamine 3-beta-galactosyltransferase 1 |
| | CHMP4C | charged multivesicular body protein 4C |
| | ANGPTL2 | angiopoietin like 2 |
| | MRPS25 | mitochondrial ribosomal protein S25 |
| | LAMA3 | laminin subunit alpha 3 |
| | FEM1C | fem-1 homolog C |
| | ETS1 | ETS proto-oncogene 1, transcription factor |
| | USP32 | ubiquitin specific peptidase 32 |
| | ABL2 | ABL proto-oncogene 2, non-receptor tyrosine kinase |
| | BRCC3 | BRCA1/BRCA2-containing complex subunit 3 |
| | MIXL1 | Mix paired-like homeobox |
| | TECRL | trans-2,3-enoyl-CoA reductase like |
| | LDLRAD3 | low density lipoprotein receptor class A domain containing 3 |
| | ZNF182 | zinc finger protein 182 |
| | ARHGEF35 | Rho guanine nucleotide exchange factor 35 |
| | ZMYM5 | zinc finger MYM-type containing 5 |
| | KHK | ketohexokinase |
| | ACTR2 | ARP2 actin related protein 2 homolog |
| | NDFIP2 | Nedd4 family interacting protein 2 |
| | RBM17 | RNA binding motif protein 17 |
| | NIPSNAP3B | nipsnap homolog 3B |
| | PDE10A | phosphodiesterase 10A |
| | CCDC173 | coiled-coil domain containing 173 |
| | GINS1 | GINS complex subunit 1 |
| | NDUFA2 | NADH:ubiquinone oxidoreductase subunit A2 |
| | FOXN2 | forkhead box N2 |
| | CYP20A1 | cytochrome P450 family 20 subfamily A member 1 |
| | PIFO | primary cilia formation |
| | PREPL | prolyl endopeptidase like |
| | ZNF280C | zinc finger protein 280C |
| | RHAG | Rh associated glycoprotein |
| | SLC25A30 | solute carrier family 25 member 30 |
| | BCL11A | BCL11A, BAF complex component |
| | CLDN22 | claudin 22 |
| | JKAMP | JNK1/MAPK8 associated membrane protein |
| | ADAM28 | ADAM metallopeptidase domain 28 |
| | CLIP4 | CAP-Gly domain containing linker protein family member 4 |
| | DZIP1 | DAZ interacting zinc finger protein 1 |

| | | |
|---|---|---|
| | RB1 | RB transcriptional corepressor 1 |
| | ZBTB20 | zinc finger and BTB domain containing 20 |
| | VAT1L | vesicle amine transport 1 like |
| | CACNA1B | calcium voltage-gated channel subunit alpha1 B |
| | FAM110C | family with sequence similarity 110 member C |
| | G2E3 | G2/M-phase specific E3 ubiquitin protein ligase |
| | ATF1 | activating transcription factor 1 |
| | COL25A1 | collagen type XXV alpha 1 chain |
| | RWDD1 | RWD domain containing 1 |
| | CHM | CHM, Rab escort protein 1 |
| | FBLN2 | fibulin 2 |
| | CSNK1G3 | casein kinase 1 gamma 3 |
| | KHSRP | KH-type splicing regulatory protein |
| | GPATCH8 | G-patch domain containing 8 |
| | CBLB | Cbl proto-oncogene B |
| | CHML | CHM like, Rab escort protein 2 |
| | SUN1 | Sad1 and UNC84 domain containing 1 |
| | USP45 | ubiquitin specific peptidase 45 |
| | SELENOT | selenoprotein T |
| | TFPI2 | tissue factor pathway inhibitor 2 |
| | ESRRG | estrogen related receptor gamma |
| | GPR18 | G protein-coupled receptor 18 |
| | TCIM | transcriptional and immune response regulator |
| | ATP2B2 | ATPase plasma membrane Ca2+ transporting 2 |
| | PDZRN3 | PDZ domain containing ring finger 3 |
| | TSNAX | translin associated factor X |
| | EPS15 | epidermal growth factor receptor pathway substrate 15 |
| | FGF2 | fibroblast growth factor 2 |
| | TBX18 | T-box 18 |
| | HTR2C | 5-hydroxytryptamine receptor 2C |
| | GRID2 | glutamate ionotropic receptor delta type subunit 2 |
| | PLA2G12A | phospholipase A2 group XIIA |
| | USP9Y | ubiquitin specific peptidase 9 Y-linked |
| | PPP4R2 | protein phosphatase 4 regulatory subunit 2 |
| | PPP1R7 | protein phosphatase 1 regulatory subunit 7 |
| | RRP15 | ribosomal RNA processing 15 homolog |
| | HCFC2 | host cell factor C2 |
| | GAS2L3 | growth arrest specific 2 like 3 |
| | NUFIP2 | nuclear FMR1 interacting protein 2 |
| | ANLN | anillin actin binding protein |
| | NEDD4 | neural precursor cell expressed, developmentally down-regulated 4, E3 ubiquitin protein ligase |

| | | |
|---|---|---|
| | LHX8 | LIM homeobox 8 |
| | NR2F2 | nuclear receptor subfamily 2 group F member 2 |
| | SP3 | Sp3 transcription factor |
| | ESYT2 | extended synaptotagmin 2 |
| | TMEM212 | transmembrane protein 212 |
| | SFRP2 | secreted frizzled related protein 2 |
| | TM6SF1 | transmembrane 6 superfamily member 1 |
| | TRAM1L1 | translocation associated membrane protein 1 like 1 |
| | PODN | podocan |
| | TYW3 | tRNA-yW synthesizing protein 3 homolog |
| | SYTL5 | synaptotagmin like 5 |
| | CYP7B1 | cytochrome P450 family 7 subfamily B member 1 |
| | DR1 | down-regulator of transcription 1 |
| | FCER1A | Fc fragment of IgE receptor Ia |
| | NXPE3 | neurexophilin and PC-esterase domain family member 3 |
| | ROBO2 | roundabout guidance receptor 2 |
| | TMEM69 | transmembrane protein 69 |
| | AIG1 | androgen induced 1 |
| | MTX3 | metaxin 3 |
| | CBX4 | chromobox 4 |
| | ZNF681 | zinc finger protein 681 |
| | WDR26 | WD repeat domain 26 |
| | HOXB5 | homeobox B5 |
| | IQCC | IQ motif containing C |
| | ARL13B | ADP ribosylation factor like GTPase 13B |
| | CEBPG | CCAAT enhancer binding protein gamma |
| | RAB33B | RAB33B, member RAS oncogene family |
| | HACE1 | HECT domain and ankyrin repeat containing E3 ubiquitin protein ligase 1 |
| | SNAP25 | synaptosome associated protein 25 |
| | MAPK6 | mitogen-activated protein kinase 6 |
| | AKR1C2 | aldo-keto reductase family 1 member C2 |
| | ITGB8 | integrin subunit beta 8 |
| | INTS6L | integrator complex subunit 6 like |
| | ISM1 | isthmin 1 |
| | SETD9 | SET domain containing 9 |
| | USP27X | ubiquitin specific peptidase 27 X-linked |
| | RSBN1L | round spermatid basic protein 1 like |
| | EMP1 | epithelial membrane protein 1 |
| | PRDM8 | PR/SET domain 8 |
| | TSPYL1 | TSPY like 1 |
| | PBRM1 | polybromo 1 |

| | | |
|---|---|---|
| | RSBN1 | round spermatid basic protein 1 |
| | CCDC186 | coiled-coil domain containing 186 |
| | MFSD9 | major facilitator superfamily domain containing 9 |
| | GCOM1 | GRINL1A complex locus 1 |
| | MAPK1IP1L | mitogen-activated protein kinase 1 interacting protein 1 like |
| | POLR2M | RNA polymerase II subunit M |
| | CAV2 | caveolin 2 |
| | LYST | lysosomal trafficking regulator |
| | THAP1 | THAP domain containing 1 |
| | ADAT2 | adenosine deaminase, tRNA specific 2 |
| | FRMD6 | FERM domain containing 6 |
| | DIPK2A | divergent protein kinase domain 2A |
| | MIER3 | MIER family member 3 |
| | EFCAB13 | EF-hand calcium binding domain 13 |
| | ZDHHC21 | zinc finger DHHC-type containing 21 |
| | UBR5 | ubiquitin protein ligase E3 component n-recognin 5 |
| | SCX | scleraxis bHLH transcription factor |
| | LRRFIP2 | LRR binding FLII interacting protein 2 |
| | SKP2 | S-phase kinase associated protein 2 |
| | PIN4 | peptidylprolyl cis/trans isomerase, NIMA-interacting 4 |
| | COMMD8 | COMM domain containing 8 |
| | OTX1 | orthodenticle homeobox 1 |
| | XPO4 | exportin 4 |
| | AFMID | arylformamidase |
| | ANOS1 | anosmin 1 |
| | MYADM | myeloid associated differentiation marker |
| Segment-2-PB1- MR2 5P | USP42 | ubiquitin specific peptidase 42 |
| | SAA2 | serum amyloid A2 |
| | SYT11 | synaptotagmin 11 |
| | LMNA | lamin A/C |
| | NME7 | NME/NM23 family member 7 |
| | EDN3 | endothelin 3 |
| | MBD6 | methyl-CpG binding domain protein 6 |
| | EYA4 | EYA transcriptional coactivator and phosphatase 4 |
| | ITPKA | inositol-trisphosphate 3-kinase A |
| | CASTOR3 | CASTOR family member 3 |
| | CHRNA6 | cholinergic receptor nicotinic alpha 6 subunit |
| | RSPO3 | R-spondin 3 |
| | ZMYM2 | zinc finger MYM-type containing 2 |
| | CD53 | CD53 molecule |
| | CD9 | CD9 molecule |

| | | |
|---|---|---|
| | SOCS2 | suppressor of cytokine signaling 2 |
| | ZFYVE9 | zinc finger FYVE-type containing 9 |
| | BCL6 | BCL6, transcription repressor |
| | KIAA2012 | KIAA2012 |
| | PIK3C2A | phosphatidylinositol-4-phosphate 3-kinase catalytic subunit type 2 alpha |
| | FBXO11 | F-box protein 11 |
| | CCDC178 | coiled-coil domain containing 178 |
| | KDELR1 | KDEL endoplasmic reticulum protein retention receptor 1 |
| | MMP16 | matrix metallopeptidase 16 |
| | COL19A1 | collagen type XIX alpha 1 chain |
| | ATP5F1C | ATP synthase F1 subunit gamma |
| | SETD5 | SET domain containing 5 |
| | MGST2 | microsomal glutathione S-transferase 2 |
| | CEP19 | centrosomal protein 19 |
| | SOX6 | SRY-box 6 |
| | GFPT2 | glutamine-fructose-6-phosphate transaminase 2 |
| | RAB39A | RAB39A, member RAS oncogene family |
| | KHDRBS1 | KH RNA binding domain containing, signal transduction associated 1 |
| | TTC39C | tetratricopeptide repeat domain 39C |
| | ZFX | zinc finger protein X-linked |
| | SH3BGRL | SH3 domain binding glutamate rich protein like |
| | LRRC17 | leucine rich repeat containing 17 |
| | NAA50 | N(alpha)-acetyltransferase 50, NatE catalytic subunit |
| | VKORC1L1 | vitamin K epoxide reductase complex subunit 1 like 1 |
| | CDC23 | cell division cycle 23 |
| | GPR161 | G protein-coupled receptor 161 |
| | PLA2R1 | phospholipase A2 receptor 1 |
| | ARHGAP26 | Rho GTPase activating protein 26 |
| | RAB9B | RAB9B, member RAS oncogene family |
| | SEC24A | SEC24 homolog A, COPII coat complex component |
| | SASH1 | SAM and SH3 domain containing 1 |
| | CLEC2D | C-type lectin domain family 2 member D |
| | THEG5 | testis highly expressed protein 5 |
| | USP35 | ubiquitin specific peptidase 35 |
| | SOGA3 | SOGA family member 3 |
| | RGS5 | regulator of G protein signaling 5 |
| | ANTXR2 | ANTXR cell adhesion molecule 2 |
| | KLHL6 | kelch like family member 6 |
| | SLC35E1 | solute carrier family 35 member E1 |
| | UBE3D | ubiquitin protein ligase E3D |
| | POGZ | pogo transposable element derived with ZNF domain |

| | | |
|---|---|---|
| | NLK | nemo like kinase |
| | NUDT15 | nudix hydrolase 15 |
| | PRDM2 | PR/SET domain 2 |
| | VMA21 | VMA21, vacuolar ATPase assembly factor |
| | FAM217B | family with sequence similarity 217 member B |
| Segment-2-PB1- MR2 3P | TBP | TATA-box binding protein |
| | WDR11 | WD repeat domain 11 |
| | MYT1L | myelin transcription factor 1 like |
| | SLIT2 | slit guidance ligand 2 |
| | TUBGCP5 | tubulin gamma complex associated protein 5 |
| | ZNF90 | zinc finger protein 90 |
| | SRBD1 | S1 RNA binding domain 1 |
| | WNK3 | WNK lysine deficient protein kinase 3 |
| | COQ8A | coenzyme Q8A |
| | CPSF7 | cleavage and polyadenylation specific factor 7 |
| | GSTA3 | glutathione S-transferase alpha 3 |
| | FOXN3 | forkhead box N3 |
| | EPHA7 | EPH receptor A7 |
| | RABGAP1L | RAB GTPase activating protein 1 like |
| | MGA | MGA, MAX dimerization protein |
| | SPATA4 | spermatogenesis associated 4 |
| | JPH1 | junctophilin 1 |
| | S100A3 | S100 calcium binding protein A3 |
| | NUS1 | NUS1, dehydrodolichyl diphosphate synthase subunit |
| | NEUROD4 | neuronal differentiation 4 |
| | KDM2A | lysine demethylase 2A |
| | KRTAP2-2 | keratin associated protein 2-2 |
| | MYT1 | myelin transcription factor 1 |
| | ZNF577 | zinc finger protein 577 |
| | RPL37 | ribosomal protein L37 |
| | PCNX1 | pecanex 1 |
| | EEF2K | eukaryotic elongation factor 2 kinase |
| | UBP1 | upstream binding protein 1 |
| | KCND1 | potassium voltage-gated channel subfamily D member 1 |
| | TXNDC17 | thioredoxin domain containing 17 |
| | PHLPP1 | PH domain and leucine rich repeat protein phosphatase 1 |
| | MAGOHB | mago homolog B, exon junction complex subunit |
| | WTAP | WT1 associated protein |
| | SLC44A1 | solute carrier family 44 member 1 |
| | ASB7 | ankyrin repeat and SOCS box containing 7 |
| | NDUFB5 | NADH:ubiquinone oxidoreductase subunit B5 |
| | ZNF562 | zinc finger protein 562 |

| | | |
|---|---|---|
| | ZSWIM8 | zinc finger SWIM-type containing 8 |
| | MBTD1 | mbt domain containing 1 |
| | PPIB | peptidylprolyl isomerase B |
| | CWC15 | CWC15 spliceosome associated protein homolog |
| | ITFG1 | integrin alpha FG-GAP repeat containing 1 |
| | RD3L | retinal degeneration 3 like |
| | GSTA1 | glutathione S-transferase alpha 1 |
| | GSTA2 | glutathione S-transferase alpha 2 |
| | SNTG1 | syntrophin gamma 1 |
| | PPTC7 | PTC7 protein phosphatase homolog |
| | RWDD2A | RWD domain containing 2A |
| | ARL6IP1 | ADP ribosylation factor like GTPase 6 interacting protein 1 |
| | ACBD5 | acyl-CoA binding domain containing 5 |
| | SPATA31D3 | SPATA31 subfamily D member 3 |
| | TENM1 | teneurin transmembrane protein 1 |
| | GFOD2 | glucose-fructose oxidoreductase domain containing 2 |
| Segment-3-PA- MD8 5P | CLN8 | CLN8, transmembrane ER and ERGIC protein |
| | STON2 | stonin 2 |
| | ADGRL3 | adhesion G protein-coupled receptor L3 |
| | ACLY | ATP citrate lyase |
| | CNTN4 | contactin 4 |
| | BIRC2 | baculoviral IAP repeat containing 2 |
| | LEPR | leptin receptor |
| | DOK6 | docking protein 6 |
| | RBM23 | RNA binding motif protein 23 |
| | OXR1 | oxidation resistance 1 |
| | GCFC2 | GC-rich sequence DNA-binding factor 2 |
| | FSHR | follicle stimulating hormone receptor |
| | KIAA1549 | KIAA1549 |
| | TDO2 | tryptophan 2,3-dioxygenase |
| | CEPT1 | choline/ethanolamine phosphotransferase 1 |
| | SSX2IP | SSX family member 2 interacting protein |
| | RCAN3 | RCAN family member 3 |
| | RAB5A | RAB5A, member RAS oncogene family |
| | EIF2A | eukaryotic translation initiation factor 2A |
| | PDE3A | phosphodiesterase 3A |
| | MAP4K4 | mitogen-activated protein kinase kinase kinase kinase 4 |
| | INSYN2 | inhibitory synaptic factor 2A |
| | HOMEZ | homeobox and leucine zipper encoding |
| | FAM20A | FAM20A, golgi associated secretory pathway pseudokinase |
| | CUL3 | cullin 3 |
| | HCFC2 | host cell factor C2 |

| | | |
|---|---|---|
| | MMP16 | matrix metallopeptidase 16 |
| | GPATCH2L | G-patch domain containing 2 like |
| | DMGDH | dimethylglycine dehydrogenase |
| | RYR3 | ryanodine receptor 3 |
| | GRID2 | glutamate ionotropic receptor delta type subunit 2 |
| | INTS2 | integrator complex subunit 2 |
| | KCNJ8 | potassium voltage-gated channel subfamily J member 8 |
| | INA | internexin neuronal intermediate filament protein alpha |
| | SNX2 | sorting nexin 2 |
| | CD80 | CD80 molecule |
| | NEDD1 | neural precursor cell expressed, developmentally down-regulated 1 |
| | IRF4 | interferon regulatory factor 4 |
| | NAA30 | N(alpha)-acetyltransferase 30, NatC catalytic subunit |
| | BTBD1 | BTB domain containing 1 |
| | STXBP5L | syntaxin binding protein 5 like |
| | DBN1 | drebrin 1 |
| | H2AFZ | H2A histone family member Z |
| | RBM28 | RNA binding motif protein 28 |
| | PRDM6 | PR/SET domain 6 |
| | RASGRP1 | RAS guanyl releasing protein 1 |
| | MARS2 | methionyl-tRNA synthetase 2, mitochondrial |
| | P2RY13 | purinergic receptor P2Y13 |
| | SP4 | Sp4 transcription factor |
| | ELOVL4 | ELOVL fatty acid elongase 4 |
| | ZNF718 | zinc finger protein 718 |
| | KCNMB3 | potassium calcium-activated channel subfamily M regulatory beta subunit 3 |
| | TSPO2 | translocator protein 2 |
| | SMG1 | SMG1, nonsense mediated mRNA decay associated PI3K related kinase |
| | ZMAT3 | zinc finger matrin-type 3 |
| | DNAH11 | dynein axonemal heavy chain 11 |
| | MRPS14 | mitochondrial ribosomal protein S14 |
| | SPRTN | SprT-like N-terminal domain |
| | MTA2 | metastasis associated 1 family member 2 |
| | FGF7 | fibroblast growth factor 7 |
| | ARF6 | ADP ribosylation factor 6 |
| | TMEM30B | transmembrane protein 30B |
| | C5orf49 | chromosome 5 open reading frame 49 |
| | SLC35C2 | solute carrier family 35 member C2 |
| | RSF1 | remodeling and spacing factor 1 |
| | DBX2 | developing brain homeobox 2 |

| | | |
|---|---|---|
| | ARMCX3 | armadillo repeat containing X-linked 3 |
| | C5orf22 | chromosome 5 open reading frame 22 |
| | LSM14A | LSM14A, mRNA processing body assembly factor |
| | ERAP1 | endoplasmic reticulum aminopeptidase 1 |
| | NUCKS1 | nuclear casein kinase and cyclin dependent kinase substrate 1 |
| | FUT9 | fucosyltransferase 9 |
| | FAM57A | family with sequence similarity 57 member A |
| | RSL1D1 | ribosomal L1 domain containing 1 |
| | GFM2 | G elongation factor mitochondrial 2 |
| | RAB8A | RAB8A, member RAS oncogene family |
| | SLC7A14 | solute carrier family 7 member 14 |
| | ZCRB1 | zinc finger CCHC-type and RNA binding motif containing 1 |
| | MTHFD2L | methylenetetrahydrofolate dehydrogenase (NADP+ dependent) 2 like |
| | UBE3A | ubiquitin protein ligase E3A |
| | MEAF6 | MYST/Esa1 associated factor 6 |
| | HOOK1 | hook microtubule tethering protein 1 |
| | MPC2 | mitochondrial pyruvate carrier 2 |
| | DNMT3A | DNA methyltransferase 3 alpha |
| Segment-3-PA- MD8 3P | ABCC5 | ATP binding cassette subfamily C member 5 |
| | SMIM17 | small integral membrane protein 17 |
| | KLRF1 | killer cell lectin like receptor F1 |
| | FAM20A | FAM20A, golgi associated secretory pathway pseudokinase |
| | HYDIN | HYDIN, axonemal central pair apparatus protein |
| | GDAP2 | ganglioside induced differentiation associated protein 2 |
| | TMEM41A | transmembrane protein 41A |
| | GASK1A | golgi associated kinase 1A |
| | SAMD4B | sterile alpha motif domain containing 4B |
| | TRAF6 | TNF receptor associated factor 6 |
| | PRMT3 | protein arginine methyltransferase 3 |
| | SLC17A5 | solute carrier family 17 member 5 |
| | SH3GL3 | SH3 domain containing GRB2 like 3, endophilin A3 |
| | SLC41A2 | solute carrier family 41 member 2 |
| | C5orf63 | chromosome 5 open reading frame 63 |
| | FRMD3 | FERM domain containing 3 |
| | CHM | CHM, Rab escort protein 1 |
| | ADAMDEC1 | ADAM like decysin 1 |
| | DOCK9 | dedicator of cytokinesis 9 |
| | RBM24 | RNA binding motif protein 24 |
| | SOS2 | SOS Ras/Rho guanine nucleotide exchange factor 2 |
| | C2orf68 | chromosome 2 open reading frame 68 |
| | APBB3 | amyloid beta precursor protein binding family B member 3 |

| | | |
|---|---|---|
| Segment-3-PA- MR23 5P | EMC7 | ER membrane protein complex subunit 7 |
| | SASS6 | SAS-6 centriolar assembly protein |
| | ZNF766 | zinc finger protein 766 |
| | CACHD1 | cache domain containing 1 |
| | PRSS12 | serine protease 12 |
| | FRMD7 | FERM domain containing 7 |
| | ZSCAN32 | zinc finger and SCAN domain containing 32 |
| | RCBTB1 | RCC1 and BTB domain containing protein 1 |
| | HCFC2 | host cell factor C2 |
| | GPNMB | glycoprotein nmb |
| | WDFY3 | WD repeat and FYVE domain containing 3 |
| | AKAP6 | A-kinase anchoring protein 6 |
| | RAP1A | RAP1A, member of RAS oncogene family |
| | DAPK1 | death associated protein kinase 1 |
| | SPCS2 | signal peptidase complex subunit 2 |
| | EIF5A2 | eukaryotic translation initiation factor 5A2 |
| | SATL1 | spermidine/spermine N1-acetyl transferase like 1 |
| | TRIM9 | tripartite motif containing 9 |
| | CDC14A | cell division cycle 14A |
| | PPTC7 | PTC7 protein phosphatase homolog |
| | CALCRL | calcitonin receptor like receptor |
| | PI4K2B | phosphatidylinositol 4-kinase type 2 beta |
| | ARHGAP5 | Rho GTPase activating protein 5 |
| | FOXP1 | forkhead box P1 |
| | ABCA9 | ATP binding cassette subfamily A member 9 |
| | PTPN13 | protein tyrosine phosphatase, non-receptor type 13 |
| | C3orf70 | chromosome 3 open reading frame 70 |
| | PMCH | pro-melanin concentrating hormone |
| | SMURF2 | SMAD specific E3 ubiquitin protein ligase 2 |
| | CCDC146 | coiled-coil domain containing 146 |
| | SP1 | Sp1 transcription factor |
| | TRAM1 | translocation associated membrane protein 1 |
| | MEX3C | mex-3 RNA binding family member C |
| | SPRY4 | sprouty RTK signaling antagonist 4 |
| | LRP2 | LDL receptor related protein 2 |
| | EML1 | EMAP like 1 |
| | PRDM4 | PR/SET domain 4 |
| | COG2 | component of oligomeric golgi complex 2 |
| | ELK4 | ELK4, ETS transcription factor |
| | F3 | coagulation factor III, tissue factor |
| | FBXL3 | F-box and leucine rich repeat protein 3 |
| | ZNF546 | zinc finger protein 546 |

| | | |
|---|---|---|
| | DIS3 | DIS3 homolog, exosome endoribonuclease and 3'-5' exoribonuclease |
| Segment-3-PA- MR23 3P | ZBTB25 | zinc finger and BTB domain containing 25 |
| | KRAS | KRAS proto-oncogene, GTPase |
| | GLS | glutaminase |
| | EIF5A2 | eukaryotic translation initiation factor 5A2 |
| | LAPTM4A | lysosomal protein transmembrane 4 alpha |
| | PCDH9 | protocadherin 9 |
| | RIMKLB | ribosomal modification protein rimK like family member B |
| | PAIP1 | poly(A) binding protein interacting protein 1 |
| | RTN3 | reticulon 3 |
| | CKAP5 | cytoskeleton associated protein 5 |
| | CYP2C19 | cytochrome P450 family 2 subfamily C member 19 |
| | PGR | progesterone receptor |
| | WBP4 | WW domain binding protein 4 |
| | NUFIP2 | nuclear FMR1 interacting protein 2 |
| | KIAA0895 | KIAA0895 |
| | ARF1 | ADP ribosylation factor 1 |
| | XPNPEP3 | X-prolyl aminopeptidase 3 |
| | SLC25A33 | solute carrier family 25 member 33 |
| | MBNL2 | muscleblind like splicing regulator 2 |
| | PDE5A | phosphodiesterase 5A |
| | SETD3 | SET domain containing 3, actin histidine methyltransferase |
| | CLEC19A | C-type lectin domain containing 19A |
| | CREBRF | CREB3 regulatory factor |
| | PNISR | PNN interacting serine and arginine rich protein |
| | SCN3A | sodium voltage-gated channel alpha subunit 3 |
| | PCDH11X | protocadherin 11 X-linked |
| | SCAI | suppressor of cancer cell invasion |
| | PCDH15 | protocadherin related 15 |
| | TRDN | triadin |
| | PHYHIPL | phytanoyl-CoA 2-hydroxylase interacting protein like |
| | GPR143 | G protein-coupled receptor 143 |
| | PPM1L | protein phosphatase, Mg2+/Mn2+ dependent 1L |
| | TMTC1 | transmembrane and tetratricopeptide repeat containing 1 |
| | ATF2 | activating transcription factor 2 |
| | GTF2A1 | general transcription factor IIA subunit 1 |
| | RBMY1F | RNA binding motif protein, Y-linked, family 1, member F |
| | RBMY1A1 | RNA binding motif protein, Y-linked, family 1, member A1 |
| | RBMY1D | RNA binding motif protein, Y-linked, family 1, member D |
| | PIK3R4 | phosphoinositide-3-kinase regulatory subunit 4 |
| | PHC3 | polyhomeotic homolog 3 |

| | | |
|---|---|---|
| | SP4 | Sp4 transcription factor |
| | FSIP1 | fibrous sheath interacting protein 1 |
| | ZFAND5 | zinc finger AN1-type containing 5 |
| | MMS22L | MMS22 like, DNA repair protein |
| | NPM3 | nucleophosmin/nucleoplasmin 3 |
| | CCNA2 | cyclin A2 |
| | EP300 | E1A binding protein p300 |
| | NDFIP1 | Nedd4 family interacting protein 1 |
| | HSPA4 | heat shock protein family A (Hsp70) member 4 |
| | SRPK1 | SRSF protein kinase 1 |
| | TM9SF3 | transmembrane 9 superfamily member 3 |
| | EPHA7 | EPH receptor A7 |
| | ACSL6 | acyl-CoA synthetase long chain family member 6 |
| | PI4K2B | phosphatidylinositol 4-kinase type 2 beta |
| | LMO7 | LIM domain 7 |
| | ATXN3 | ataxin 3 |
| | USP32 | ubiquitin specific peptidase 32 |
| | C2CD4A | C2 calcium dependent domain containing 4A |
| | TRIM10 | tripartite motif containing 10 |
| | NUP133 | nucleoporin 133 |
| | CADM2 | cell adhesion molecule 2 |
| | STXBP5 | syntaxin binding protein 5 |
| | GRB14 | growth factor receptor bound protein 14 |
| | UBR5 | ubiquitin protein ligase E3 component n-recognin 5 |
| | ZNF365 | zinc finger protein 365 |
| | ATP1B4 | ATPase Na+/K+ transporting family member beta 4 |
| | EHBP1 | EH domain binding protein 1 |
| | MYO6 | myosin VI |
| | YWHAZ | tyrosine 3-monooxygenase/tryptophan 5-monooxygenase activation protein zeta |
| | TMEM168 | transmembrane protein 168 |
| | RIOK2 | RIO kinase 2 |
| | DCLRE1C | DNA cross-link repair 1C |
| | SEPTIN2 | septin 2 |
| | SCOC | short coiled-coil protein |
| | FDX1 | ferredoxin 1 |
| | SGMS1 | sphingomyelin synthase 1 |
| | PLEKHB2 | pleckstrin homology domain containing B2 |
| | USP6 | ubiquitin specific peptidase 6 |
| | CNOT6L | CCR4-NOT transcription complex subunit 6 like |
| | VAPA | VAMP associated protein A |
| | HMGA2 | high mobility group AT-hook 2 |

| | | |
|---|---|---|
| | STON1 | stonin 1 |
| | KHDRBS1 | KH RNA binding domain containing, signal transduction associated 1 |
| | CYP51A1 | cytochrome P450 family 51 subfamily A member 1 |
| | CPEB2 | cytoplasmic polyadenylation element binding protein 2 |
| | MAPK10 | mitogen-activated protein kinase 10 |
| | EEA1 | early endosome antigen 1 |
| | SYCP3 | synaptonemal complex protein 3 |
| | FAM177A1 | family with sequence similarity 177 member A1 |
| | GTPBP10 | GTP binding protein 10 |
| | RBBP8 | RB binding protein 8, endonuclease |
| | IFI44L | interferon induced protein 44 like |
| | LRRN1 | leucine rich repeat neuronal 1 |
| | CEP85L | centrosomal protein 85 like |
| | PCDH11Y | protocadherin 11 Y-linked |
| | NAMPT | nicotinamide phosphoribosyltransferase |
| | HERC4 | HECT and RLD domain containing E3 ubiquitin protein ligase 4 |
| | KMT2C | lysine methyltransferase 2C |
| | GTF3C3 | general transcription factor IIIC subunit 3 |
| | DYTN | dystrotelin |
| | SYT4 | synaptotagmin 4 |
| | POLK | DNA polymerase kappa |
| | USP28 | ubiquitin specific peptidase 28 |
| | SHANK2 | SH3 and multiple ankyrin repeat domains 2 |
| | ZNF264 | zinc finger protein 264 |
| | BARD1 | BRCA1 associated RING domain 1 |
| | RTKN2 | rhotekin 2 |
| | RB1 | RB transcriptional corepressor 1 |
| | RALYL | RALY RNA binding protein like |
| | CAPS2 | calcyphosine 2 |
| | RNF6 | ring finger protein 6 |
| | GRHL2 | grainyhead like transcription factor 2 |
| | ELF2 | E74 like ETS transcription factor 2 |
| | HCFC2 | host cell factor C2 |
| | ZFAND4 | zinc finger AN1-type containing 4 |
| | NIT2 | nitrilase family member 2 |
| | SLTM | SAFB like transcription modulator |
| | BET1 | Bet1 golgi vesicular membrane trafficking protein |
| | DCAF4L2 | DDB1 and CUL4 associated factor 4 like 2 |
| | SNRPB2 | small nuclear ribonucleoprotein polypeptide B2 |
| | PCBP2 | poly(rC) binding protein 2 |
| | MAGT1 | magnesium transporter 1 |

| | | |
|---|---|---|
| | STOM | stomatin |
| | MATN3 | matrilin 3 |
| | TMEM56 | transmembrane protein 56 |
| | R3HDM1 | R3H domain containing 1 |
| | RBM46 | RNA binding motif protein 46 |
| | PIP4P2 | phosphatidylinositol-4,5-bisphosphate 4-phosphatase 2 |
| | ME1 | malic enzyme 1 |
| | TCIM | transcriptional and immune response regulator |
| | APAF1 | apoptotic peptidase activating factor 1 |
| | SLX4IP | SLX4 interacting protein |
| | RPRD1A | regulation of nuclear pre-mRNA domain containing 1A |
| | ITGA9 | integrin subunit alpha 9 |
| | CDC16 | cell division cycle 16 |
| | USP14 | ubiquitin specific peptidase 14 |
| | SFPQ | splicing factor proline and glutamine rich |
| | ZNF518A | zinc finger protein 518A |
| | BRWD1 | bromodomain and WD repeat domain containing 1 |
| | CCDC152 | coiled-coil domain containing 152 |
| | PLS1 | plastin 1 |
| | FEM1B | fem-1 homolog B |
| | LRRC2 | leucine rich repeat containing 2 |
| | PPP2CA | protein phosphatase 2 catalytic subunit alpha |
| | KMT5B | lysine methyltransferase 5B |
| | TSPAN2 | tetraspanin 2 |
| | MARK1 | microtubule affinity regulating kinase 1 |
| | GPC6 | glypican 6 |
| | ATXN1 | ataxin 1 |
| | REEP3 | receptor accessory protein 3 |
| | RPS6KA3 | ribosomal protein S6 kinase A3 |
| | MAPKAPK2 | mitogen-activated protein kinase-activated protein kinase 2 |
| | PTH2R | parathyroid hormone 2 receptor |
| | TIMM9 | translocase of inner mitochondrial membrane 9 |
| | ARF6 | ADP ribosylation factor 6 |
| | UBA5 | ubiquitin like modifier activating enzyme 5 |
| | ZSCAN23 | zinc finger and SCAN domain containing 23 |
| | CD24 | CD24 molecule |
| | RASSF9 | Ras association domain family member 9 |
| | NBN | nibrin |
| | MS4A2 | membrane spanning 4-domains A2 |
| | UPF3B | UPF3B, regulator of nonsense mediated mRNA decay |
| | DDX5 | DEAD-box helicase 5 |
| | PLCL2 | phospholipase C like 2 |

| | | |
|---|---|---|
| | RUBCNL | rubicon like autophagy enhancer |
| | PLCH1 | phospholipase C eta 1 |
| | WDR20 | WD repeat domain 20 |
| | RPL22L1 | ribosomal protein L22 like 1 |
| | YPEL2 | yippee like 2 |
| | FAM221A | family with sequence similarity 221 member A |
| | RIMS2 | regulating synaptic membrane exocytosis 2 |
| | MANEA | mannosidase endo-alpha |
| | CFHR4 | complement factor H related 4 |
| | FBXW2 | F-box and WD repeat domain containing 2 |
| | ZBTB20 | zinc finger and BTB domain containing 20 |
| | MSL2 | MSL complex subunit 2 |
| | PRKAA2 | protein kinase AMP-activated catalytic subunit alpha 2 |
| | ATAD2B | ATPase family, AAA domain containing 2B |
| | STRBP | spermatid perinuclear RNA binding protein |
| | MTR | 5-methyltetrahydrofolate-homocysteine methyltransferase |
| | EFNA5 | ephrin A5 |
| | MIB1 | mindbomb E3 ubiquitin protein ligase 1 |
| | C1orf116 | chromosome 1 open reading frame 116 |
| | MOSMO | modulator of smoothened |
| | RBMY1E | RNA binding motif protein, Y-linked, family 1, member E |
| | FRMD4B | FERM domain containing 4B |
| | SPIN4 | spindlin family member 4 |
| Segment-3-PA- MR15 5P | NOVA1 | NOVA alternative splicing regulator 1 |
| | ADAM10 | ADAM metallopeptidase domain 10 |
| | SRSF3 | serine and arginine rich splicing factor 3 |
| | CD69 | CD69 molecule |
| | ARID1B | AT-rich interaction domain 1B |
| | PTPRK | protein tyrosine phosphatase, receptor type K |
| | EFR3A | EFR3 homolog A |
| | BICD2 | BICD cargo adaptor 2 |
| | DUSP10 | dual specificity phosphatase 10 |
| | REV3L | REV3 like, DNA directed polymerase zeta catalytic subunit |
| | TRHDE | thyrotropin releasing hormone degrading enzyme |
| | ANKS1A | ankyrin repeat and sterile alpha motif domain containing 1A |
| | ZEB2 | zinc finger E-box binding homeobox 2 |
| | ANP32E | acidic nuclear phosphoprotein 32 family member E |
| | SLC24A3 | solute carrier family 24 member 3 |
| | CPSF6 | cleavage and polyadenylation specific factor 6 |
| | TGFBR3 | transforming growth factor beta receptor 3 |
| | BACH2 | BTB domain and CNC homolog 2 |
| | DNAJB9 | DnaJ heat shock protein family (Hsp40) member B9 |

| | | |
|---|---|---|
| | TBC1D8 | TBC1 domain family member 8 |
| | SOSTDC1 | sclerostin domain containing 1 |
| | VCAM1 | vascular cell adhesion molecule 1 |
| | RNF146 | ring finger protein 146 |
| | EPC2 | enhancer of polycomb homolog 2 |
| | TENT4B | terminal nucleotidyltransferase 4B |
| | UBE2E2 | ubiquitin conjugating enzyme E2 E2 |
| | NEFH | neurofilament heavy |
| | WASL | Wiskott-Aldrich syndrome like |
| | DGKH | diacylglycerol kinase eta |
| | ZER1 | zyg-11 related cell cycle regulator |
| | SMG7 | SMG7, nonsense mediated mRNA decay factor |
| | HIPK3 | homeodomain interacting protein kinase 3 |
| | MPP1 | membrane palmitoylated protein 1 |
| | CBFA2T2 | CBFA2/RUNX1 translocation partner 2 |
| | FAM126B | family with sequence similarity 126 member B |
| | AP1G1 | adaptor related protein complex 1 subunit gamma 1 |
| | DERL1 | derlin 1 |
| | CASK | calcium/calmodulin dependent serine protein kinase |
| | TGIF1 | TGFB induced factor homeobox 1 |
| | AXDND1 | axonemal dynein light chain domain containing 1 |
| | RBMS3 | RNA binding motif single stranded interacting protein 3 |
| | MOAP1 | modulator of apoptosis 1 |
| | USP9X | ubiquitin specific peptidase 9 X-linked |
| | DNER | delta/notch like EGF repeat containing |
| | ITM2B | integral membrane protein 2B |
| | ANO6 | anoctamin 6 |
| | GRAMD1B | GRAM domain containing 1B |
| | GFPT2 | glutamine-fructose-6-phosphate transaminase 2 |
| | PTGER4 | prostaglandin E receptor 4 |
| | LHFPL2 | LHFPL tetraspan subfamily member 2 |
| | SERTAD2 | SERTA domain containing 2 |
| | ERGIC2 | ERGIC and golgi 2 |
| | CREBRF | CREB3 regulatory factor |
| | MAP1B | microtubule associated protein 1B |
| | PDP1 | pyruvate dehyrogenase phosphatase catalytic subunit 1 |
| | VDAC2 | voltage dependent anion channel 2 |
| | PHTF2 | putative homeodomain transcription factor 2 |
| | SEMA6D | semaphorin 6D |
| | LATS2 | large tumor suppressor kinase 2 |
| | ITPR1 | inositol 1,4,5-trisphosphate receptor type 1 |
| | NEO1 | neogenin 1 |

| | | |
|---|---|---|
| | B3GALT2 | beta-1,3-galactosyltransferase 2 |
| | MYO15A | myosin XVA |
| | MDGA2 | MAM domain containing glycosylphosphatidylinositol anchor 2 |
| | FAR1 | fatty acyl-CoA reductase 1 |
| | SAMD12 | sterile alpha motif domain containing 12 |
| | MTMR4 | myotubularin related protein 4 |
| | FAM43A | family with sequence similarity 43 member A |
| | EDEM1 | ER degradation enhancing alpha-mannosidase like protein 1 |
| | MRPS33 | mitochondrial ribosomal protein S33 |
| | JMY | junction mediating and regulatory protein, p53 cofactor |
| | FNIP1 | folliculin interacting protein 1 |
| | PRKCE | protein kinase C epsilon |
| | ZNF214 | zinc finger protein 214 |
| | COG3 | component of oligomeric golgi complex 3 |
| | HS3ST3B1 | heparan sulfate-glucosamine 3-sulfotransferase 3B1 |
| | CPEB2 | cytoplasmic polyadenylation element binding protein 2 |
| | OLFML2B | olfactomedin like 2B |
| | LCORL | ligand dependent nuclear receptor corepressor like |
| | NRCAM | neuronal cell adhesion molecule |
| | MOGAT2 | monoacylglycerol O-acyltransferase 2 |
| | NEURL1B | neuralized E3 ubiquitin protein ligase 1B |
| | EXOC5 | exocyst complex component 5 |
| | ERC2 | ELKS/RAB6-interacting/CAST family member 2 |
| | RFX6 | regulatory factor X6 |
| | HIVEP1 | human immunodeficiency virus type I enhancer binding protein 1 |
| | ZNF236 | zinc finger protein 236 |
| | EPB41L2 | erythrocyte membrane protein band 4.1 like 2 |
| | IL6ST | interleukin 6 signal transducer |
| | CPEB4 | cytoplasmic polyadenylation element binding protein 4 |
| | ITGA6 | integrin subunit alpha 6 |
| | SPIN1 | spindlin 1 |
| | ABR | ABR, RhoGEF and GTPase activating protein |
| | NUFIP2 | nuclear FMR1 interacting protein 2 |
| | BCL11B | BCL11B, BAF complex component |
| | TWF1 | twinfilin actin binding protein 1 |
| | CDKN1C | cyclin dependent kinase inhibitor 1C |
| | MAP2K4 | mitogen-activated protein kinase kinase 4 |
| | TSHZ1 | teashirt zinc finger homeobox 1 |
| | TMEM229A | transmembrane protein 229A |
| | SOX4 | SRY-box 4 |
| | ESR1 | estrogen receptor 1 |

| | | |
|---|---|---|
| | FOXP2 | forkhead box P2 |
| | HOXD10 | homeobox D10 |
| | HECW1 | HECT, C2 and WW domain containing E3 ubiquitin protein ligase 1 |
| | ALCAM | activated leukocyte cell adhesion molecule |
| | RPS6KA4 | ribosomal protein S6 kinase A4 |
| | NLGN4X | neuroligin 4 X-linked |
| | MOB1B | MOB kinase activator 1B |
| | BCL2L11 | BCL2 like 11 |
| | MEF2A | myocyte enhancer factor 2A |
| | ELOA | elongin A |
| | TBL1XR1 | transducin beta like 1 X-linked receptor 1 |
| | ELK4 | ELK4, ETS transcription factor |
| | AKAP1 | A-kinase anchoring protein 1 |
| | BTG2 | BTG anti-proliferation factor 2 |
| | PTF1A | pancreas associated transcription factor 1a |
| | TRMT6 | tRNA methyltransferase 6 |
| | PGS1 | phosphatidylglycerophosphate synthase 1 |
| | ZNF24 | zinc finger protein 24 |
| | ATRN | attractin |
| | ZFC3H1 | zinc finger C3H1-type containing |
| | RAB3GAP1 | RAB3 GTPase activating protein catalytic subunit 1 |
| | SSU72 | SSU72 homolog, RNA polymerase II CTD phosphatase |
| | PLEKHB2 | pleckstrin homology domain containing B2 |
| | PPP1R3F | protein phosphatase 1 regulatory subunit 3F |
| | APLF | aprataxin and PNKP like factor |
| | IDH3A | isocitrate dehydrogenase 3 (NAD(+)) alpha |
| | RIC1 | RIC1 homolog, RAB6A GEF complex partner 1 |
| | TMED5 | transmembrane p24 trafficking protein 5 |
| | ST8SIA4 | ST8 alpha-N-acetyl-neuraminide alpha-2,8-sialyltransferase 4 |
| | RFLNA | refilin A |
| | ZNF664-RFLNA | ZNF664-RFLNA readthrough |
| | CYTH3 | cytohesin 3 |
| | ARL8B | ADP ribosylation factor like GTPase 8B |
| | PRDM16 | PR/SET domain 16 |
| | RAB8B | RAB8B, member RAS oncogene family |
| | ESRP1 | epithelial splicing regulatory protein 1 |
| | P3H3 | prolyl 3-hydroxylase 3 |
| | SLC6A2 | solute carrier family 6 member 2 |
| | ABCE1 | ATP binding cassette subfamily E member 1 |
| | C3orf80 | chromosome 3 open reading frame 80 |
| | HS3ST5 | heparan sulfate-glucosamine 3-sulfotransferase 5 |

| | | |
|---|---|---|
| | FAM160B1 | family with sequence similarity 160 member B1 |
| | AMOT | angiomotin |
| | FAM135A | family with sequence similarity 135 member A |
| | KPNA3 | karyopherin subunit alpha 3 |
| | HAND2 | heart and neural crest derivatives expressed 2 |
| | RHPN2 | rhophilin Rho GTPase binding protein 2 |
| | VAMP3 | vesicle associated membrane protein 3 |
| | MS4A2 | membrane spanning 4-domains A2 |
| | NHLRC1 | NHL repeat containing E3 ubiquitin protein ligase 1 |
| | USP28 | ubiquitin specific peptidase 28 |
| | MCTP1 | multiple C2 and transmembrane domain containing 1 |
| | FKBP9 | FKBP prolyl isomerase 9 |
| | GTF2A1 | general transcription factor IIA subunit 1 |
| | KIAA1109 | KIAA1109 |
| | TOGARAM1 | TOG array regulator of axonemal microtubules 1 |
| | SCN3A | sodium voltage-gated channel alpha subunit 3 |
| | AVL9 | AVL9 cell migration associated |
| | KLF16 | Kruppel like factor 16 |
| | NSG1 | neuronal vesicle trafficking associated 1 |
| | ANO4 | anoctamin 4 |
| | AIDA | axin interactor, dorsalization associated |
| | DLG2 | discs large MAGUK scaffold protein 2 |
| | SLC44A5 | solute carrier family 44 member 5 |
| | CUX1 | cut like homeobox 1 |
| | CXorf38 | chromosome X open reading frame 38 |
| | TGFBR1 | transforming growth factor beta receptor 1 |
| | RBPMS2 | RNA binding protein, mRNA processing factor 2 |
| | MAST4 | microtubule associated serine/threonine kinase family member 4 |
| | ZSWIM5 | zinc finger SWIM-type containing 5 |
| | CACNA2D1 | calcium voltage-gated channel auxiliary subunit alpha2delta 1 |
| | PDK4 | pyruvate dehydrogenase kinase 4 |
| | DYRK1A | dual specificity tyrosine phosphorylation regulated kinase 1A |
| | PTPRD | protein tyrosine phosphatase, receptor type D |
| | VEGFB | vascular endothelial growth factor B |
| | PIK3R3 | phosphoinositide-3-kinase regulatory subunit 3 |
| | SREK1 | splicing regulatory glutamic acid and lysine rich protein 1 |
| | GID4 | GID complex subunit 4 homolog |
| | SLC32A1 | solute carrier family 32 member 1 |
| | XPR1 | xenotropic and polytropic retrovirus receptor 1 |
| | WDR82 | WD repeat domain 82 |
| | XPO4 | exportin 4 |
| | ITPRID2 | ITPR interacting domain containing 2 |

| | | |
|---|---|---|
| | FBXO28 | F-box protein 28 |
| | CXXC4 | CXXC finger protein 4 |
| | SOGA1 | suppressor of glucose, autophagy associated 1 |
| | TRIM33 | tripartite motif containing 33 |
| | KCND2 | potassium voltage-gated channel subfamily D member 2 |
| | PHACTR2 | phosphatase and actin regulator 2 |
| | PRRC2B | proline rich coiled-coil 2B |
| | MORC3 | MORC family CW-type zinc finger 3 |
| | LIMS1 | LIM zinc finger domain containing 1 |
| | SLC9A7 | solute carrier family 9 member A7 |
| | UBASH3B | ubiquitin associated and SH3 domain containing B |
| | WDR47 | WD repeat domain 47 |
| | PGK1 | phosphoglycerate kinase 1 |
| | CLDN11 | claudin 11 |
| | ST6GAL2 | ST6 beta-galactoside alpha-2,6-sialyltransferase 2 |
| | GLRA1 | glycine receptor alpha 1 |
| | CCT2 | chaperonin containing TCP1 subunit 2 |
| | TOB1 | transducer of ERBB2, 1 |
| | APPL1 | adaptor protein, phosphotyrosine interacting with PH domain and leucine zipper 1 |
| | ARHGEF3 | Rho guanine nucleotide exchange factor 3 |
| | QKI | QKI, KH domain containing RNA binding |
| | EP300 | E1A binding protein p300 |
| | UBE3A | ubiquitin protein ligase E3A |
| | STAG1 | stromal antigen 1 |
| | PEAK1 | pseudopodium enriched atypical kinase 1 |
| | TENT4A | terminal nucleotidyltransferase 4A |
| | RAD21 | RAD21 cohesin complex component |
| | ITPK1 | inositol-tetrakisphosphate 1-kinase |
| | CTDSPL | CTD small phosphatase like |
| | RIMKLA | ribosomal modification protein rimK like family member A |
| | RSBN1 | round spermatid basic protein 1 |
| | RGL1 | ral guanine nucleotide dissociation stimulator like 1 |
| | PCDH17 | protocadherin 17 |
| | KCNN3 | potassium calcium-activated channel subfamily N member 3 |
| | WASHC5 | WASH complex subunit 5 |
| | ADCYAP1 | adenylate cyclase activating polypeptide 1 |
| | MAF | MAF bZIP transcription factor |
| | RBFOX1 | RNA binding fox-1 homolog 1 |
| | ATRX | ATRX, chromatin remodeler |
| | NDUFA2 | NADH:ubiquinone oxidoreductase subunit A2 |
| | ATXN3 | ataxin 3 |

| | | |
|---|---|---|
| | USP34 | ubiquitin specific peptidase 34 |
| | CPEB3 | cytoplasmic polyadenylation element binding protein 3 |
| | SYT14 | synaptotagmin 14 |
| | SMOC2 | SPARC related modular calcium binding 2 |
| | CACNB4 | calcium voltage-gated channel auxiliary subunit beta 4 |
| | C5orf63 | chromosome 5 open reading frame 63 |
| | RBM24 | RNA binding motif protein 24 |
| | NRK | Nik related kinase |
| | SNN | stannin |
| | USP47 | ubiquitin specific peptidase 47 |
| | ITGA8 | integrin subunit alpha 8 |
| | MBNL3 | muscleblind like splicing regulator 3 |
| | CHST7 | carbohydrate sulfotransferase 7 |
| | ARHGAP24 | Rho GTPase activating protein 24 |
| | PCGF5 | polycomb group ring finger 5 |
| | DDX3X | DEAD-box helicase 3 X-linked |
| | STAT5B | signal transducer and activator of transcription 5B |
| | IGF1R | insulin like growth factor 1 receptor |
| | CTBP2 | C-terminal binding protein 2 |
| | PJA2 | praja ring finger ubiquitin ligase 2 |
| | MFF | mitochondrial fission factor |
| | LRRC17 | leucine rich repeat containing 17 |
| | LOC110117498-PIK3R3 | LOC110117498-PIK3R3 readthrough |
| | MIA3 | MIA SH3 domain ER export factor 3 |
| | CUL4A | cullin 4A |
| | SLC25A36 | solute carrier family 25 member 36 |
| | DPY30 | dpy-30, histone methyltransferase complex regulatory subunit |
| | MPL | MPL proto-oncogene, thrombopoietin receptor |
| | SLC34A3 | solute carrier family 34 member 3 |
| | SYNJ1 | synaptojanin 1 |
| | SNX13 | sorting nexin 13 |
| | UBE2W | ubiquitin conjugating enzyme E2 W |
| | TSC22D1 | TSC22 domain family member 1 |
| | NTRK2 | neurotrophic receptor tyrosine kinase 2 |
| | STAC | SH3 and cysteine rich domain |
| | DUSP5 | dual specificity phosphatase 5 |
| Segment-3-PA- MR15 3P | ASXL3 | ASXL transcriptional regulator 3 |
| | PRRC2B | proline rich coiled-coil 2B |
| | NFIB | nuclear factor I B |
| | LRRC4 | leucine rich repeat containing 4 |
| | ARMC10 | armadillo repeat containing 10 |

| | | |
|---|---|---|
| | GNPNAT1 | glucosamine-phosphate N-acetyltransferase 1 |
| | UBE2K | ubiquitin conjugating enzyme E2 K |
| | HMGXB3 | HMG-box containing 3 |
| | RTN4 | reticulon 4 |
| | SRSF1 | serine and arginine rich splicing factor 1 |
| | DIP2B | disco interacting protein 2 homolog B |
| | CRAT | carnitine O-acetyltransferase |
| | STT3A | STT3A, catalytic subunit of the oligosaccharyltransferase complex |
| | TENT5C | terminal nucleotidyltransferase 5C |
| | TMTC2 | transmembrane and tetratricopeptide repeat containing 2 |
| | CCDC127 | coiled-coil domain containing 127 |
| | CDR2L | cerebellar degeneration related protein 2 like |
| | ZBED6CL | ZBED6 C-terminal like |
| | NKTR | natural killer cell triggering receptor |
| | KDM2A | lysine demethylase 2A |
| | YWHAQ | tyrosine 3-monooxygenase/tryptophan 5-monooxygenase activation protein theta |
| | BTNL9 | butyrophilin like 9 |
| | HSPB8 | heat shock protein family B (small) member 8 |
| | ABCB7 | ATP binding cassette subfamily B member 7 |
| | KIAA0513 | KIAA0513 |
| | R3HDM1 | R3H domain containing 1 |
| | COPS6 | COP9 signalosome subunit 6 |
| | NFIA | nuclear factor I A |
| | SLC39A11 | solute carrier family 39 member 11 |
| | CAPN1 | calpain 1 |
| | RCAN2 | regulator of calcineurin 2 |
| | NPNT | nephronectin |
| | OPA1 | OPA1, mitochondrial dynamin like GTPase |
| | GATA2 | GATA binding protein 2 |
| | DDX6 | DEAD-box helicase 6 |
| | FAM53B | family with sequence similarity 53 member B |
| | ADD3 | adducin 3 |
| | NME4 | NME/NM23 nucleoside diphosphate kinase 4 |
| | SYAP1 | synapse associated protein 1 |
| | NPAS3 | neuronal PAS domain protein 3 |
| | CUX1 | cut like homeobox 1 |
| | ELAVL3 | ELAV like RNA binding protein 3 |
| | CEP85L | centrosomal protein 85 like |
| | DEDD | death effector domain containing |
| | GJC1 | gap junction protein gamma 1 |
| | FBXO21 | F-box protein 21 |

| | | |
|---|---|---|
| | MICAL3 | microtubule associated monooxygenase, calponin and LIM domain containing 3 |
| | SLC7A7 | solute carrier family 7 member 7 |
| | HIP1R | huntingtin interacting protein 1 related |
| | PROX1 | prospero homeobox 1 |
| | ADRA1A | adrenoceptor alpha 1A |
| | ZNF362 | zinc finger protein 362 |
| | ACSL1 | acyl-CoA synthetase long chain family member 1 |
| | RAPH1 | Ras association (RalGDS/AF-6) and pleckstrin homology domains 1 |
| | LOC102724265 | uncharacterized LOC102724265 |
| | SSPN | sarcospan |
| | ARF6 | ADP ribosylation factor 6 |
| | C5orf30 | chromosome 5 open reading frame 30 |
| | PAN3 | poly(A) specific ribonuclease subunit PAN3 |
| | PPP3CA | protein phosphatase 3 catalytic subunit alpha |
| | ESRP2 | epithelial splicing regulatory protein 2 |
| | ERBB2 | erb-b2 receptor tyrosine kinase 2 |
| | FXR1 | FMR1 autosomal homolog 1 |
| | ATAD2B | ATPase family, AAA domain containing 2B |
| | PPM1H | protein phosphatase, Mg2+/Mn2+ dependent 1H |
| | ADGRL4 | adhesion G protein-coupled receptor L4 |
| | LPCAT3 | lysophosphatidylcholine acyltransferase 3 |
| | CACNB2 | calcium voltage-gated channel auxiliary subunit beta 2 |
| | PPP3CC | protein phosphatase 3 catalytic subunit gamma |
| | UBE2M | ubiquitin conjugating enzyme E2 M |
| | C10orf82 | chromosome 10 open reading frame 82 |
| | SYP | synaptophysin |
| | RCN1 | reticulocalbin 1 |
| | MEF2A | myocyte enhancer factor 2A |
| | HIPK1 | homeodomain interacting protein kinase 1 |
| | TMC6 | transmembrane channel like 6 |
| | FAM155B | family with sequence similarity 155 member B |
| | IPO11 | importin 11 |
| | MTF2 | metal response element binding transcription factor 2 |
| | TCF12 | transcription factor 12 |
| | CDR2 | cerebellar degeneration related protein 2 |
| | ZNF331 | zinc finger protein 331 |
| | CHORDC1 | cysteine and histidine rich domain containing 1 |
| | TMEM240 | transmembrane protein 240 |
| | XIRP2 | xin actin binding repeat containing 2 |
| | CD160 | CD160 molecule |
| | AAK1 | AP2 associated kinase 1 |

| | | |
|---|---|---|
| | SOX4 | SRY-box 4 |
| | SCAMP5 | secretory carrier membrane protein 5 |
| | SEPTIN8 | septin 8 |
| | TRIM45 | tripartite motif containing 45 |
| | SLCO3A1 | solute carrier organic anion transporter family member 3A1 |
| | CNTN2 | contactin 2 |
| | EI24 | EI24, autophagy associated transmembrane protein |
| | IGFBP3 | insulin like growth factor binding protein 3 |
| | LRRC31 | leucine rich repeat containing 31 |
| | SHC4 | SHC adaptor protein 4 |
| | HOXB2 | homeobox B2 |
| | ERGIC1 | endoplasmic reticulum-golgi intermediate compartment 1 |
| | ANO4 | anoctamin 4 |
| | KCNJ3 | potassium voltage-gated channel subfamily J member 3 |
| | TNRC6B | trinucleotide repeat containing 6B |
| | SAR1A | secretion associated Ras related GTPase 1A |
| | PRKCI | protein kinase C iota |
| Segment-4-HA- MR9 5P | ANKRD12 | ankyrin repeat domain 12 |
| | SIX1 | SIX homeobox 1 |
| | FAM126B | family with sequence similarity 126 member B |
| | RORA | RAR related orphan receptor A |
| | HOOK1 | hook microtubule tethering protein 1 |
| | PLXNA2 | plexin A2 |
| | DNAJC1 | DnaJ heat shock protein family (Hsp40) member C1 |
| | NCKAP1 | NCK associated protein 1 |
| | AGO1 | argonaute RISC catalytic component 1 |
| | RPS6KA3 | ribosomal protein S6 kinase A3 |
| | TOB1 | transducer of ERBB2, 1 |
| | TMEM65 | transmembrane protein 65 |
| | VIL1 | villin 1 |
| | UNC5C | unc-5 netrin receptor C |
| | PCNA | proliferating cell nuclear antigen |
| | BNC2 | basonuclin 2 |
| | COL1A2 | collagen type I alpha 2 chain |
| | STK24 | serine/threonine kinase 24 |
| | MAP3K1 | mitogen-activated protein kinase kinase kinase 1 |
| | NDFIP1 | Nedd4 family interacting protein 1 |
| | TSHZ1 | teashirt zinc finger homeobox 1 |
| | EPHB1 | EPH receptor B1 |
| | CREBRF | CREB3 regulatory factor |
| | ELF2 | E74 like ETS transcription factor 2 |
| | CROT | carnitine O-octanoyltransferase |

| | | |
|---|---|---|
| | BAZ2B | bromodomain adjacent to zinc finger domain 2B |
| | ZNF77 | zinc finger protein 77 |
| | ZNF532 | zinc finger protein 532 |
| | ELP5 | elongator acetyltransferase complex subunit 5 |
| | FAM205C | family with sequence similarity 205 member C |
| | FCGR2B | Fc fragment of IgG receptor IIb |
| | PWWP2A | PWWP domain containing 2A |
| | TP53INP2 | tumor protein p53 inducible nuclear protein 2 |
| | NDUFA12 | NADH:ubiquinone oxidoreductase subunit A12 |
| | ACTR6 | ARP6 actin related protein 6 homolog |
| | CDK8 | cyclin dependent kinase 8 |
| | MYO5B | myosin VB |
| | PAH | phenylalanine hydroxylase |
| | ASB11 | ankyrin repeat and SOCS box containing 11 |
| | COL1A1 | collagen type I alpha 1 chain |
| | FAM189A1 | family with sequence similarity 189 member A1 |
| | LRRN2 | leucine rich repeat neuronal 2 |
| | HDAC3 | histone deacetylase 3 |
| | PMP22 | peripheral myelin protein 22 |
| | DNAJB6 | DnaJ heat shock protein family (Hsp40) member B6 |
| | TAF4 | TATA-box binding protein associated factor 4 |
| | ADAMTS17 | ADAM metallopeptidase with thrombospondin type 1 motif 17 |
| | ELMSAN1 | ELM2 and Myb/SANT domain containing 1 |
| | ARL13B | ADP ribosylation factor like GTPase 13B |
| | KIAA1217 | KIAA1217 |
| | KMT2E | lysine methyltransferase 2E |
| | POTEF | POTE ankyrin domain family member F |
| | TTC14 | tetratricopeptide repeat domain 14 |
| | BBS5 | Bardet-Biedl syndrome 5 |
| | ARFGEF3 | ARFGEF family member 3 |
| | UQCC1 | ubiquinol-cytochrome c reductase complex assembly factor 1 |
| | CDH2 | cadherin 2 |
| | ANKRD28 | ankyrin repeat domain 28 |
| | PPP1R3A | protein phosphatase 1 regulatory subunit 3A |
| | CLK2 | CDC like kinase 2 |
| | PPARGC1A | PPARG coactivator 1 alpha |
| | EFCAB7 | EF-hand calcium binding domain 7 |
| | KAT14 | lysine acetyltransferase 14 |
| | IDI1 | isopentenyl-diphosphate delta isomerase 1 |
| | LCORL | ligand dependent nuclear receptor corepressor like |
| | WEE1 | WEE1 G2 checkpoint kinase |
| | SLC22A23 | solute carrier family 22 member 23 |

| | | |
|---|---|---|
| | DUS4L | dihydrouridine synthase 4 like |
| | CDK19 | cyclin dependent kinase 19 |
| | MEGF9 | multiple EGF like domains 9 |
| | DYNLT3 | dynein light chain Tctex-type 3 |
| | APOO | apolipoprotein O |
| | MCMBP | minichromosome maintenance complex binding protein |
| | ZBED2 | zinc finger BED-type containing 2 |
| | SORBS1 | sorbin and SH3 domain containing 1 |
| | HABP2 | hyaluronan binding protein 2 |
| | KIF3B | kinesin family member 3B |
| | SALL1 | spalt like transcription factor 1 |
| | KHDRBS1 | KH RNA binding domain containing, signal transduction associated 1 |
| | COL4A1 | collagen type IV alpha 1 chain |
| | ATF2 | activating transcription factor 2 |
| | NT5DC3 | 5'-nucleotidase domain containing 3 |
| | ST13 | ST13, Hsp70 interacting protein |
| | SPCS2 | signal peptidase complex subunit 2 |
| | HEY2 | hes related family bHLH transcription factor with YRPW motif 2 |
| | CCDC93 | coiled-coil domain containing 93 |
| | ZNF385B | zinc finger protein 385B |
| | GRK3 | G protein-coupled receptor kinase 3 |
| | SPTBN4 | spectrin beta, non-erythrocytic 4 |
| | KCNQ5 | potassium voltage-gated channel subfamily Q member 5 |
| | ST6GALNAC5 | ST6 N-acetylgalactosaminide alpha-2,6-sialyltransferase 5 |
| | PRPS1 | phosphoribosyl pyrophosphate synthetase 1 |
| | CMTR2 | cap methyltransferase 2 |
| | C12orf65 | chromosome 12 open reading frame 65 |
| | CIAO2A | cytosolic iron-sulfur assembly component 2A |
| | ERLEC1 | endoplasmic reticulum lectin 1 |
| | S1PR2 | sphingosine-1-phosphate receptor 2 |
| | SUN1 | Sad1 and UNC84 domain containing 1 |
| | PSD3 | pleckstrin and Sec7 domain containing 3 |
| | CTTNBP2 | cortactin binding protein 2 |
| | TPM1 | tropomyosin 1 |
| | PCDH18 | protocadherin 18 |
| | SORT1 | sortilin 1 |
| Segment-4-HA-MR9 3P | SLC25A46 | solute carrier family 25 member 46 |
| | KIF1B | kinesin family member 1B |
| | IKZF2 | IKAROS family zinc finger 2 |
| | FBXL3 | F-box and leucine rich repeat protein 3 |
| | BMPR2 | bone morphogenetic protein receptor type 2 |

| | | |
|---|---|---|
| | GSK3B | glycogen synthase kinase 3 beta |
| | SOX4 | SRY-box 4 |
| | CAAP1 | caspase activity and apoptosis inhibitor 1 |
| | G3BP1 | G3BP stress granule assembly factor 1 |
| | KCNQ5 | potassium voltage-gated channel subfamily Q member 5 |
| | ABLIM1 | actin binding LIM protein 1 |
| | TIPARP | TCDD inducible poly(ADP-ribose) polymerase |
| | AEBP2 | AE binding protein 2 |
| | ARHGAP25 | Rho GTPase activating protein 25 |
| | ZNF704 | zinc finger protein 704 |
| | CACNG2 | calcium voltage-gated channel auxiliary subunit gamma 2 |
| | KIAA1210 | KIAA1210 |
| | TMCC1 | transmembrane and coiled-coil domain family 1 |
| | RRAGD | Ras related GTP binding D |
| | NEXMIF | neurite extension and migration factor |
| | WNT5A | Wnt family member 5A |
| | PTPN4 | protein tyrosine phosphatase, non-receptor type 4 |
| | UBE4B | ubiquitination factor E4B |
| | RPS6KB1 | ribosomal protein S6 kinase B1 |
| | SPOCK3 | SPARC (osteonectin), cwcv and kazal like domains proteoglycan 3 |
| | TNRC6A | trinucleotide repeat containing 6A |
| | OTX2 | orthodenticle homeobox 2 |
| | ARPC5 | actin related protein 2/3 complex subunit 5 |
| | FAM172A | family with sequence similarity 172 member A |
| | KPNA6 | karyopherin subunit alpha 6 |
| | TPD52L3 | TPD52 like 3 |
| | ETFRF1 | electron transfer flavoprotein regulatory factor 1 |
| | PKD2 | polycystin 2, transient receptor potential cation channel |
| | PIEZO2 | piezo type mechanosensitive ion channel component 2 |
| | XKR6 | XK related 6 |
| | C15orf41 | chromosome 15 open reading frame 41 |
| | IL17A | interleukin 17A |
| | PPP1R3B | protein phosphatase 1 regulatory subunit 3B |
| | PHAX | phosphorylated adaptor for RNA export |
| | XPO1 | exportin 1 |
| | CDC73 | cell division cycle 73 |
| | TMPRSS11E | transmembrane serine protease 11E |
| | MBNL3 | muscleblind like splicing regulator 3 |
| | STXBP5 | syntaxin binding protein 5 |
| | CEP120 | centrosomal protein 120 |
| | FGF12 | fibroblast growth factor 12 |

| | | |
|---|---|---|
| | SGK3 | serum/glucocorticoid regulated kinase family member 3 |
| | PTN | pleiotrophin |
| | MINDY2 | MINDY lysine 48 deubiquitinase 2 |
| | LIN28B | lin-28 homolog B |
| | COL4A4 | collagen type IV alpha 4 chain |
| | MXI1 | MAX interactor 1, dimerization protein |
| | CAST | calpastatin |
| | ZDHHC21 | zinc finger DHHC-type containing 21 |
| | FMR1 | fragile X mental retardation 1 |
| | MTURN | maturin, neural progenitor differentiation regulator homolog |
| | EDNRB | endothelin receptor type B |
| | FAM20B | FAM20B, glycosaminoglycan xylosylkinase |
| | NEGR1 | neuronal growth regulator 1 |
| | ZNF106 | zinc finger protein 106 |
| | RBM41 | RNA binding motif protein 41 |
| | LAMC1 | laminin subunit gamma 1 |
| | KLHL32 | kelch like family member 32 |
| | C6orf47 | chromosome 6 open reading frame 47 |
| | ARHGEF10L | Rho guanine nucleotide exchange factor 10 like |
| | SCN1A | sodium voltage-gated channel alpha subunit 1 |
| | MAPRE1 | microtubule associated protein RP/EB family member 1 |
| | CUL4B | cullin 4B |
| | PDZRN4 | PDZ domain containing ring finger 4 |
| | NAA30 | N(alpha)-acetyltransferase 30, NatC catalytic subunit |
| | CREB5 | cAMP responsive element binding protein 5 |
| | NPEPPS | aminopeptidase puromycin sensitive |
| | PHACTR2 | phosphatase and actin regulator 2 |
| | GUCY1A2 | guanylate cyclase 1 soluble subunit alpha 2 |
| | SPAG1 | sperm associated antigen 1 |
| | C1QTNF7 | C1q and TNF related 7 |
| | TMOD2 | tropomodulin 2 |
| | COCH | cochlin |
| | MED30 | mediator complex subunit 30 |
| | HDGFL3 | HDGF like 3 |
| | YTHDF3 | YTH N6-methyladenosine RNA binding protein 3 |
| | KLHL28 | kelch like family member 28 |
| | SNAPC3 | small nuclear RNA activating complex polypeptide 3 |
| | ETF1 | eukaryotic translation termination factor 1 |
| | LRRTM2 | leucine rich repeat transmembrane neuronal 2 |
| | GRIP2 | glutamate receptor interacting protein 2 |
| | PCDH7 | protocadherin 7 |

| | | |
|---|---|---|
| | B3GNT3 | UDP-GlcNAc:betaGal beta-1,3-N-acetylglucosaminyltransferase 3 |
| | TBC1D8B | TBC1 domain family member 8B |
| | C18orf63 | chromosome 18 open reading frame 63 |
| | THAP12 | THAP domain containing 12 |
| | AKTIP | AKT interacting protein |
| | MARVELD3 | MARVEL domain containing 3 |
| | ZCCHC9 | zinc finger CCHC-type containing 9 |
| | GABRG1 | gamma-aminobutyric acid type A receptor gamma1 subunit |
| | TNPO1 | transportin 1 |
| | PRKAA2 | protein kinase AMP-activated catalytic subunit alpha 2 |
| | KLF5 | Kruppel like factor 5 |
| | FBXO30 | F-box protein 30 |
| | PROX1 | prospero homeobox 1 |
| | OPRM1 | opioid receptor mu 1 |
| | PICALM | phosphatidylinositol binding clathrin assembly protein |
| | GLUL | glutamate-ammonia ligase |
| | BTLA | B and T lymphocyte associated |
| | SLC15A5 | solute carrier family 15 member 5 |
| | CTNNA2 | catenin alpha 2 |
| | SH3KBP1 | SH3 domain containing kinase binding protein 1 |
| | INO80D | INO80 complex subunit D |
| | PLPPR4 | phospholipid phosphatase related 4 |
| | CHEK1 | checkpoint kinase 1 |
| | NSD3 | nuclear receptor binding SET domain protein 3 |
| | GABRB3 | gamma-aminobutyric acid type A receptor beta3 subunit |
| | RALGAPA2 | Ral GTPase activating protein catalytic alpha subunit 2 |
| | API5 | apoptosis inhibitor 5 |
| | SCN8A | sodium voltage-gated channel alpha subunit 8 |
| | DNM3 | dynamin 3 |
| | DIP2B | disco interacting protein 2 homolog B |
| | ZNF280D | zinc finger protein 280D |
| | TMEM19 | transmembrane protein 19 |
| | WDSUB1 | WD repeat, sterile alpha motif and U-box domain containing 1 |
| | BTBD11 | BTB domain containing 11 |
| | IGF1 | insulin like growth factor 1 |
| | MFSD4A | major facilitator superfamily domain containing 4A |
| | UBE2A | ubiquitin conjugating enzyme E2 A |
| | BCL11B | BCL11B, BAF complex component |
| | SESTD1 | SEC14 and spectrin domain containing 1 |
| | TTC17 | tetratricopeptide repeat domain 17 |
| | CDR2L | cerebellar degeneration related protein 2 like |

| | | |
|---|---|---|
| | RNF13 | ring finger protein 13 |
| | C8orf44-SGK3 | C8orf44-SGK3 readthrough |
| | CENPS | centromere protein S |
| | SLC2A13 | solute carrier family 2 member 13 |
| | ABCE1 | ATP binding cassette subfamily E member 1 |
| | LHX9 | LIM homeobox 9 |
| | ACOT12 | acyl-CoA thioesterase 12 |
| | RIF1 | replication timing regulatory factor 1 |
| | AHDC1 | AT-hook DNA binding motif containing 1 |
| | MATN2 | matrilin 2 |
| | KCNMB2 | potassium calcium-activated channel subfamily M regulatory beta subunit 2 |
| | HDAC9 | histone deacetylase 9 |
| | RIPOR1 | RHO family interacting cell polarization regulator 1 |
| | ZNF197 | zinc finger protein 197 |
| | CPEB2 | cytoplasmic polyadenylation element binding protein 2 |
| | TRAF3IP1 | TRAF3 interacting protein 1 |
| | CEPT1 | choline/ethanolamine phosphotransferase 1 |
| | FXN | frataxin |
| | SLC16A9 | solute carrier family 16 member 9 |
| | PRDM6 | PR/SET domain 6 |
| | SIX3 | SIX homeobox 3 |
| | STRIP1 | striatin interacting protein 1 |
| | VPS13A | vacuolar protein sorting 13 homolog A |
| | ATF7IP | activating transcription factor 7 interacting protein |
| | KCNB1 | potassium voltage-gated channel subfamily B member 1 |
| | PHF6 | PHD finger protein 6 |
| | PLXDC2 | plexin domain containing 2 |
| | PHF14 | PHD finger protein 14 |
| | LPGAT1 | lysophosphatidylglycerol acyltransferase 1 |
| | CISD1 | CDGSH iron sulfur domain 1 |
| | SHISA9 | shisa family member 9 |
| | NDST3 | N-deacetylase and N-sulfotransferase 3 |
| | NUP50 | nucleoporin 50 |
| | GRPEL2 | GrpE like 2, mitochondrial |
| | CAMK4 | calcium/calmodulin dependent protein kinase IV |
| | EPM2AIP1 | EPM2A interacting protein 1 |
| | PTPRT | protein tyrosine phosphatase, receptor type T |
| | ICA1L | islet cell autoantigen 1 like |
| | SETX | senataxin |
| | CALCR | calcitonin receptor |
| | CLOCK | clock circadian regulator |

| | | |
|---|---|---|
| | TMED6 | transmembrane p24 trafficking protein 6 |
| | PCDH19 | protocadherin 19 |
| | TMEM56 | transmembrane protein 56 |
| | TARP | TCR gamma alternate reading frame protein |
| | ARHGAP42 | Rho GTPase activating protein 42 |
| | GPC6 | glypican 6 |
| | CDCA2 | cell division cycle associated 2 |
| | IRX3 | iroquois homeobox 3 |
| | FAM83B | family with sequence similarity 83 member B |
| | C1orf21 | chromosome 1 open reading frame 21 |
| | RPGRIP1L | RPGRIP1 like |
| | GAN | gigaxonin |
| | CWC27 | CWC27 spliceosome associated protein homolog |
| | ZNF480 | zinc finger protein 480 |
| | MLXIPL | MLX interacting protein like |
| | PIK3R1 | phosphoinositide-3-kinase regulatory subunit 1 |
| | SELENOI | selenoprotein I |
| | CNTNAP2 | contactin associated protein like 2 |
| | PYM1 | PYM homolog 1, exon junction complex associated factor |
| | VXN | vexin |
| | CREB3L1 | cAMP responsive element binding protein 3 like 1 |
| | C17orf112 | chromosome 17 open reading frame 112 |
| | HOOK3 | hook microtubule tethering protein 3 |
| | CDC42SE1 | CDC42 small effector 1 |
| | GCA | grancalcin |
| | IVD | isovaleryl-CoA dehydrogenase |
| | UGT2B15 | UDP glucuronosyltransferase family 2 member B15 |
| | CENPP | centromere protein P |
| | CACYBP | calcyclin binding protein |
| | A1CF | APOBEC1 complementation factor |
| | ETV1 | ETS variant 1 |
| | CADM2 | cell adhesion molecule 2 |
| | NFATC3 | nuclear factor of activated T cells 3 |
| | ANGPTL1 | angiopoietin like 1 |
| | ZCCHC14 | zinc finger CCHC-type containing 14 |
| | TEPP | testis, prostate and placenta expressed |
| | TRABD2A | TraB domain containing 2A |
| | FCRL4 | Fc receptor like 4 |
| | PUM2 | pumilio RNA binding family member 2 |
| | SLC35F1 | solute carrier family 35 member F1 |
| | RAPGEF4 | Rap guanine nucleotide exchange factor 4 |
| | ZNF329 | zinc finger protein 329 |

| | | |
|---|---|---|
| | SLC48A1 | solute carrier family 48 member 1 |
| | RGS17 | regulator of G protein signaling 17 |
| | PLK4 | polo like kinase 4 |
| | ZNF587 | zinc finger protein 587 |
| | NRXN1 | neurexin 1 |
| | DUSP19 | dual specificity phosphatase 19 |
| | ADARB2 | adenosine deaminase, RNA specific B2 (inactive) |
| | DALRD3 | DALR anticodon binding domain containing 3 |
| | PTPN12 | protein tyrosine phosphatase, non-receptor type 12 |
| | BCL6 | BCL6, transcription repressor |
| | PHLDB2 | pleckstrin homology like domain family B member 2 |
| | AFAP1L2 | actin filament associated protein 1 like 2 |
| | EPHA5 | EPH receptor A5 |
| | GXYLT1 | glucoside xylosyltransferase 1 |
| | CAPRIN1 | cell cycle associated protein 1 |
| | GYPA | glycophorin A (MNS blood group) |
| | FAM120C | family with sequence similarity 120C |
| | EP300 | E1A binding protein p300 |
| | CPSF6 | cleavage and polyadenylation specific factor 6 |
| | GPR183 | G protein-coupled receptor 183 |
| | BNIP3 | BCL2 interacting protein 3 |
| | CD55 | CD55 molecule (Cromer blood group) |
| | GTF3C4 | general transcription factor IIIC subunit 4 |
| | OVOL2 | ovo like zinc finger 2 |
| | GAPVD1 | GTPase activating protein and VPS9 domains 1 |
| | CELF5 | CUGBP Elav-like family member 5 |
| | NRG2 | neuregulin 2 |
| | NAB1 | NGFI-A binding protein 1 |
| | LOXHD1 | lipoxygenase homology domains 1 |
| | SYT14 | synaptotagmin 14 |
| | TFB1M | transcription factor B1, mitochondrial |
| | TET1 | tet methylcytosine dioxygenase 1 |
| | PLXNA2 | plexin A2 |
| | DPYSL2 | dihydropyrimidinase like 2 |
| | DENND1B | DENN domain containing 1B |
| | MAP3K20 | mitogen-activated protein kinase kinase kinase 20 |
| | SLC16A12 | solute carrier family 16 member 12 |
| | LMBR1 | limb development membrane protein 1 |
| | ARID3A | AT-rich interaction domain 3A |
| | PIGT | phosphatidylinositol glycan anchor biosynthesis class T |
| | MAP2 | microtubule associated protein 2 |
| | MEDAG | mesenteric estrogen dependent adipogenesis |

| | | |
|---|---|---|
| | EYS | eyes shut homolog |
| | SAMD4A | sterile alpha motif domain containing 4A |
| | RCOR3 | REST corepressor 3 |
| | CER1 | cerberus 1, DAN family BMP antagonist |
| | PIGBOS1 | PIGB opposite strand 1 |
| | SOBP | sine oculis binding protein homolog |
| | LPAR4 | lysophosphatidic acid receptor 4 |
| | DAZL | deleted in azoospermia like |
| | RALGDS | ral guanine nucleotide dissociation stimulator |
| | TAOK3 | TAO kinase 3 |
| | GUF1 | GUF1 homolog, GTPase |
| | EPN2 | epsin 2 |
| | PTPN14 | protein tyrosine phosphatase, non-receptor type 14 |
| | PAPOLB | poly(A) polymerase beta |
| | CNTN1 | contactin 1 |
| | PPM1L | protein phosphatase, Mg2+/Mn2+ dependent 1L |
| | MED13 | mediator complex subunit 13 |
| | MEPE | matrix extracellular phosphoglycoprotein |
| | SPIRE1 | spire type actin nucleation factor 1 |
| | ZCWPW2 | zinc finger CW-type and PWWP domain containing 2 |
| | COQ10B | coenzyme Q10B |
| | ABCA5 | ATP binding cassette subfamily A member 5 |
| | BCKDHB | branched chain keto acid dehydrogenase E1 subunit beta |
| | SRPK1 | SRSF protein kinase 1 |
| | KDM4C | lysine demethylase 4C |
| | AHSA2P | activator of HSP90 ATPase homolog 2, pseudogene |
| | CCDC86 | coiled-coil domain containing 86 |
| | EGR3 | early growth response 3 |
| | ALDH1L2 | aldehyde dehydrogenase 1 family member L2 |
| | AMMECR1 | Alport syndrome, mental retardation, midface hypoplasia and elliptocytosis chromosomal region gene 1 |
| | KIAA1143 | KIAA1143 |
| | SLC25A24 | solute carrier family 25 member 24 |
| | STK39 | serine/threonine kinase 39 |
| | ZBTB34 | zinc finger and BTB domain containing 34 |
| | SPRY3 | sprouty RTK signaling antagonist 3 |
| | FAM161A | FAM161A, centrosomal protein |
| | FAM110B | family with sequence similarity 110 member B |
| | CNTN5 | contactin 5 |
| | KCNIP4 | potassium voltage-gated channel interacting protein 4 |
| | TTC28 | tetratricopeptide repeat domain 28 |
| | MCUR1 | mitochondrial calcium uniporter regulator 1 |

| | | |
|---|---|---|
| | SETMAR | SET domain and mariner transposase fusion gene |
| | ARL6IP6 | ADP ribosylation factor like GTPase 6 interacting protein 6 |
| | RAB39A | RAB39A, member RAS oncogene family |
| | EFEMP1 | EGF containing fibulin extracellular matrix protein 1 |
| | KIF26A | kinesin family member 26A |
| | TRIM5 | tripartite motif containing 5 |
| | ZNHIT3 | zinc finger HIT-type containing 3 |
| | SORBS1 | sorbin and SH3 domain containing 1 |
| | RBMX | RNA binding motif protein X-linked |
| | FLT1 | fms related tyrosine kinase 1 |
| | FERMT2 | fermitin family member 2 |
| | SPATA13 | spermatogenesis associated 13 |
| | RETREG1 | reticulophagy regulator 1 |
| | RNF207 | ring finger protein 207 |
| | EIF3B | eukaryotic translation initiation factor 3 subunit B |
| | VEZF1 | vascular endothelial zinc finger 1 |
| | NPAP1 | nuclear pore associated protein 1 |
| | TMEM215 | transmembrane protein 215 |
| | LMNA | lamin A/C |
| | CEP97 | centrosomal protein 97 |
| | ILDR2 | immunoglobulin like domain containing receptor 2 |
| | METTL8 | methyltransferase like 8 |
| | TIGAR | TP53 induced glycolysis regulatory phosphatase |
| | TMPO | thymopoietin |
| | JADE1 | jade family PHD finger 1 |
| | QDPR | quinoid dihydropteridine reductase |
| | MACO1 | macoilin 1 |
| | TMED4 | transmembrane p24 trafficking protein 4 |
| | MPP7 | membrane palmitoylated protein 7 |
| | ERBIN | erbb2 interacting protein |
| | PF4V1 | platelet factor 4 variant 1 |
| | CXCL6 | C-X-C motif chemokine ligand 6 |
| | FICD | FIC domain containing |
| | EPG5 | ectopic P-granules autophagy protein 5 homolog |
| | TENT2 | terminal nucleotidyltransferase 2 |
| | RAB30 | RAB30, member RAS oncogene family |
| | DMTF1 | cyclin D binding myb like transcription factor 1 |
| | SMAD9 | SMAD family member 9 |
| | ERCC4 | ERCC excision repair 4, endonuclease catalytic subunit |
| | DCUN1D5 | defective in cullin neddylation 1 domain containing 5 |
| | C11orf71 | chromosome 11 open reading frame 71 |
| | BLZF1 | basic leucine zipper nuclear factor 1 |

| | | |
|---|---|---|
| | PRPF4 | pre-mRNA processing factor 4 |
| | AGGF1 | angiogenic factor with G-patch and FHA domains 1 |
| | LRBA | LPS responsive beige-like anchor protein |
| | TFDP2 | transcription factor Dp-2 |
| | PTMA | prothymosin alpha |
| | FZD10 | frizzled class receptor 10 |
| | ACOT13 | acyl-CoA thioesterase 13 |
| | DSG3 | desmoglein 3 |
| | ISCU | iron-sulfur cluster assembly enzyme |
| | CDS1 | CDP-diacylglycerol synthase 1 |
| | MGAT4A | alpha-1,3-mannosyl-glycoprotein 4-beta-N-acetylglucosaminyltransferase A |
| | LYSMD3 | LysM domain containing 3 |
| | CNST | consortin, connexin sorting protein |
| | EMP1 | epithelial membrane protein 1 |
| | MDM1 | Mdm1 nuclear protein |
| | C2orf88 | chromosome 2 open reading frame 88 |
| | ZNF570 | zinc finger protein 570 |
| | BRD3 | bromodomain containing 3 |
| | PCSK2 | proprotein convertase subtilisin/kexin type 2 |
| | SNX1 | sorting nexin 1 |
| | ASPN | asporin |
| | PHEX | phosphate regulating endopeptidase homolog X-linked |
| | GBP7 | guanylate binding protein 7 |
| | CAB39 | calcium binding protein 39 |
| | LIMCH1 | LIM and calponin homology domains 1 |
| | MME | membrane metalloendopeptidase |
| | RAB23 | RAB23, member RAS oncogene family |
| | OTP | orthopedia homeobox |
| | NIPSNAP3B | nipsnap homolog 3B |
| | ITGA4 | integrin subunit alpha 4 |
| | ATP9A | ATPase phospholipid transporting 9A (putative) |
| | CELA1 | chymotrypsin like elastase family member 1 |
| | HORMAD1 | HORMA domain containing 1 |
| | MYT1 | myelin transcription factor 1 |
| | NAALADL2 | N-acetylated alpha-linked acidic dipeptidase like 2 |
| | COL14A1 | collagen type XIV alpha 1 chain |
| | HMGA2 | high mobility group AT-hook 2 |
| | ZNF366 | zinc finger protein 366 |
| | RAB3B | RAB3B, member RAS oncogene family |
| | OTX1 | orthodenticle homeobox 1 |
| | BBS12 | Bardet-Biedl syndrome 12 |

| | | |
|---|---|---|
| | VGLL3 | vestigial like family member 3 |
| | MED6 | mediator complex subunit 6 |
| | FAM81B | family with sequence similarity 81 member B |
| | CHL1 | cell adhesion molecule L1 like |
| | FBXO28 | F-box protein 28 |
| | SERINC3 | serine incorporator 3 |
| | PLCXD3 | phosphatidylinositol specific phospholipase C X domain containing 3 |
| | BORCS7 | BLOC-1 related complex subunit 7 |
| | CFLAR | CASP8 and FADD like apoptosis regulator |
| | SPAG9 | sperm associated antigen 9 |
| | PLCL1 | phospholipase C like 1 (inactive) |
| | ZNF138 | zinc finger protein 138 |
| | TRIM2 | tripartite motif containing 2 |
| | FAT3 | FAT atypical cadherin 3 |
| | TBC1D24 | TBC1 domain family member 24 |
| | NEDD4L | neural precursor cell expressed, developmentally down-regulated 4-like, E3 ubiquitin protein ligase |
| | RNF125 | ring finger protein 125 |
| | RPF1 | ribosome production factor 1 homolog |
| | CCDC59 | coiled-coil domain containing 59 |
| | BICRAL | BRD4 interacting chromatin remodeling complex associated protein like |
| | MIB1 | mindbomb E3 ubiquitin protein ligase 1 |
| | POU3F2 | POU class 3 homeobox 2 |
| | RCC2 | regulator of chromosome condensation 2 |
| | ZDHHC23 | zinc finger DHHC-type containing 23 |
| | SCARA5 | scavenger receptor class A member 5 |
| | ARHGAP32 | Rho GTPase activating protein 32 |
| | CSNK1G1 | casein kinase 1 gamma 1 |
| | AIPL1 | aryl hydrocarbon receptor interacting protein like 1 |
| | IL12B | interleukin 12B |
| | ZNF345 | zinc finger protein 345 |
| | C11orf1 | chromosome 11 open reading frame 1 |
| | HDAC2 | histone deacetylase 2 |
| | BCL2 | BCL2, apoptosis regulator |
| | CSTF3 | cleavage stimulation factor subunit 3 |
| | LHX3 | LIM homeobox 3 |
| | ASH1L | ASH1 like histone lysine methyltransferase |
| | GLRA3 | glycine receptor alpha 3 |
| | NFIX | nuclear factor I X |
| | ARSK | arylsulfatase family member K |
| | PALM2 | paralemmin 2 |

| | | |
|---|---|---|
| | SLC24A2 | solute carrier family 24 member 2 |
| | CEP135 | centrosomal protein 135 |
| | POF1B | POF1B, actin binding protein |
| | CCND1 | cyclin D1 |
| | CDH20 | cadherin 20 |
| | PPP2R3C | protein phosphatase 2 regulatory subunit B"gamma |
| | CGGBP1 | CGG triplet repeat binding protein 1 |
| | FBXW2 | F-box and WD repeat domain containing 2 |
| | NUDT4 | nudix hydrolase 4 |
| Segment-7-M- MR5 5P | ZNF213 | zinc finger protein 213 |
| | TRHDE | thyrotropin releasing hormone degrading enzyme |
| | USP46 | ubiquitin specific peptidase 46 |
| | PIK3R5 | phosphoinositide-3-kinase regulatory subunit 5 |
| | ZNF827 | zinc finger protein 827 |
| | PFKP | phosphofructokinase, platelet |
| | PDIK1L | PDLIM1 interacting kinase 1 like |
| | TNKS2 | tankyrase 2 |
| | ZC3H12C | zinc finger CCCH-type containing 12C |
| | ATXN7 | ataxin 7 |
| | PPARA | peroxisome proliferator activated receptor alpha |
| | GXYLT1 | glucoside xylosyltransferase 1 |
| | GAS2 | growth arrest specific 2 |
| | CCDC6 | coiled-coil domain containing 6 |
| | DOCK3 | dedicator of cytokinesis 3 |
| | ARL5B | ADP ribosylation factor like GTPase 5B |
| | ATXN1 | ataxin 1 |
| | NR2E1 | nuclear receptor subfamily 2 group E member 1 |
| | IL6ST | interleukin 6 signal transducer |
| | ADAMTSL3 | ADAMTS like 3 |
| | CDK19 | cyclin dependent kinase 19 |
| | ZNF711 | zinc finger protein 711 |
| | SPOP | speckle type BTB/POZ protein |
| | DYRK2 | dual specificity tyrosine phosphorylation regulated kinase 2 |
| | CAPRIN2 | caprin family member 2 |
| | SLAIN2 | SLAIN motif family member 2 |
| | BICC1 | BicC family RNA binding protein 1 |
| | TTC23 | tetratricopeptide repeat domain 23 |
| | SLC16A6 | solute carrier family 16 member 6 |
| | MAPK10 | mitogen-activated protein kinase 10 |
| | ESRP2 | epithelial splicing regulatory protein 2 |
| | CYP2U1 | cytochrome P450 family 2 subfamily U member 1 |
| | AKTIP | AKT interacting protein |

| | | |
|---|---|---|
| | PTBP2 | polypyrimidine tract binding protein 2 |
| | TM2D3 | TM2 domain containing 3 |
| | SELENOT | selenoprotein T |
| | CAMTA1 | calmodulin binding transcription activator 1 |
| | PTCHD1 | patched domain containing 1 |
| | MYT1L | myelin transcription factor 1 like |
| | QSER1 | glutamine and serine rich 1 |
| | CUBN | cubilin |
| | TP53INP1 | tumor protein p53 inducible nuclear protein 1 |
| | FAT4 | FAT atypical cadherin 4 |
| | AHRR | aryl-hydrocarbon receptor repressor |
| | AGTR2 | angiotensin II receptor type 2 |
| | PRKAR2A | protein kinase cAMP-dependent type II regulatory subunit alpha |
| | LRRC8B | leucine rich repeat containing 8 VRAC subunit B |
| | FEM1C | fem-1 homolog C |
| | C1orf21 | chromosome 1 open reading frame 21 |
| | RND3 | Rho family GTPase 3 |
| | HLF | HLF, PAR bZIP transcription factor |
| | ADAM18 | ADAM metallopeptidase domain 18 |
| | DDI1 | DNA damage inducible 1 homolog 1 |
| | SNRNP200 | small nuclear ribonucleoprotein U5 subunit 200 |
| | CPM | carboxypeptidase M |
| | ABHD18 | abhydrolase domain containing 18 |
| | GON4L | gon-4 like |
| | PHACTR4 | phosphatase and actin regulator 4 |
| | ZNF12 | zinc finger protein 12 |
| | JMJD1C | jumonji domain containing 1C |
| | HPRT1 | hypoxanthine phosphoribosyltransferase 1 |
| | ERBB3 | erb-b2 receptor tyrosine kinase 3 |
| | SLC6A11 | solute carrier family 6 member 11 |
| | PPP4R3B | protein phosphatase 4 regulatory subunit 3B |
| | RNF38 | ring finger protein 38 |
| | PLPP4 | phospholipid phosphatase 4 |
| | CYSTM1 | cysteine rich transmembrane module containing 1 |
| | EFCAB5 | EF-hand calcium binding domain 5 |
| | ABL2 | ABL proto-oncogene 2, non-receptor tyrosine kinase |
| | ZDHHC3 | zinc finger DHHC-type containing 3 |
| | ZFR | zinc finger RNA binding protein |
| | CUX1 | cut like homeobox 1 |
| | TRAK1 | trafficking kinesin protein 1 |
| | USP3 | ubiquitin specific peptidase 3 |
| | TP63 | tumor protein p63 |

| | | |
|---|---|---|
| | MLLT6 | MLLT6, PHD finger containing |
| | EPC1 | enhancer of polycomb homolog 1 |
| | ZFHX3 | zinc finger homeobox 3 |
| | PFN2 | profilin 2 |
| | FOXK2 | forkhead box K2 |
| | ARL17A | ADP ribosylation factor like GTPase 17A |
| | ZFYVE16 | zinc finger FYVE-type containing 16 |
| | ADCY1 | adenylate cyclase 1 |
| | SMAD9 | SMAD family member 9 |
| | TMCO6 | transmembrane and coiled-coil domains 6 |
| | ARHGEF10 | Rho guanine nucleotide exchange factor 10 |
| | MARCHF8 | membrane associated ring-CH-type finger 8 |
| | FBXL17 | F-box and leucine rich repeat protein 17 |
| | KLB | klotho beta |
| | THRB | thyroid hormone receptor beta |
| | PCYT1B | phosphate cytidylyltransferase 1, choline, beta |
| | FBN1 | fibrillin 1 |
| | KAT6A | lysine acetyltransferase 6A |
| | PTPN4 | protein tyrosine phosphatase, non-receptor type 4 |
| | TTPAL | alpha tocopherol transfer protein like |
| | PPP3CA | protein phosphatase 3 catalytic subunit alpha |
| | KLF7 | Kruppel like factor 7 |
| | KCNRG | potassium channel regulator |
| | PRR32 | proline rich 32 |
| | MAST4 | microtubule associated serine/threonine kinase family member 4 |
| | PLLP | plasmolipin |
| | SSX4B | SSX family member 4B |
| | FGL2 | fibrinogen like 2 |
| | PCDH10 | protocadherin 10 |
| | FGFR1OP | FGFR1 oncogene partner |
| | CSGALNACT2 | chondroitin sulfate N-acetylgalactosaminyltransferase 2 |
| | TNRC6B | trinucleotide repeat containing 6B |
| | HCFC2 | host cell factor C2 |
| | SFMBT1 | Scm like with four mbt domains 1 |
| | NUP88 | nucleoporin 88 |
| | CADM2 | cell adhesion molecule 2 |
| | NOD1 | nucleotide binding oligomerization domain containing 1 |
| | RABEP1 | rabaptin, RAB GTPase binding effector protein 1 |
| | UBL3 | ubiquitin like 3 |
| | ZNF24 | zinc finger protein 24 |
| | CTC1 | CST telomere replication complex component 1 |

| | | |
|---|---|---|
| | TANC1 | tetratricopeptide repeat, ankyrin repeat and coiled-coil containing 1 |
| | MLX | MLX, MAX dimerization protein |
| | TNFAIP1 | TNF alpha induced protein 1 |
| | CXXC5 | CXXC finger protein 5 |
| | KIN | Kin17 DNA and RNA binding protein |
| | RNF17 | ring finger protein 17 |
| | PXYLP1 | 2-phosphoxylose phosphatase 1 |
| | NFIB | nuclear factor I B |
| | ARHGEF3 | Rho guanine nucleotide exchange factor 3 |
| | GIPC2 | GIPC PDZ domain containing family member 2 |
| | CDS1 | CDP-diacylglycerol synthase 1 |
| | AHNAK | AHNAK nucleoprotein |
| | ALX4 | ALX homeobox 4 |
| Segment-7-M- MR5 3P | CGN | cingulin |
| | TTF2 | transcription termination factor 2 |
| | C17orf107 | chromosome 17 open reading frame 107 |
| | HECTD2 | HECT domain E3 ubiquitin protein ligase 2 |
| | ALDH6A1 | aldehyde dehydrogenase 6 family member A1 |
| | ZNF747 | zinc finger protein 747 |
| | CASD1 | CAS1 domain containing 1 |
| | ADH1B | alcohol dehydrogenase 1B (class I), beta polypeptide |
| | EIF2AK3 | eukaryotic translation initiation factor 2 alpha kinase 3 |
| | BCAM | basal cell adhesion molecule (Lutheran blood group) |
| | ZNF860 | zinc finger protein 860 |
| | CNGB3 | cyclic nucleotide gated channel beta 3 |
| | IGFBP7 | insulin like growth factor binding protein 7 |
| | CCDC189 | coiled-coil domain containing 189 |
| | MECP2 | methyl-CpG binding protein 2 |
| | FAM192A | family with sequence similarity 192 member A |
| | SRGAP2 | SLIT-ROBO Rho GTPase activating protein 2 |
| | RPL10 | ribosomal protein L10 |
| | ADIRF | adipogenesis regulatory factor |
| | OPALIN | oligodendrocytic myelin paranodal and inner loop protein |
| | WAPL | WAPL cohesin release factor |
| | FRMD3 | FERM domain containing 3 |
| | DEFB125 | defensin beta 125 |
| | NKAIN1 | sodium/potassium transporting ATPase interacting 1 |
| | SLAMF6 | SLAM family member 6 |
| | RSPO1 | R-spondin 1 |
| | SLC9A1 | solute carrier family 9 member A1 |
| | ALDH1A2 | aldehyde dehydrogenase 1 family member A2 |

| | | |
|---|---|---|
| | ZNF813 | zinc finger protein 813 |
| | PLB1 | phospholipase B1 |
| | UFL1 | UFM1 specific ligase 1 |
| | ATP2B2 | ATPase plasma membrane Ca2+ transporting 2 |
| | TMEM203 | transmembrane protein 203 |
| | FAM227B | family with sequence similarity 227 member B |
| | ZC3H11A | zinc finger CCCH-type containing 11A |
| | TRDMT1 | tRNA aspartic acid methyltransferase 1 |

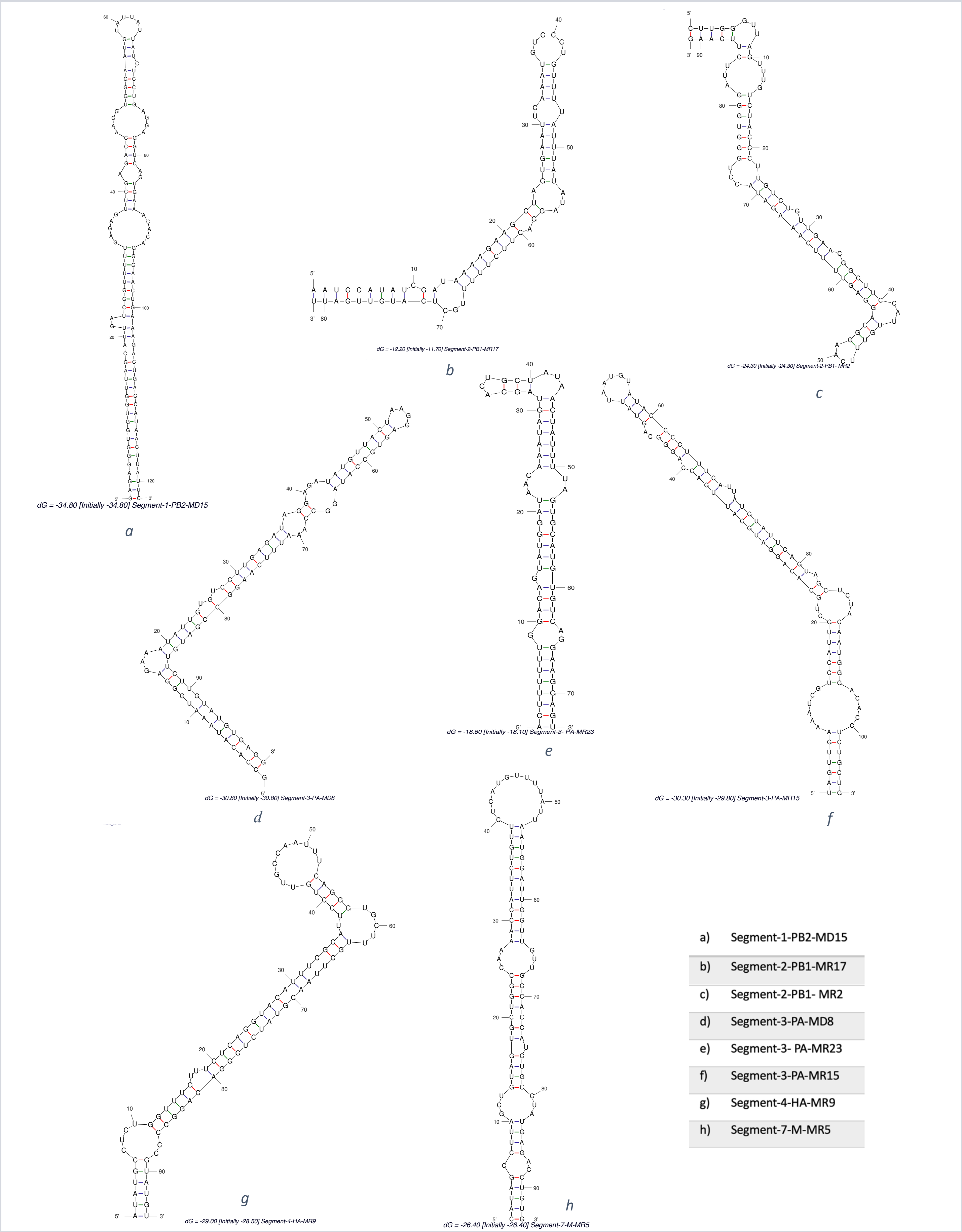


Supplementary Figure S1: pre miRNAs secondary structures predicted by Mfold.
